# Unraveling cfDNA Fragmentation Patterns: Linking Chromatin Architecture to Cancer Diagnostic Sensitivity

**DOI:** 10.1101/2024.11.16.623159

**Authors:** Andrew D Johnston, Fiach Antaw, Jennifer Lu, Clare Stirzaker, Matt Trau, Darren Korbie

## Abstract

Circulating cell-free DNA (cfDNA) liquid biopsies offer a powerful, non-invasive approach for cancer diagnostics, leveraging DNA fragments primarily released during apoptosis. Using droplet digital PCR on cfDNA samples from 88 breast and colorectal cancer patients, we uncovered highly consistent nucleosome breakpoints across individuals. Proximity to these cfDNA breakpoints diminishes the sensitivity of somatic mutation assays, including those targeting *IDH1* and *NRAS* hotspot mutations, underscoring the importance of precise nucleosome positioning data. We introduce a probabilistic nucleosome protection scoring method that enables high-resolution mapping of nucleosome positions and cfDNA breakpoints. These maps reveal distinct nucleosome spacing patterns between heterochromatin and euchromatin, corresponding to different chromatin fiber topologies. Analysis of X-chromosome inactivation further supports the topoisomer model: heterochromatin’s 187 bp nucleosome repeat length differs from euchromatin’s 182 bp, generating wave interference patterns in nucleosome signals. Our findings suggest that loop-mediated transitions from T2 to T1 topoisomers shape chromatin accessibility and cfDNA fragmentation, advancing the diagnostic potential of liquid biopsy assays.

**Highlights:**

- Nucleosome breakpoint conservation impacts somatic mutation assay sensitivity
- Probabilistic protection scoring enhances cfDNA nucleosome peak resolution
- Chromatin states exhibit unique nucleosome spacing linked to distinct topologies, influencing cfDNA fragmentation
- Chromosome X-inactivation phasing analyses reveal waveform interference in nucleosome positioning signals

**Graphical Abstract:** 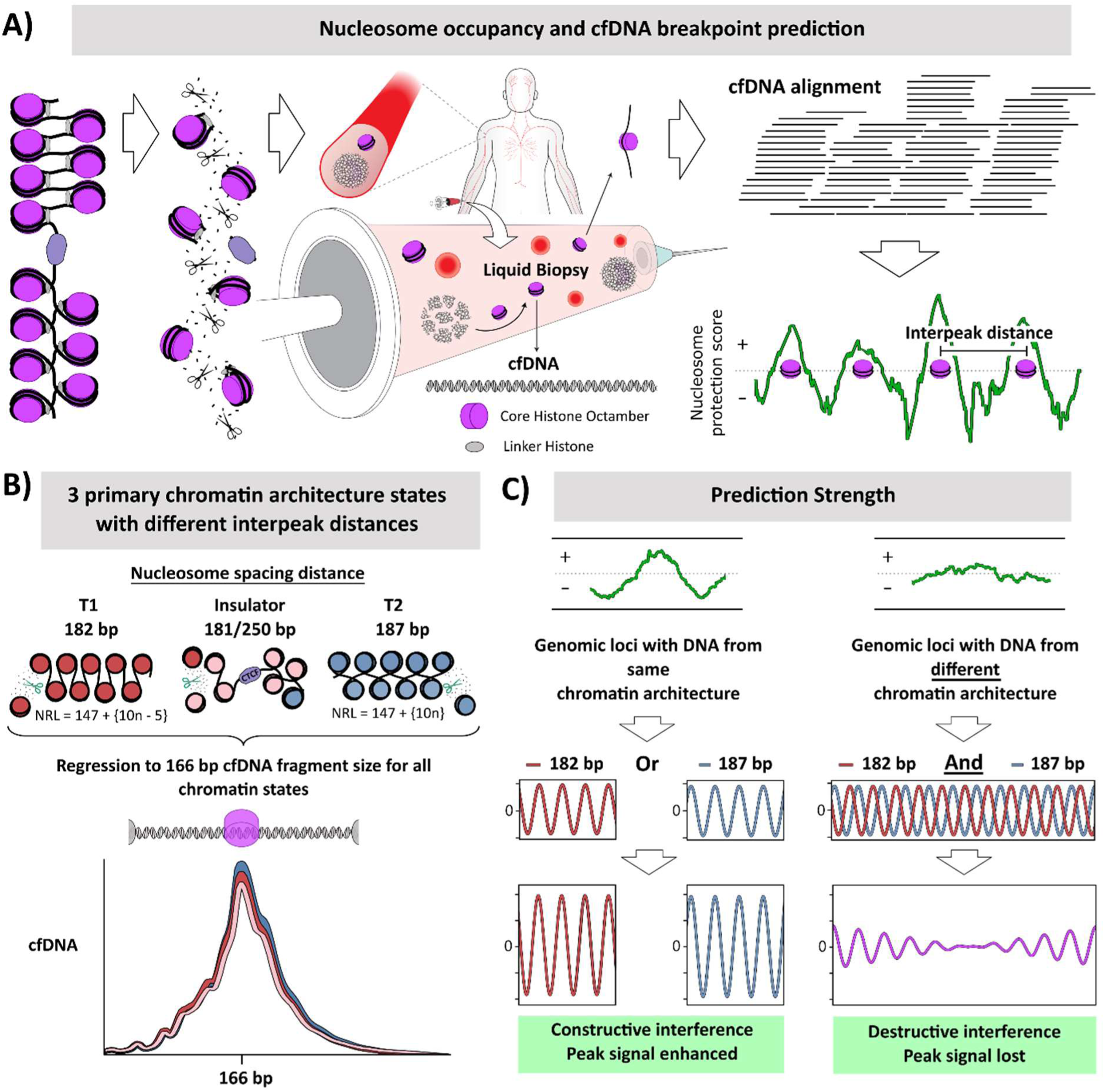

## Introduction

Circulating cell-free DNA (cfDNA) has emerged as a critical tool for non-invasive diagnostics, particularly in oncology, where liquid biopsies enable the detection of tumor-derived mutations and other genetic and epigenetic alterations without the need for invasive procedures^1,2^. Released primarily during apoptosis of hematopoietic cells, cfDNA is generated when nucleosomal DNA is protected from nuclease digestion, while the exposed linker DNA is cleaved^3^. This produces characteristic fragment length profiles that peak at 166 base pairs, corresponding to the length of DNA bound by histones^4^. cfDNA analysis has expanded beyond mutation detection to encompass the field of fragmentomics, where the study of cfDNA fragmentation patterns provides insights into chromatin architecture and tissue-specific gene regulation^5^.

In cfDNA samples with significant circulating tumor DNA (ctDNA) levels, the nucleosome positioning signal diverges from its typical hematopoietic pattern, revealing cancer-specific signals that can be used to detect tumor-associated changes in plasma^6^. These non-random fragmentation patterns reflect underlying epigenetic regulation and indicate variations in gene activity, especially near transcription start sites and regulatory regions. This approach highlights the unique gene activity of cancer cells compared to non-cancerous cells contributing to the cfDNA pool^7^.

A key challenge in liquid biopsy development is designing assays with optimal sensitivity for detecting mutations in ctDNA. Mutations in nucleosome-protected regions may be more readily detectable due to preservation during cfDNA biogenesis, while those in fragmented linker regions can be harder to identify^8^. Additionally, cfDNA contains fragments from multiple tissue types, with nucleosome positioning varying by cell type^9^, complicating cancer signal identification. Understanding these variations is essential for enhancing assay sensitivity, particularly in detecting early-stage and minimal residual disease and tracking tumor evolution.

While research has shown that nucleosome positioning is critical in shaping cfDNA fragmentation patterns, much is still unknown about how these patterns vary across different chromatin states, such as euchromatin and heterochromatin, due to shifts in nucleosome positioning between these states^10^. In particular, understanding how nucleosome positioning differs in tumor versus normal tissues due to altered chromatin states could enhance the diagnostic and prognostic value of cfDNA-based tests.

As the field of fragmentomics continues to evolve, there is a need for more refined methods to analyze how chromatin organization influences cfDNA fragmentation. In particular, understanding the conservation of nucleosome positioning across individuals and how this conservation interacts with cancer-associated changes in chromatin structure is an area of growing interest. Addressing these knowledge gaps could lead to significant advancements in liquid biopsy technologies, improving diagnostic sensitivity and enabling more precise monitoring of disease progression.

## Results and Discussion

To examine nucleosome protection and cfDNA breakpoint dynamics in a relevant model of human health and disease, cfDNA from n=88 individuals with cancer (n=54 breast cancer and n=34 colorectal cancer) was profiled using ddPCR assays targeting four nucleosome-protected peaks and their concomitant cfDNA breakpoints. Nucleosome peaks were those generated by Snyder et. al. (2016)^6^ using their windowed protection scoring method on cfDNA from over 50 healthy donors in the CH01 sample pool (**Table S1**, **Figure S1** and **Data S1**). To fine-map each peak, one ddPCR assay was centered on the predicted nucleosome binding site, where the maximum amount of intact cfDNA molecules should be present; and two additional flanking assays were centered where the most DNA breakage should occur, 80bp upstream and 80bp downstream of the nucleosome binding site (**Figure 1A**). Remarkably, comparison of amplifiable DNA copies at the nucleosome protection site versus the predicted upstream and downstream cfDNA breakpoints demonstrated extremely high statistical differences (**Fig1B**, p=<0.000001, one-tailed Mann-Whitney U test; **Table S2**) for every site selected for validation in both sets of cancer samples, with box plots showing no overlap for any of the regions analyzed (**Data S2**). To exclude the possibility of PCR amplification bias and confirm these results represented a real biological phenomenon of near-universal nucleosome positioning, two additional control experiments using HMW gDNA were undertaken on the hypotheses that (i) intact HMW gDNA should show no difference between the tiled ddPCR assays; and (ii) sonicated gDNA stochastically-broken and gel-purified to different fragment sizes (137, 162, 186, and 213 bp) should not exhibit concentration patterns similar to cfDNA in the tiled ddPCR assays, despite sharing similar fragment length profiles (**Fig1C**). Notably, both these gDNA controls failed to show the same statistical effects seen in the cfDNA cancer samples, thereby confirming the existence of nucleosome binding sites and corollary cfDNA breakpoints that exhibit near-universal positional conservation in the general population.

**Figure 1.**
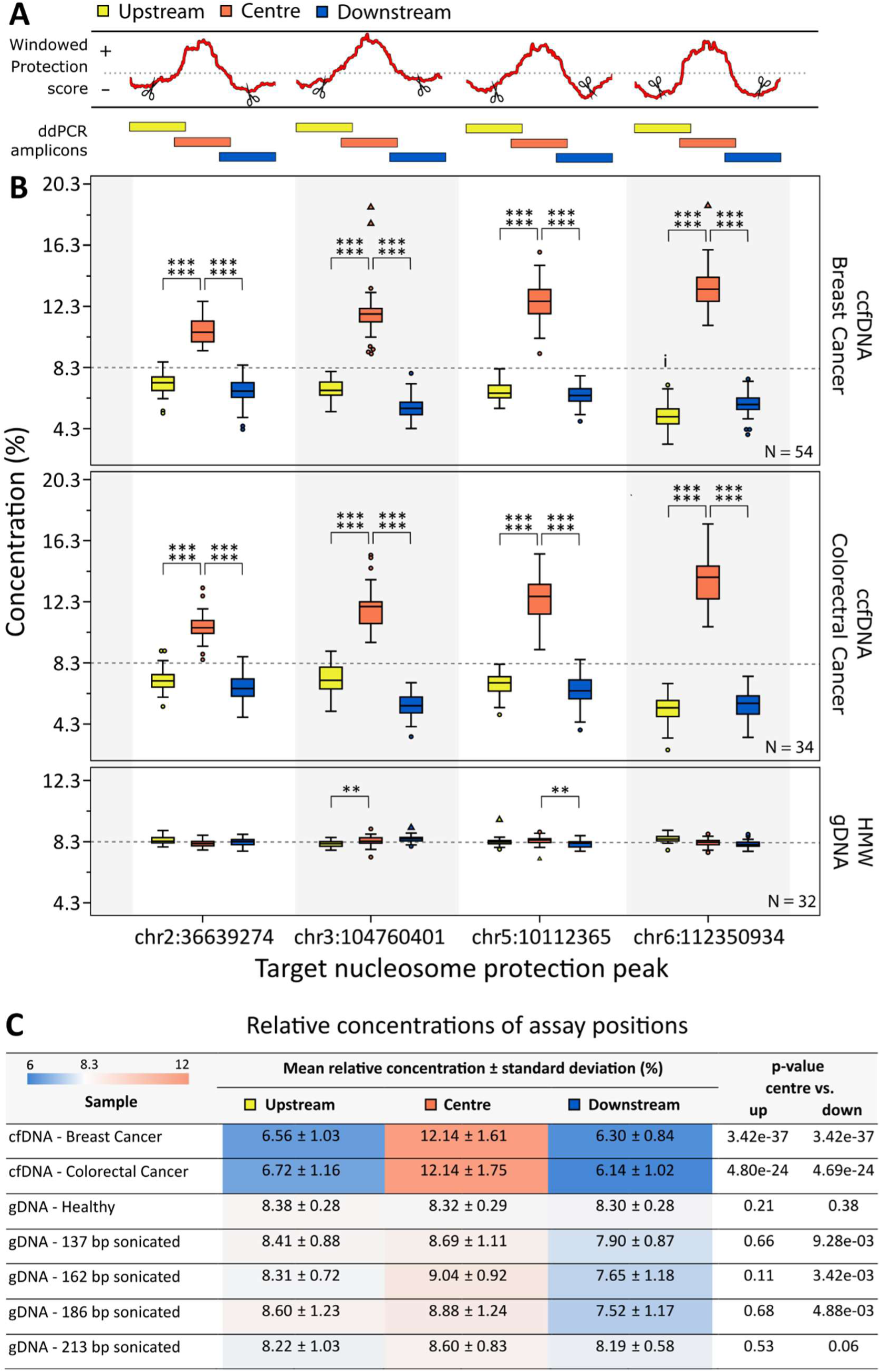
Fine-mapping of predicted cfDNA breakpoints identifies near-universally conserved nucleosomes in cfDNA samples. (**A**) Multiplexed ddPCR assays were designed to target a subset of genomic loci predicted to be most protected by nucleosome binding; together with proximal upstream and downstream ddPCR assays centered on the predicted cfDNA breakpoint site. (**B**) n=88 cfDNA samples from breast and colorectal cancer were analyzed using the multiplexed ddPCR assays. Plots show the DNA concentrations determined for each sample as a percentage across all 12 assays, which controls for differences in sample input amounts, with the expected concentration under the null hypothesis indicated at 8.3% (1:12). Outliers and extreme outliers are represented by circles and triangles, respectively. Statistical significance for each region within the samples was determined using one-sided Mann-Whitney U tests, with levels of significance denoted by *p < 0.05, **p < 0.01, ***p < 0.001, ****p < 0.0001, *****p < 0.00001, ******p < 0.000001. Control experiments with high molecular weight genomic DNA (HMW gDNA) stochastically fragmented to various sizes failed to reach the level of significance observed for patient cfDNA samples. (**C**) Table summarizing results for Wilcoxon signed-rank tests used to evaluate statistical significance of concentration differences among assay positions.

### PCR assays for somatic mutation analysis have reduced sensitivity near positionally-conserved nucleosome breakpoints

This positional conservation of nucleosome breakpoints has significant implications for diagnostic cfDNA assays, particularly in clinical settings where there is a growing interest in using cfDNA as a source of biopsy material. In cancer for example, if a known somatic mutation locus were to occur proximal to a positionally-conserved cfDNA breakpoint, PCR-based diagnostics targeting that mutation in a cfDNA sample could be severely compromised. For example, the chr6:112350934 nucleosome peak shows a nearly 3-fold difference in sensitivity between the nucleosome and breakpoint assays (**Fig1B**; 13.3% versus 5.2% relative concentration).

To assess the implications of cfDNA breakpoints on PCR-based somatic mutation assays for liquid biopsy analysis, the genomic coordinates of the 1104 most common somatic mutations^11,12^ were intersected with the CH01 nucleosome prediction map (Data S3). This analysis indicated that 448 (41%) of these canonical somatic mutations sit less than 30 bp from a predicted breakpoint; notably, this tenuous positioning could potentially afflict some of the most common mutation hotspots in cancer (Table S3). To explore this possibility, additional ddPCR multiplex assays were designed (Table S4) to assess the potential difference in diagnostic sensitivity when targeting four common somatic mutations hotspots located proximal to cfDNA breakpoints: *IDH1* (R132), *NRAS* (Q61), *KRAS* (G12), and *BRAF* (V600). To further test the efficacy of the windowed protection fragmentomic model, two of these mutations were selected near peaks with weak nucleosome protection scores (*KRAS* and *BRAF*), and two selected with relatively strong nucleosome positioning (*IDH1* and *NRAS*). ddPCR analysis of cfDNA isolated from breast cancer (n=16) and colorectal cancer (n=14) patients demonstrated that the two somatic mutations near the more prominent nucleosome peaks (*IDH1* and *NRAS*) exhibited a ∼1.5-fold loss in sensitivity (Figure S2, 14.9% versus 9.7% and 15.1% versus 10.9% relative concentration, respectively). For *BRAF* (V600), the hotspot assay concentration was significantly lower in the colorectal but not the breast cancer cohort, although the trend was the same in both. Like the previous experiment, control experiments using HMW and fragmented gDNA did not exhibit the same effects seen in the patient cfDNA samples. These results further validated the efficacy of using cfDNA-derived nucleosome protection to inform assay positioning, with significant implications for PCR-based profiling of cfDNA for somatic mutations.

### DNA length differences associated with nucleosome spacing and chromatin topology are lost during cfDNA biogenesis

To investigate the relationship between chromatin state and nucleosome occupancy, chromatin immunoprecipitation sequencing (ChIP-seq) data from ChromHMM analysis of the GM12878 B-lymphoblastoid cell line was analyzed^13^. A total of 13 unique chromatin states were used to annotate the CH01 map of nucleosome protection peaks. Notably, a comparison of heterochromatin, euchromatin, and chromatin bound by CTCF (i.e., “insulator” chromatin) revealed significant differences in nucleosome spacing among these three chromatin states (**Figure 2A**, p < 0.000001, Mann-Whitney U test; **Figure S3** and **Table S5**). Specifically, (i) heterochromatin showed a mode nucleosome spacing of 187 bp between adjacent nucleosomes, (ii) euchromatin had a mode spacing of 182 bp, and (iii) insulator chromatin bound by CTCF displayed a mode spacing of 181 bp, but with a narrower distribution and a secondary peak at 250 bp. The 187 bp spacing of heterochromatin corresponds to the 147 bp + {10n} nucleosome-repeat length characteristic of T2 chromatin fiber confirmations, as described by Norouzi et al. (2015)^14^, where 147 bp of DNA is wrapped around a nucleosome, plus 10 bp increments of linker DNA. Similarly, the 181-182 bp spacing of euchromatin and insulators aligns with the 147 bp + {10n – 5} repeat length, characteristic of the more flexible and accessible T1 topoisomers (**Figure 2B**). Notably, when cfDNA fragment size distributions for these ChromHMM-annotated chromatin states were plotted, all states showed an identical cfDNA fragment size distribution centered at 166 bp (**Figure 2C**).

**Figure 2.**
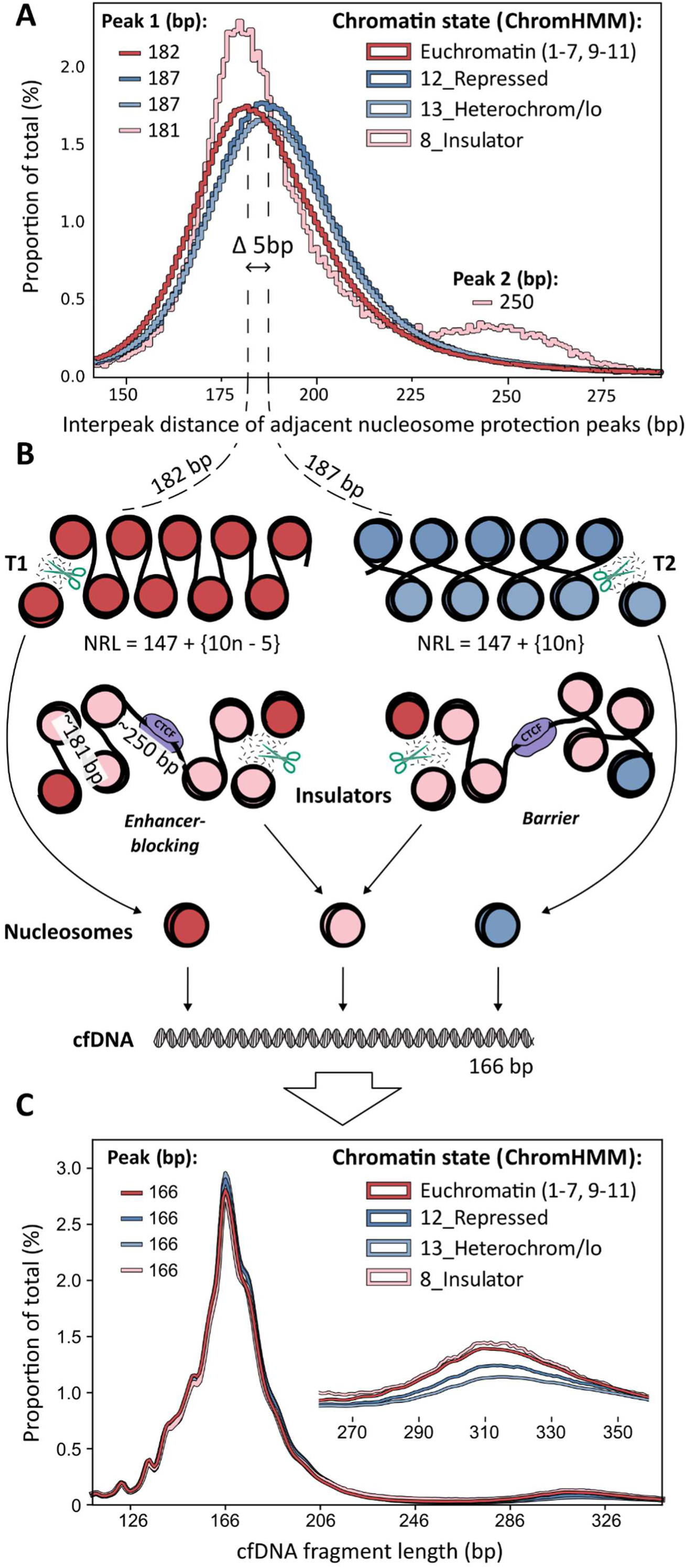
cfDNA fragments regress to a modal size of 166bp despite differences in chromatin state origin and nucleosome spacing. Chromatin states from ChromHMM were used to annotate our nucleosome occupancy map. (**A**) The distance between adjacent nucleosomes is different for euchromatin (red), heterochromatin (blue), and CTCF-bound insulator chromatin (pink). A notable 5 bp difference in nucleosome spacing between heterochromatin and euchromatin was observed, as denoted by the hatched vertical lines. (**B**) Euchromatin nucleosomes separated by 182bp correspond to T1 topoisomers, and heterochromatin nucleosomes separated by 187bp correspond to T2 topoisomers. CTCF-bound insulator regions are depicted a discrete chromatin state with a different topology and fragmentation profile, which separates euchromatin and heterochromatin. (**C**) Despite differences in the length of DNA between adjacent nucleosomes, distribution plots of cfDNA fragment size from euchromatin, heterochromatin, and CTCF-bound insulators all appear to regress to a modal size of 166bp, as determined by the length of sequenced cfDNA fragments.

To verify the previous observations relating to chromatin state and nucleosome spacing, a series of secondary analyses were next undertaken. First, it was hypothesized that if the chromatin state associations previously observed in **Figure 2** were a real phenomenon, then the relationship between chromatin topology and nucleosome spacing would be lost if ChromHMM chromatin state regions were randomly shuffled throughout each chromosome, resulting in highly similar nucleosome spacing lengths for all chromatin states analyzed. In line with the hypothesis, this shuffling resulted in all chromatin states exhibiting virtually identical nucleosome spacing (**Fig 3A-C**), confirming the original observation that the 181/182/187 bp differences were tied to specific genomic architectures.

**Figure 3.**
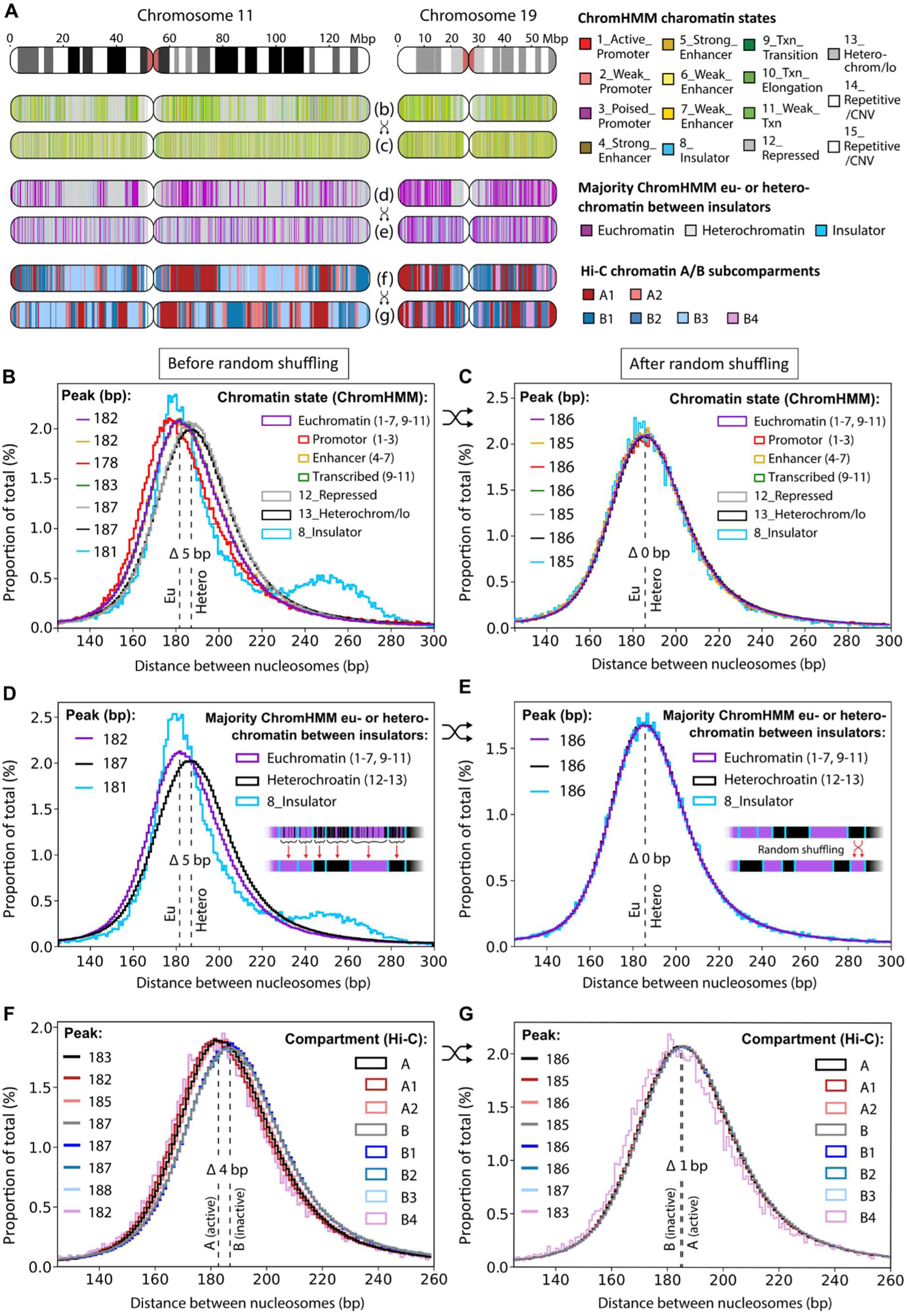
Nucleosome repeat lengths of chromatin categories before and after random shuffling. (**A**) Chromatin state categories on representative chromosomes 11 and 19 are shown before and after random shuffling. Observed differences in nucleosome spacing between heterochromatin and euchromatin are denoted by the hatched vertical lines on plots B-G. (**B**) Histograms showing the interpeak distances between nucleosome protection peaks (NPPs) aligned to chromatin states in the GM12878 B-lymphoblastoid cell line (based on ChromHMM analysis). (**C**) Interpeak distances after random shuffling of chromatin states shown in panel (B). (**D**) Chromatin states were redefined as either euchromatin or heterochromatin based on their predominant presence between two CTCF-bound insulators. (E) Interpeak distances after random shuffling of the redefined chromatin states in panel (D). (F) Histogram showing the interpeak distances between NPPs aligned to chromatin sub-compartments derived from Hi-C contact maps in the GM12878 B-lymphoblastoid cell line. (G) Interpeak distances after random shuffling of chromatin sub-compartments shown in panel (F).

Next, to explore how robustly CTCF-bound insulators demarcate overall genome architecture and topology, the predominant form of chromatin between each pair of CTCF-bound insulators was determined (i.e., either heterochromatin or euchromatin), and each CTCF-bounded genomic block was redefined as being wholly composed of either heterochromatin or euchromatin (**Figure 3D, inset illustration**). Notably, this simplified chromatin state annotation preserved the 182/187 bp nucleosome spacings previously observed and supports the hypothesis that chromatin state transitions are both demarcated and constrained by CTCF insulator barriers^15^. Similarly, random shuffling this simplified chromatin annotation resulted in a loss of nucleosome spacing differences (**Fig 3E**). Finally, the previous analyses were repeated using a second, independent chromatin annotation dataset based on the Hi-C chromatin contact A/B compartments and sub-compartments^16^. In line with our previous analyses the active/euchromatin compartment (denoted compartment A in this dataset) exhibited 183 bp spacing; and the inactive/heterochromatin (denoted compartment B) exhibited 187 bp spacing, with all states again exhibiting identical cfDNA fragment sizes centered on 166 bp (**Figure 3F**, **Figure S4** & **S5**). Similarly, the 183/187 bp spacing difference and disparities observed among sub-compartments were nullified when A/B chromatin compartments annotations underwent random shuffling (**Fig 3G**).

### An Improved Informatics Method for Fragmentomic Analysis and Nucleosome Peak-Calling

The analysis of cfDNA fragment patterns and nucleosome positioning demands precise computational methods, but our findings highlight limitations in the windowed protection scoring (WPS) approach of Snyder et al. (2016)^6^. Data from Figures 2 and 3 show the need to better distinguish subtle differences in nucleosome spacing, particularly the 5 bp variation between euchromatin (182 bp) and heterochromatin (187 bp). While the WPS method is a powerful and innovative approach for calling nucleosome positioning from cfDNA—and one we have employed in our analyses thus far—we have observed a few key limitations. Specifically, the method’s high signal-to-noise ratio and inability to perfectly normalize to a 0 baseline hinder its ability to detect weaker peaks and contribute to its reliance on considerable sequencing depths.

In response to these limitations, we developed a new informatic **Nucleosome Protection Scoring (NPS)** strategy that examines where cfDNA fragments are cleaved in relation to its proximal nucleosome-protection site. This new fragmentomic model begins with the observation that nucleosomes typically protect 166 bp of DNA and hypothesizes that (i) a probability distribution score weighted to consider both ends of each cfDNA fragment and (ii) which adjusts for fragment length variability could (iii) effectively predict nucleosome positions and cfDNA breakpoints across the genome. For our new method, as illustrated in **Figure 4**, two mirrored probability distributions are created for each cfDNA fragment: the first distribution starts at the 5’ fragment end, and the second initializes at the 3’ end. Next, a linear function applies a probability score to each base within a 166 bp window for both 5’ and 3’ prediction distributions as follows: at the cleavage site (i.e., the terminal 5’ or 3’ base in the DNA fragment) the score is weighted to 0 for the terminal base, then increases linearly base-to-base as the function moves inwards, until a maximum value of +1 is reached 83 bp from the fragment end. After reaching the maximum value of +1, the probability score decreases linearly for 83 bp and returns to 0 when 166 bp from the initial fragment end is reached. For fragments larger than 166 bp a 0 score is applied to all remaining bases; conversely, fragments smaller than 166 bp have probability distributions that extend beyond their length (**Figure 4A**). Next, the 5’ and 3’ fragment distributions are summed to generate a combined scoring distribution that is then centered around a mean of 0, resulting in minimum and maximum possible scores of -1 and +1, respectively (**Figure 4B**). For fragments exactly 166 bp in length the 5’ and 3’ probability distributions perfectly overlap, yielding a triangular distribution with a maximal combined score of +1 at the 2 bp cfDNA fragment center – indicating the base positions with the highest probability of nucleosome protection. As fragment lengths deviate from 166 bp, the overlap between the score distributions for each half of the fragment shifts from a peak to a plateau – a broadening of distribution that represents increased uncertainty in nucleosome prediction for that fragment. Of note, this new fragmentomic pipeline implicitly results in data normalization, as the sum of all prediction scores equals 0 for every DNA fragment. This internal normalization enhances peak detection, and contrasts with WPS, which normalizes to the median score within 1000 bp. As well, while the new NPS probabilistic scoring method is based on a cfDNA fragment size of 166 bp as seen in **Figure 2C**, fragment size can be easily adjusted for other contexts that generate different fragment populations (e.g., micrococcal nuclease digestions, ATAC-Seq, CUT&RUN; **Figure S6**).

**Figure 4.**
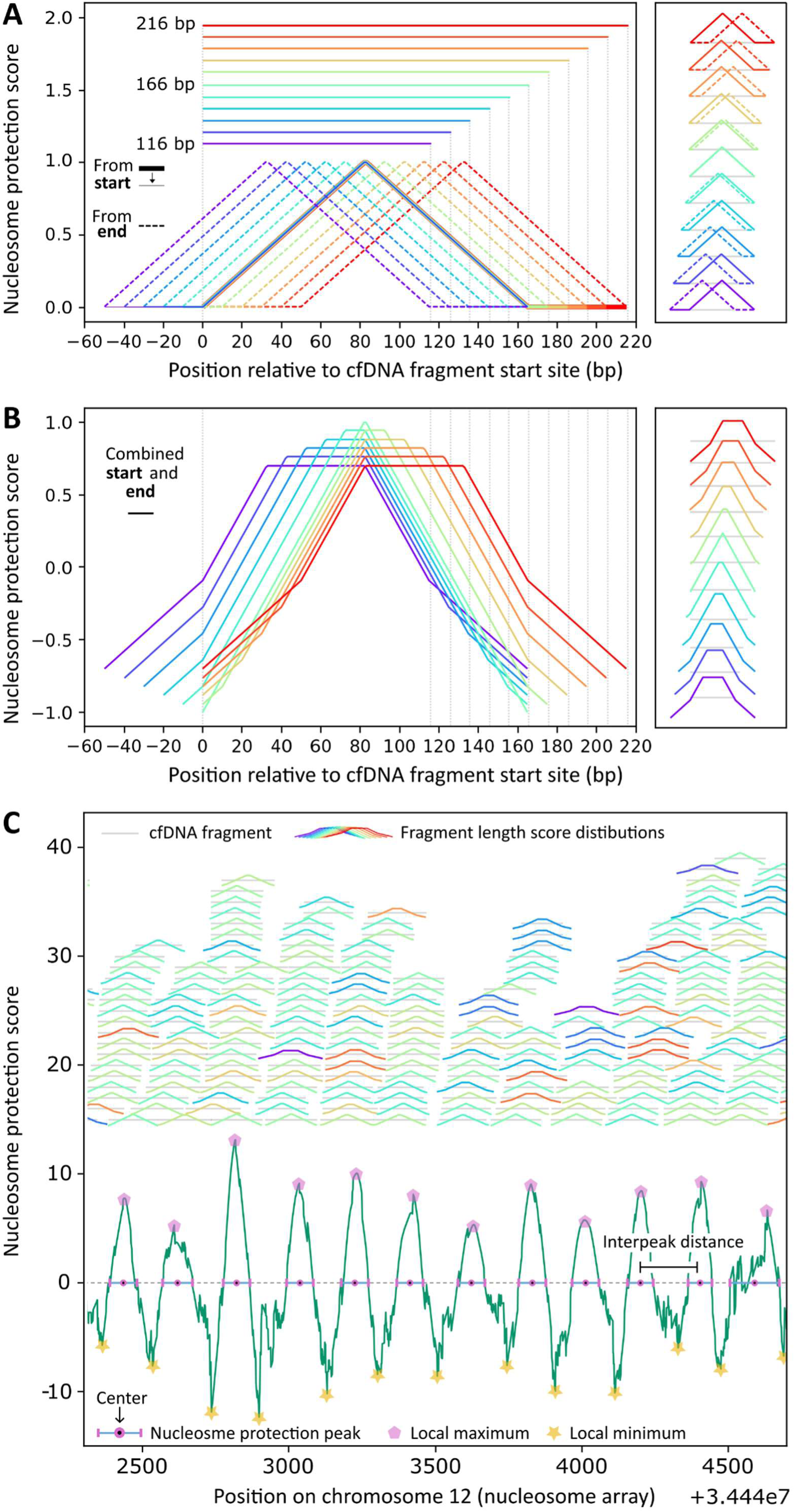
A new informatic method for fragmentomic analysis and nucleosome occupancy prediction. (**A**) Our model computes a probability distribution score weighted to consider both ends of each cfDNA fragment and which adjusts for fragment length variability. When dealing with multiple fragment lengths, including those shorter and longer than the canonical 166 bp cfDNA fragment, the scoring approach increases the protection score towards the center of each fragment. Fragments that align with the ideal 166 bp length show optimal overlap in their protection score distributions, which reflects the model’s assumption. As fragment lengths deviate from 166 bp, the protection score distributions become broader and less precise, reflecting greater uncertainty in nucleosome position prediction. (**B**) The resulting scoring distribution for different cfDNA fragment lengths is shown, which results in distinct plateaus and peaks based on fragment length size. A plateau indicates regions where multiple bases share the same nucleosome protection scores. In contrast, when a fragment length corresponds to the quintessential cfDNA fragment size it results in a maximal nucleosome position score. This enhances signal to noise ratio when estimating nucleosome placement and provide high-confidence predictions for nucleosome positioning. (**C**) The informatic method was applied to a region of chromosome 12 with highly conserved nucleosome positioning. To illustrate the method’s function, scoring was applied to a single cfDNA sample sequenced at low depth, allowing each individual fragment to be visualized. Individual cfDNA fragments, represented by colored curves, contribute to the overall nucleosome protection score, represented by the green trace in the lower portion of the panel. The nucleosome protection peak centers correspond to the predicted nucleosome dyad from which repeat length is determined, while local minima represent the genomic regions most susceptible to cleavage.

To demonstrate the benefits of our new NPS method, we compared regions surrounding the eight sites fine-mapped by ddPCR assays in Figures 1 and S2 against results from the WPS method. When we aggregated combined distributions from overlapping cfDNA fragments to infer nucleosome occupancy, NPS significantly enhanced the signal-to-noise ratio, producing more distinct nucleosome protection peaks (**Figure S7**). These peaks represent genomic positions most protected from nuclease cleavage, and therefore hypothesized as mid-points for nucleosome binding (**Figure 4C**). Additionally, regions with the lowest scores, corresponding to cfDNA breakpoints—areas where DNA cleavage is most frequent—were also more clearly defined.

Although there is substantial agreement between the two methods, the NPS approach produces smoother and more balanced profiles, resulting in better-resolved predictions for both cfDNA breakpoints and nucleosome occupancy peaks. Notably, the distribution of genome-wide spacing between WPS nucleosome peaks shows increments of 2-3 nucleosomes, suggesting missed nucleosome positions. These spacing increments do not appear in the NPS nucleosome peak distributions (**Figure S8**).

### Waveform properties of cfDNA nucleosome peaks are subject to constructive and destructive interference patterns

Aggregate nucleosome signals at transcription factor binding sites, especially CTCF-bound regions and transcription start sites, are used to study nucleosome positioning changes and their correlation with gene activity^7,17^. Previous studies have reported nucleosome phasing signals flanking these genomic features but without comparing them to baseline controls of average nucleosome phasing or positioning strengths. Despite this, these phasing signals are commonly interpreted as evidence of well-positioned nucleosome arrays^18,19^.

In our analyses, we found that these phasing signals are generally weaker than genome-wide nucleosome alignment signals, with the notable exception of CTCF (**Figure 5A-D**). This highlights a key point: if phasing signals indicate well-positioned nucleosomes near transcription factor binding sites, then similar signals when aligning to randomly sampled nucleosome positions across the genome would imply organized arrays everywhere. Otherwise, while the presence of phasing signals at active regulatory sites suggests a correlation with nucleosome positioning, this correlation does not imply consistently well-positioned arrays or strong nucleosome alignment.

**Figure 5.**
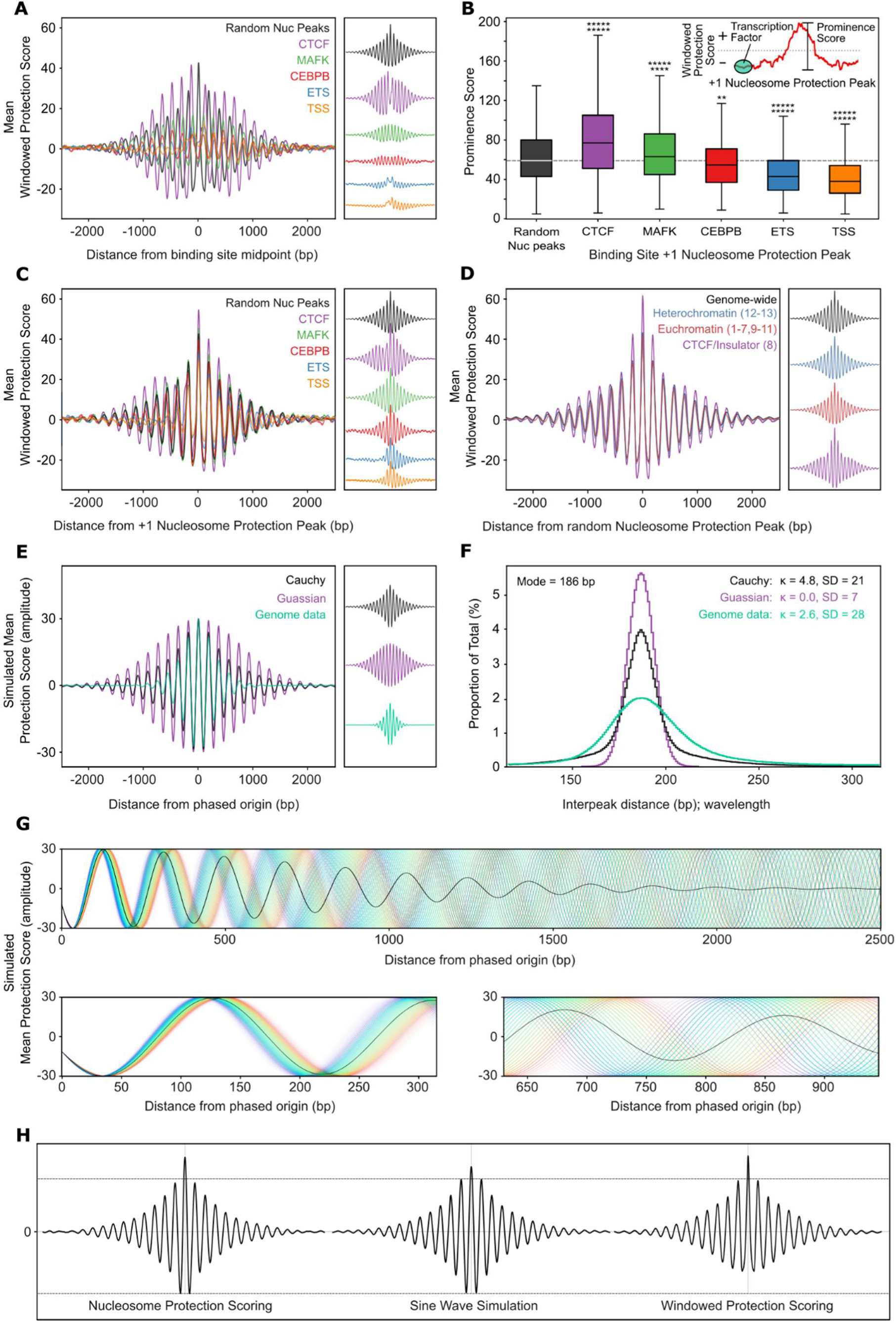
Nucleosome phasing and protection strengths relative to distance from genomic features. (**A**) Average windowed protection score (WPS) plotted as a function of distance from transcription factor binding sites and transcription start sites (TSS). (**B**) Boxplots of nucleosome protection peak (NPP) strength measured downstream of TF binding sites. Statistical significance of NPPs near these genomic features is assessed using two-sided Mann-Whitney U tests, comparing NPP strength against a set of 10,000 randomly selected NPPs. Significant differences are denoted by p < 0.05 (*), p < 0.01 (**), with additional stars representing decreasing p-value orders of magnitude. (**C**) Mean WPS plotted against distance from NPPs located directly downstream of TF binding sites and TSSs. (**D**) Mean WPS relative to distance from 100,000 randomly selected NPPs across each chromatin category (euchromatin, heterochromatin, and insulator chromatin). (**E**) Simulation of aggregate nucleosome protection signals using 1 million cosine waves with varying interpeak distances. The resulting signal profiles are shown for a Cauchy distribution of distances (representing WPS aligned to a random subset of NPPs) and a Gaussian distribution (representing WPS aligned to CTCF binding sites). This simulation recapitulates the observed wave interference patterns in aggregate WPS data. (**F**) Histogram showing the distributions of interpeak distances (wavelengths) used in wave simulations. These distributions reflect the proportions of interpeak distances that contribute to the averaged signals in panel E. (**G**) Sinusoidal wave phasing simulation of protection scores aligned to CTCF binding sites using the Gaussian distribution of interpeak distances in panel F. The colored lines represent individual sinusoidal waves with interpeak distances ranging from 151 bp (violet) to 218 bp (red). The black line represents the averaged summation of all sinusoidal waves, simulating the phased nucleosome protection signal near CTCF binding sites. (**H**) Comparison of aggregate scoring waves to aggregate Cauchy sine wave simulation from panel E. Windowed protection and nucleosome protection scores were aligned to a random subset of 100,000 nucleosome peaks called using their respective methods. Nucleosome protection scoring better matches the sinusoidal waveform.

To identify well-positioned nucleosomes, it is essential to evaluate the strength of aggregate nucleosome signals. Our analysis shows that both the phasing signals of averaged nucleosome scores at CTCF-bound sites and the positioning scores of their +1 nucleosomes are significantly stronger than the genome-wide average, determined by averaging scores aligned to a random subset of nucleosome peaks (p < 0.000001, Mann-Whitney U test). In contrast, signals at the transcription start sites of active genes are notably weaker than this genome-wide average (p < 0.000001, Mann-Whitney U test). Other examined sites, including those with elevated +1 nucleosome scores (though none as high as CTCF), showed weaker phasing signals than the nucleosome-aligned average. These results suggest that CTCF-bound loop anchors exert a unique organizing influence.

Sine wave-based simulations further suggest that these phasing signals arise from wave interference effects (**Figure 5E-F**). When simulating aggregated genomic regions with varying nucleosome repeat lengths, we observed constructive and destructive interference patterns that align with the phasing patterns observed in aggregate nucleosome scoring alignments. These findings highlight the role of nucleosome spacing variability in shaping aggregate signals.

This variability, both within and between chromatin states, contributes to wave interference that disrupts uniform signal alignment during aggregation. However, at CTCF-bound sites, nucleosome scoring is maximized, suggesting these sites serve as the phased origin of nucleosome spacing where wave interference is maximally constructive (**Figure 5G**; **Figure S9**). Moving away from these origins, the interference becomes increasingly destructive, leading to a progressive decline in aggregate signal strength.

### X-Inactivation as a Model for Chromatin State-Dependent Nucleosome Spacing

Compared to sine wave simulations, the WPS method shows a positive skew in protection scores, while NPS closely mirrors the sinusoidal waveform (**Figure 5H**). Given this functional similarity of NPS to a sine wave, the difference in nucleosome repeat lengths between euchromatin (182 bp) and heterochromatin (187 bp) should, in theory, result in wavelength differences. To explore this hypothesis, we first performed waveform simulations of nucleosome protection scores based on our nucleosome occupancy model. As shown in **Figure 6A**, although the two nucleosome peak waves for heterochromatin and euchromatin are initially in phase at the start of the simulation, the 5 bp offset ultimately causes nucleosome peak-scoring patterns to separate.

**Figure 6.**
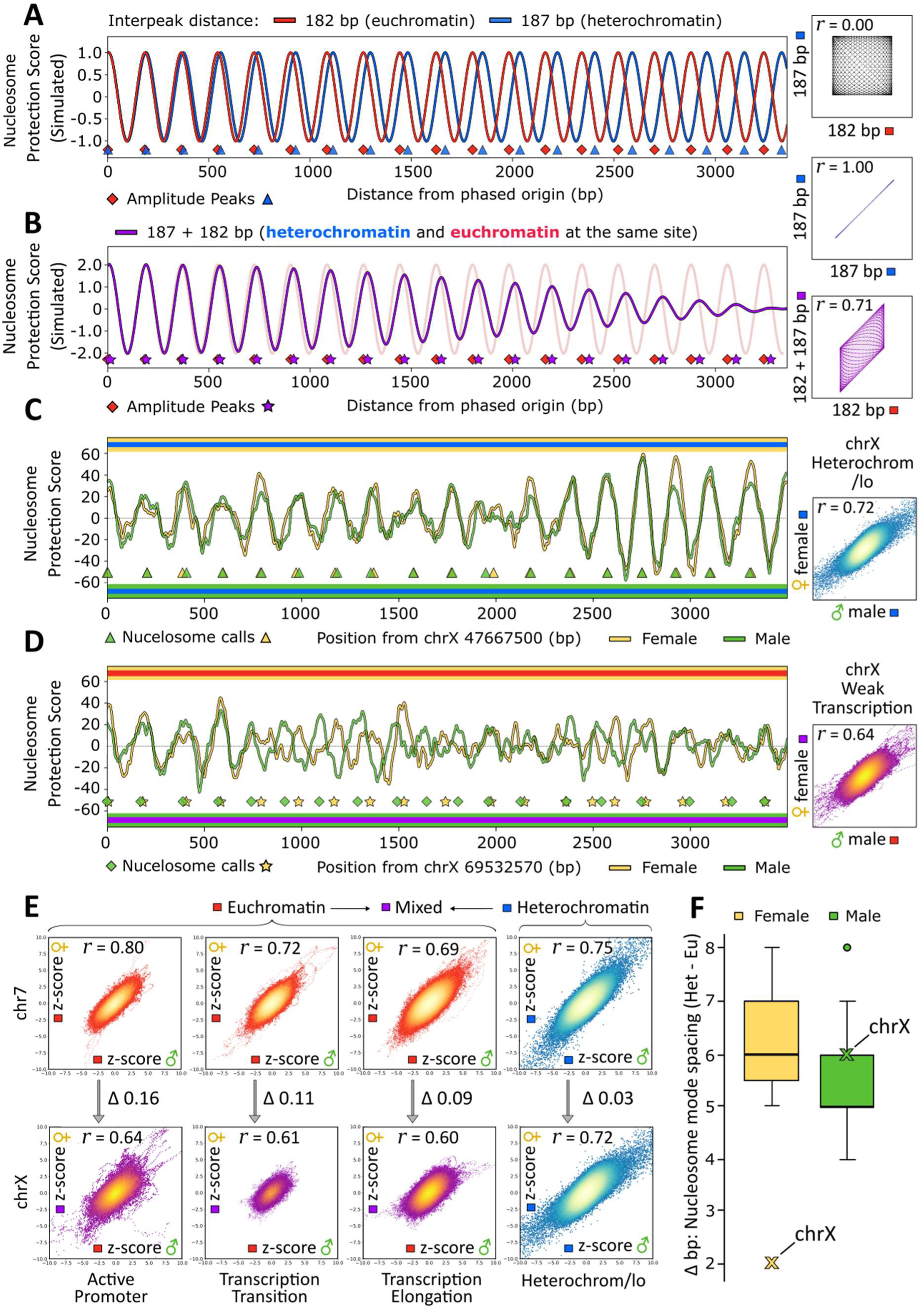
Waveform properties of cfDNA nucleosome peaks are subject to constructive and destructive interference patterns. (**A**) Sinusoidal wave simulations where the spacing between nucleosomes was modelled to be either 182bp (euchromatin, in red) or 187bp (heterochromatin, in blue). Red diamonds and blue triangles underneath indicate the peak height position, i.e., the center point where a nucleosome would be expected to bind. (**B**) When *in silico* nucleosome prediction sine waves for euchromatin and heterochromatin are superimposed and combined—as would be hypothesized for cfDNA samples derived from mixed cell types–constructive and destructive interreference occurs, leading to a beat-frequency signal which amplifies or ablates nucleosome occupancy predictions. (**C**) Nucleosome prediction scores were generated for a region of chrX annotated as heterochromatin, using cfDNA from male (yellow) and female (green) donors. When cfDNA fragments are derived from cell-types with the same chromatin state, nucleosome prediction scores exhibit a high-degree of correlation between different samples. In contrast, (**D**) when nucleosome prediction was performed on cfDNA fragments derived from a mixture of euchromatin and heterochromatin (female cfDNA, in green), resolution of nucleosome occupancy peaks was compromised and correlation with male cfDNA in a single chromatin state (yellow) was diminished. (**E**) Pearson correlation analysis of z-transformed nucleosome protection scores across chrX and autosomal control chr7, showing a reduced correlation between males and females in euchromatic regions of chrX. (**F**) Difference in mode nucleosome spacing between heterochromatin and euchromatin regions per chromosome, highlighting a reduction in chrX for female cfDNAs, as predicted by our model due to X-inactivation.

A key implication of the nucleosome waveform hypothesis is that cfDNA from different cell types (e.g., neutrophils, lymphocytes, and vascular endothelial cells), each contributing varying chromatin states (euchromatin and heterochromatin), would lead to constructive or destructive interference due to the overlapping cfDNA signals at specific intervals, altering the nucleosome peak-calling signal. To test this, we superimposed *in silico* nucleosome prediction sine waves for euchromatin and heterochromatin, mimicking the mixture of cfDNA from different cell types. This generated a beat-frequency signal, effectively amplifying or canceling nucleosome occupancy prediction due to the mixture of chromatin states in cfDNA (**Figure 6B**). According to the model, this interference occurs in ∼7 kb blocks: maximum destructive interference between the two waveforms occurs at 3366 bp, with re-alignment at 6732 bp. We hypothesize that unresolved nucleosome footprints at certain loci (e.g., **Figure S2B**) may result from analyzing mixed cfDNA fragments from different chromatin states.

While *in silico* modeling demonstrated the potential effects of the 5 bp nucleosome spacing offset on nucleosome position calling (**Figure S10**), quantifying this effect in actual cfDNA samples presented additional challenges. To accurately test the hypothesis, we needed to identify genomic regions where one homologous chromosome was heterochromatic and the other euchromatic. A suitable model system for this analysis is X-chromosome inactivation in females, where the inactive X chromosome is generally heterochromatic in the euchromatic regions of the active X homologue.

To test if the predicted 5 bp shift in nucleosome spacing between euchromatin and heterochromatin influences cfDNA fragmentation patterns, we compared nucleosome protection scores from male and female cfDNA samples. We hypothesized that nucleosome protection signals between males and females would align more closely in heterochromatin regions (**Figure 6C**) and progressively diverge in euchromatin regions on chrX (**Figure 6D**). Indeed, the correlations in nucleosome protection scores between male and female samples showed similarly strong alignment in heterochromatin on the autosomal control chr7 compared to chrX (Pearson’s r = 0.75 vs. 0.72, respectively; **Figure 6E**; **Figure S12**). Conversely, correlations on chrX in euchromatic regions were significantly weaker than on chr7, especially in poised promoters (r = 0.81 vs. 0.64, p < 0.000001), active promoters (r = 0.80 vs. 0.64, p < 0.000001), transcription transition (r = 0.72 vs. 0.61, p < 0.000001), and transcription elongation regions (r = 0.69 vs. 0.60, p < 0.000001). Similar trends were observed when comparing nucleosome spacing and agreement between matched nearest-neighbor peaks in male versus female samples (**Figure S12** and **Table S6**).

Finally, we calculated mode nucleosome spacing differences between euchromatin and heterochromatin across chromosomes in both male and female samples (**Figure 6F**; **Figure S13** and **Table S7**). As expected, the nucleosome spacing difference between euchromatin and heterochromatin on chrX in females was notably smaller (2 bp) than on chr7 (6 bp) or other autosomes (median = 6 bp). In males, the spacing difference on chrX was the same as autosomes (6 bp), consistent with the absence of X-inactivation. These findings support our hypothesis that nucleosome positioning on the X chromosome in females reflects a composite signal from heterochromatin and euchromatin, reinforcing the notion that nucleosome repeat length differs by 5-6 bp between these chromatin states and adjusts dynamically with chromatin state transitions.

## Conclusion

Our study provides new insights into chromatin biology and cfDNA-based diagnostics by linking nucleosome repeat lengths (NRLs) in cfDNA to distinct chromatin topoisomers. Specifically, we align the 182 bp NRL of euchromatin with the T1 topoisomer and the 187 bp NRL of heterochromatin with the T2 topoisomer, supporting the *in vivo* validity of the topoisomer model proposed by Norouzi et al. (2015)^14^. This model provides a valuable framework for understanding how chromatin organizes into distinct functional states.

Under the topoisomer model, NRL is essential for distinguishing euchromatin from heterochromatin. While population-wide analyses often reveal irregular nucleosome positioning due to cell-to-cell variability^19^, single-cell nanopore sequencing of MNase-digested fragments shows much more regular nucleosome spacing within individual cells^8^. This regularity at the single-cell level, contrasted with diffuse nucleosome signals in population-wide analyses, suggests that nucleosomes move in a coordinated manner, maintaining synchronized positioning across a region. We propose that this coordinated movement results from loop-mediated transitions between chromatin topologies, where T2 topoisomers (187 bp spacing) convert to T1 topoisomers (182 bp spacing), with the diffuse signals observed in population-wide analyses arising from wave interference patterns generated by shifts in chromatin topology. Unlike the classic “beads on a string” model, our “beads on a looped spring” model captures the dynamic and responsive nature of chromatin, emphasizing its ability to reorganize adaptively across different chromatin states (**Figure 7**).

**Figure 7.**
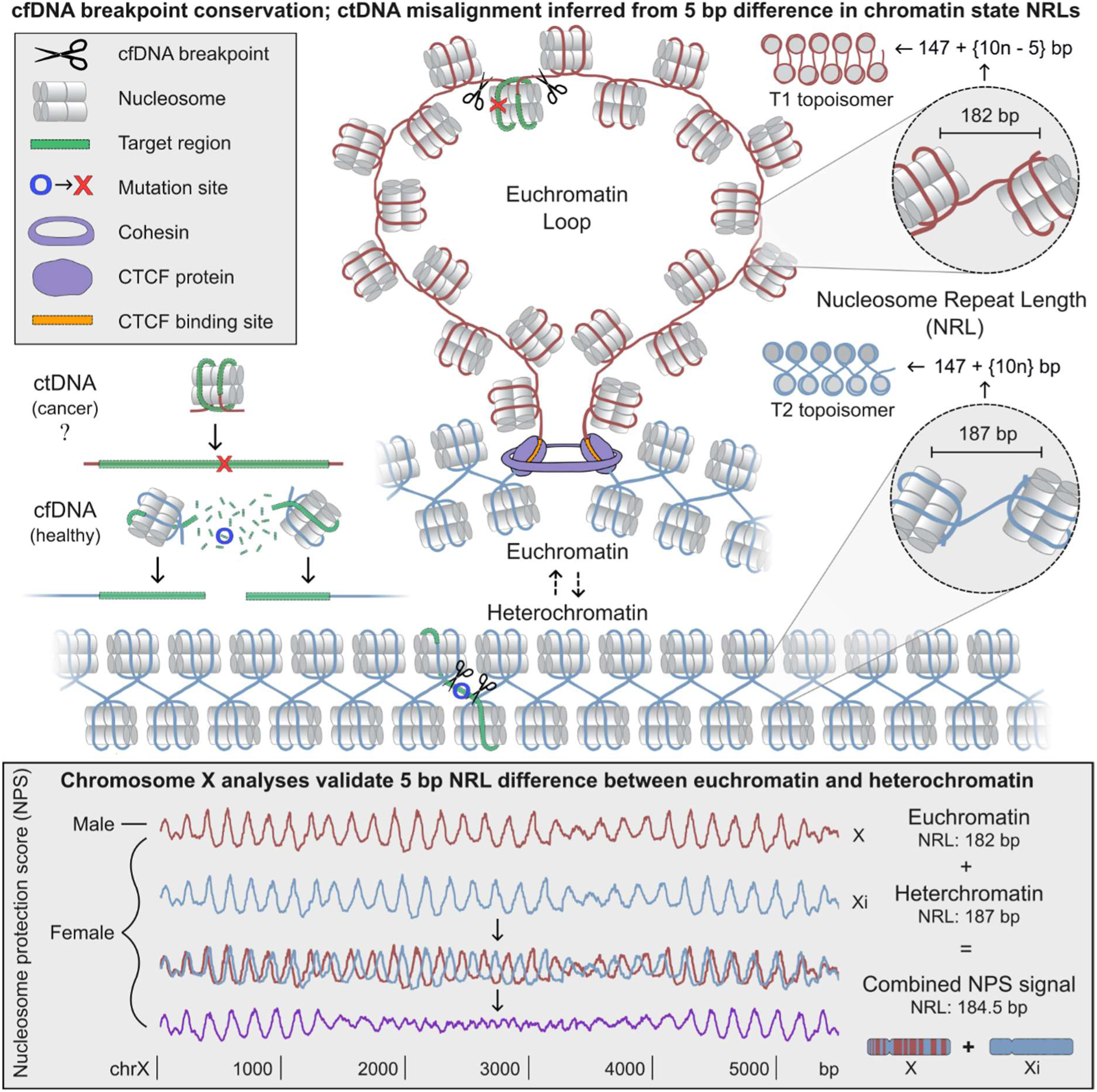
Model of how euchromatin forms due to shifts in nucleosome repeat length within chromatin loops. This model proposes that nucleosome repeat length (NRL) dynamics, driven by chromatin loop extrusion, are key to the formation of open chromatin. Euchromatin is characterized by a reduced NRL of 182 bp compared to the 187 bp NRL of heterochromatin. These distinct NRLs produce different chromatin topoisomers: T1 for euchromatin and T2 for heterochromatin, as demonstrated in the *in silico* and *in vitro* work of Norouzi et al. (2015)^14^. Our data analyses of *in vivo* nucleosome repeat lengths support this topoisomer model. Our “beads on a looped spring” model proposes that chromatin loops dynamically transition between these topoisomers, facilitating the shift from condensed heterochromatin to open euchromatin. Analysis of chrX in females reinforces this dynamic, showing that euchromatin’s T1 spacing shifts to T2 spacing within X-inactivated regions. For cfDNA liquid biopsy assays, understanding how these NRL shifts modulate chromatin accessibility and nucleosome protection is crucial for improving ctDNA detection. Targeting regions with misaligned nucleosome protection peaks that differ between tumor and non-tumor tissue types can enhance the precision of assays by focusing on the most cancer-relevant cfDNA fragments. When targeting somatic mutations near conserved cfDNA breakpoints, assay sensitivity can be improved by reducing amplicon size and positioning them as close as possible to the nearest nucleosome protection peak within the cancer tissue’s cell type to minimize the impact of conserved nucleosome breakpoints.

In this model, the NRL of a genomic region reflects the proportion of time it exists as looped T1 chromatin fibers. For example, regions with weak transcription exhibit an NRL of 183 bp, positioned between the 180-181 bp NRLs observed in transcription transition and elongation, and the 187 bp NRLs in Polycomb-repressed regions and heterochromatin. Similarly, the 179 bp NRL of weak promoters falls between the 177 bp NRL of strong promoters and the 182 bp NRL of poised promoters. This model suggests that these intermediate NRLs emerge from intermittent looping within cells or differential regulation across cells, leading to a toggling between open and closed chromatin configurations. This dynamic behavior is expected to produce oscillations in nucleosome positioning signals, resembling the patterns we observed in euchromatic regions of the female X chromosome.

Building on the established role of CTCF in chromatin organization, this study underscores CTCF binding sites as uniquely associated with strong nucleosome phasing and positioning signals. These robust signals suggest that chromatin loop anchors at these sites act as phased origins of nucleosome spacing. From this, we infer that CTCF-anchored chromatin loops play a crucial role in establishing and maintaining nucleosome spacing, with this spacing intrinsically linked to transitions between distinct chromatin topologies. We propose that CTCF/cohesin-mediated loop extrusion adjusts NRLs to facilitate transitions in chromatin topology.

In their 2023 study, Bomber et al.^20^ demonstrated that SMARCA5 co-localizes with CTCF and is essential for maintaining nucleosome spacing, as its degradation leads to increased nucleosome repeat length and disrupted nucleosomal phasing around CTCF binding sites. Building upon these findings, we propose that during CTCF/cohesin-mediated loop extrusion, SMARCA5 actively slides nucleosomes to establish a 147 + {10n – 5} bp spacing, facilitating the transition from T2 to T1 chromatin topology and anchoring nucleosome positioning within the looped structure. This association of T1 topology with chromatin loops also provides a possible model for how mitotic chromatin condenses: during mitosis, the entire chromosome transitions into a looped T1 state, dissolving phase separation between T1 (euchromatin) and T2 (heterochromatin) fibers present during interphase. This uniform looping aligns with the findings of Gibcus et al. (2018)^21^, who demonstrated that mitotic chromosomes are organized as chromatin loops radiating from a central scaffold, facilitating efficient chromatin compaction into dense, liquid crystal-like structures^22,23^. Notably, the stretched forms of T1 and T2 topoisomers correspond well with the 5 to 24 nm disordered chains observed within *in vivo* interphase and mitotic chromatin using ChromEMT^24,25^.

Recognizing how nucleosome positioning and chromatin topology affect cfDNA fragmentation enables the design of assays targeting regions where cfDNA remains intact, enhancing the detection of somatic mutations and other cancer-associated changes. The sensitivity of cfDNA-based assays is strongly influenced by nucleosome positioning, which directly impacts variant detection. Our findings indicate that assay sensitivity is significantly higher when somatic mutations are located within nucleosome-protected regions of cfDNA.

Additionally, mixed chromatin states impact nucleosome peak-calling. In regions where chromatin states differ among the cell types contributing to cfDNA, destructive interference between nucleosome signals can occur. Potential within-cell shifts in chromatin topology throughout the cell cycle may further contribute to this interference^26,27^, complicating accurate nucleosome position calling but not necessarily affecting mutation-detection sensitivity. Instead, sensitivity depends on the assay’s ability to target mutations relative to nucleosome protection within the cancer cell population, rather than on bulk cfDNA peak signals. This underscores the importance of developing an atlas of nucleosome positions and breakpoints across diverse cell types to enable more precise assay design. Such an atlas could be created using the enhanced nucleosome peak-calling method described here, applied to nuclease-digested chromatin from individual cell types, such as cfMNase-Seq^28^.

Beyond nucleosome positioning, cell-type-specific DNA methylation patterns provide a powerful means of identifying the tissue of origin in ctDNA^29^. To improve the sensitivity of cfDNA assays for detecting ctDNA, we propose targeting tissue-specific methylation within regions of strong but differential nucleosome protection. By focusing on genomic regions where the tissue of origin exhibits stable methylation patterns rarely disrupted by cancer^30,31^, we can develop more universal cancer detection assays that enrich for cfDNA originating from the tumor’s tissue. Hodges et al. (2011)^32^ demonstrated that DNA methylation changes often correlate with nucleosome positioning across hematopoietic lineages, suggesting that regions with tissue-specific differential methylation are also likely to display differential nucleosome positioning. This approach leverages the stability of methylation and nucleosome patterns unique to specific cell types, ensuring that even when cancer-associated changes occur, the underlying tissue-specific signals remain detectable.

For cancers arising from cell types that do not typically release cfDNA under healthy conditions, incorporating these stable markers into liquid biopsy assays can improve the ability to distinguish ctDNA from background cfDNA. For cell types that do contribute to healthy cfDNA, integrating the most common cancer-associated changes in DNA methylation and nucleosome positioning—including topological changes that result in nucleosome peaks that are out of phase between ctDNA and background cfDNA—could significantly enhance the assay’s accuracy and sensitivity.

Overall, our study advances the understanding of how chromatin architecture shapes cfDNA fragmentation, with direct applications for enhancing liquid biopsy diagnostics. By leveraging insights into nucleosome positioning and recognizing variations in chromatin topology, we lay the groundwork for improving diagnostic sensitivity, particularly for early-stage cancer detection and for monitoring minimal residual disease and recurrence, where gains in assay sensitivity are paramount.

## Supporting information

Additional Supplementary Materials - Unreferenced

## Acknowledgements

We thank Davood Norouzi for his valuable correspondence and Sasha Main for her insightful feedback on this manuscript.

## Author contributions

Conceptualization: AJ, DK

Data curation:

Formal analysis: AJ

Funding acquisition: MT, CS, DK

Investigation: AJ, DK

Methodology: AJ, FA

Project administration: AJ, DK

Resources: DK, MT

Software: AJ, FA, JL

Supervision: MT, DK

Validation: AJ, DK

Visualization: AJ

Writing – original draft: AJ

Writing – review & editing: AJ, DK, CS, MT, JL, FA

