## Additional Supplementary Materials - Unreferenced for "Unraveling cfDNA Fragmentation Patterns: Linking Chromatin Architecture to Cancer Diagnostic Sensitivity"

### Extended Supplementary Figures

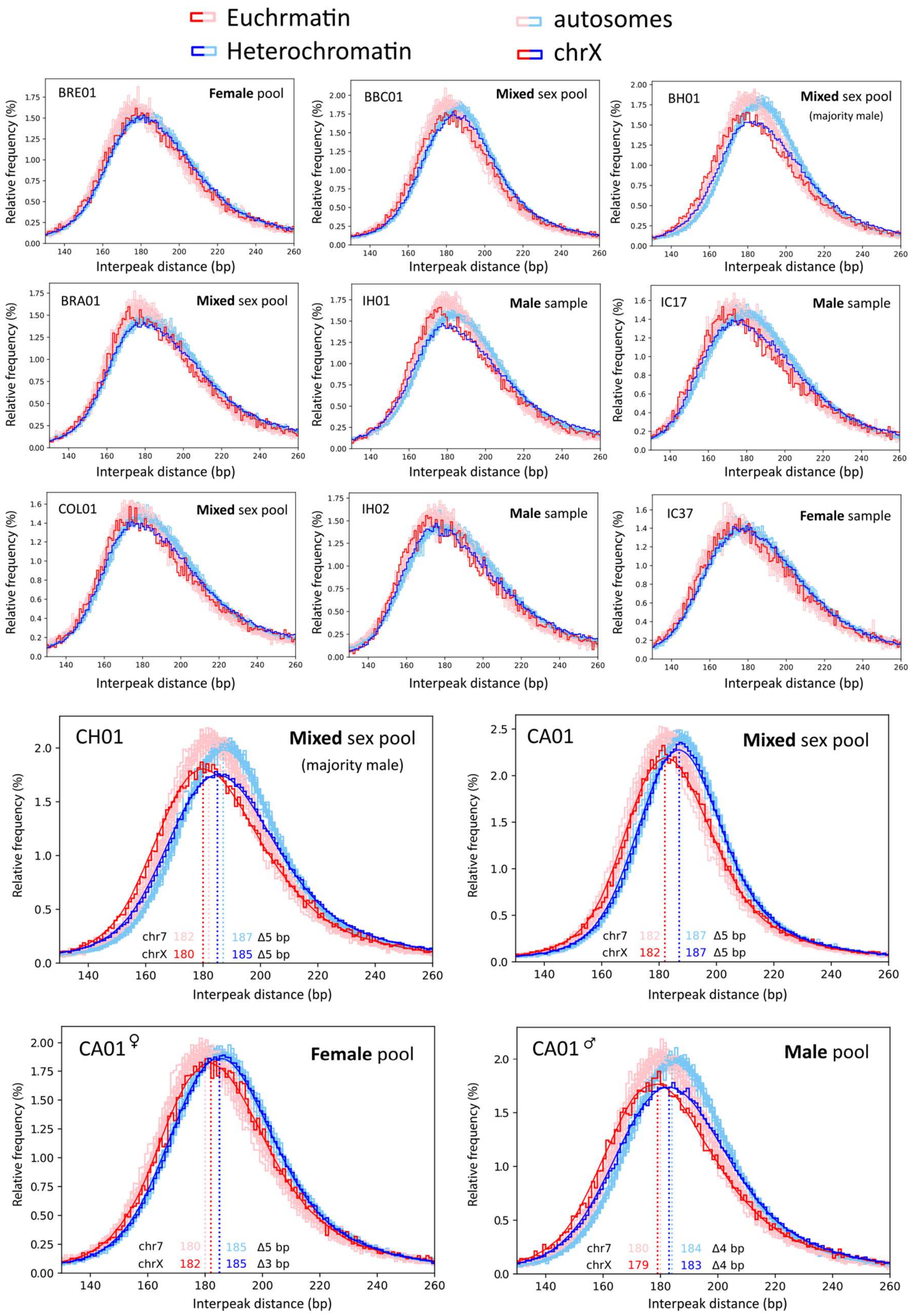

**Supplementary Figure S16. Histograms of the distance between adjacent nucleosome protection peaks (NPPs) genome wide.** Differences between Ernst et al. (2016)[2] euchromatin (states 1-7, 9-11) and heterochromatin (states 12-13) mode NPP spacing after Savitzky-Golay smoothing are displayed for chrX and chr7 for sample pools with high sequencing coverage. The sex of a patient for individual samples, or sexes contained within sample pools, are displayed. For the BH01, CH01 and CA01 sample pools, the sexes displayed are based on WPS differences between chrX and autosomes, as the exact percentage breakdown of sexes contained in these pools is unknown.

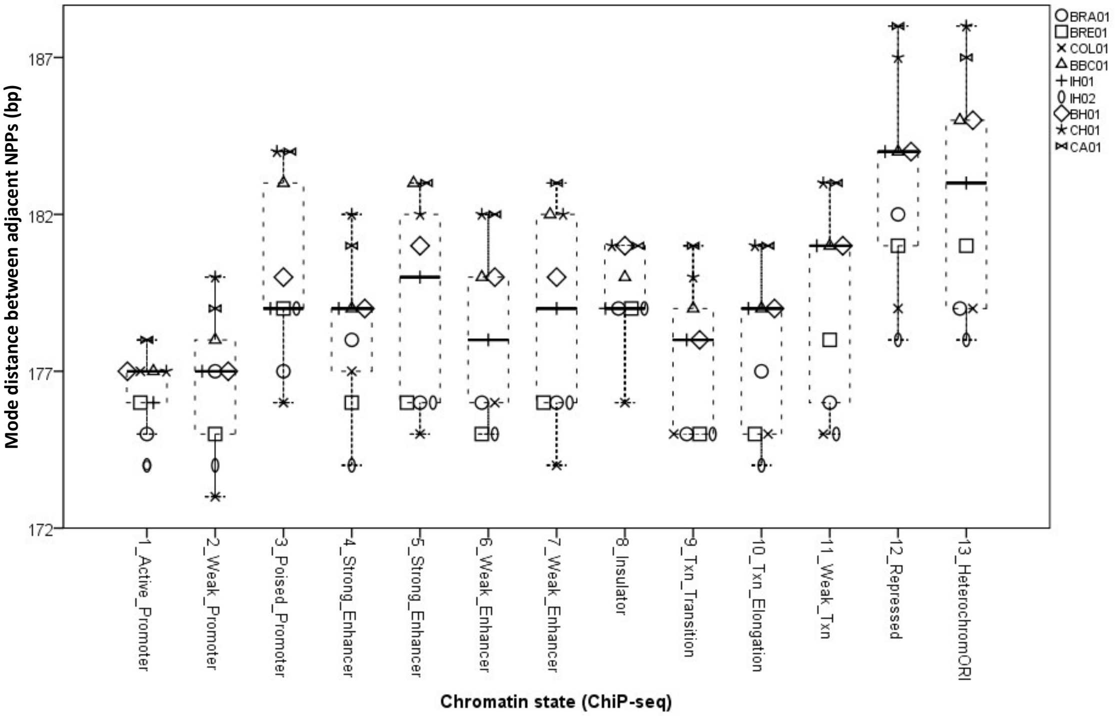


**Supplementary Figure S17. Mode interpeak distance of nucleosome protection peaks (NPPs) by sample and chromatin state after Savitzky-Golay smoothing.** The bottom line of each box represents the 25th percentile, top line the 75th percentile, and thick middle line the median. Whiskers extend up to a maximum of 1.5 times the height of the box. Any values that fall outside this range are classified as outliers.

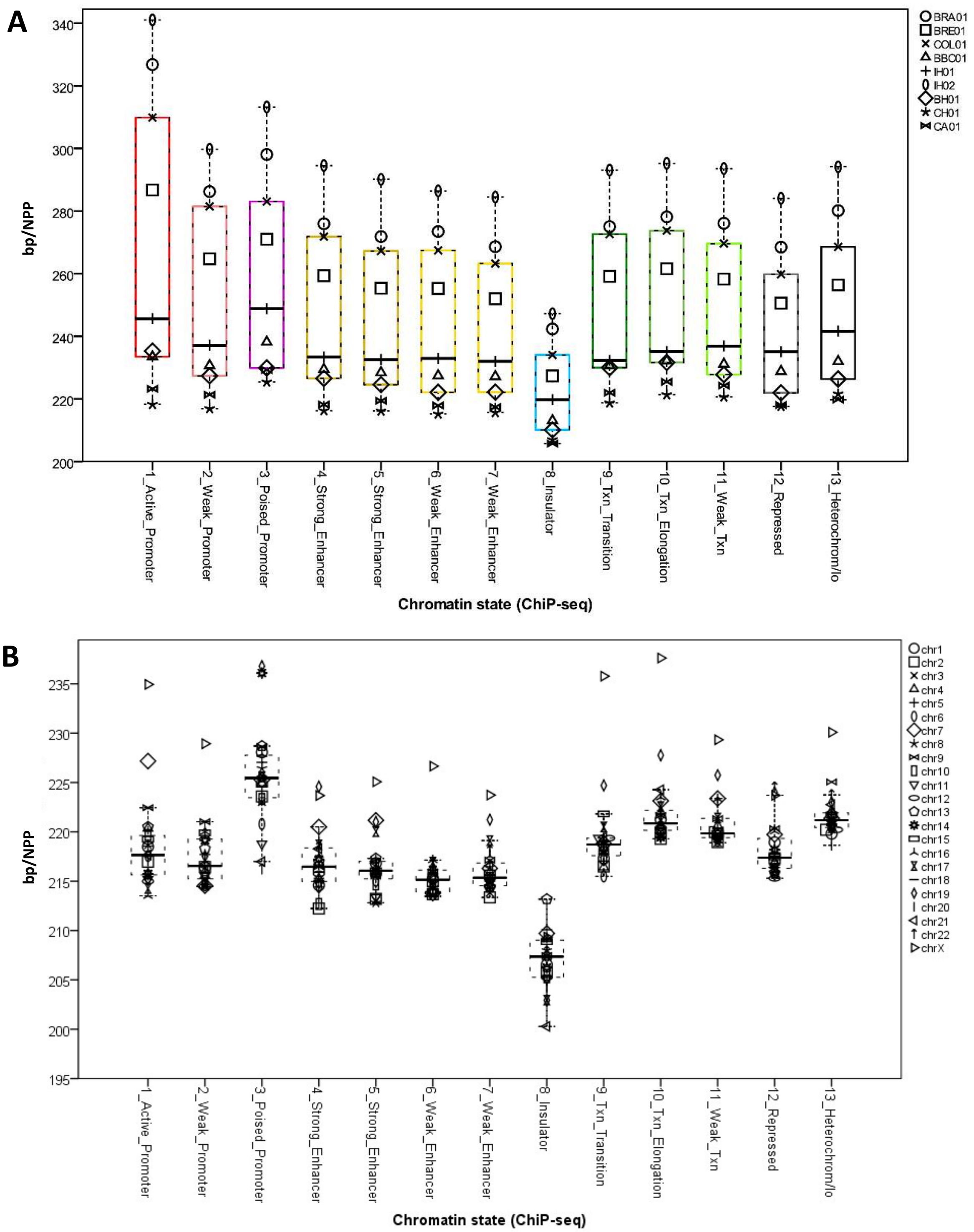


**Supplementary Figure S17. Chromatin state nucleosome densities.** (**A**) Nucleosome density (NPP/bp) of each chromatin state for each sample. (**B**) Nucleosome density for each chromatin state for each chromosome in CH01 sample pool. Lower base pairs (bp) per nucleosome protection peak (NPP) translates to higher nucleosome density. The bottom line of each box represents the 25th percentile, top line the 75th percentile, and thick middle line the median. Whiskers extend up to a maximum of 1.5 times the height of the box. Any values that fall outside this range are classified as outliers.

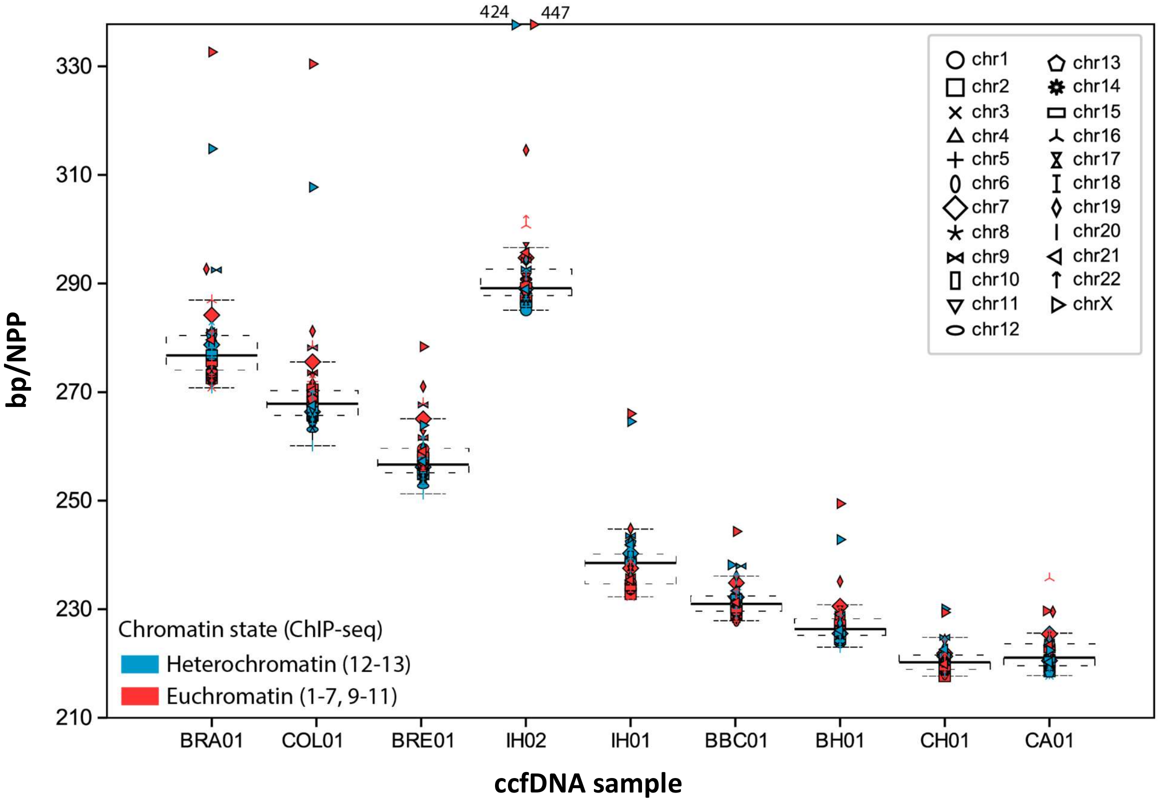


**Supplementary Figure S18. Nucleosome density by sample and chromosome.** Lower base pairs (bp) per nucleosome protection peak (NPP) translates to higher nucleosome density. The bottom line of each box represents the 25th percentile, top line the 75th percentile, and thick middle line the median. Whiskers extend up to a maximum of 1.5 times the height of the box. Any values that fall outside this range are classified as outliers.

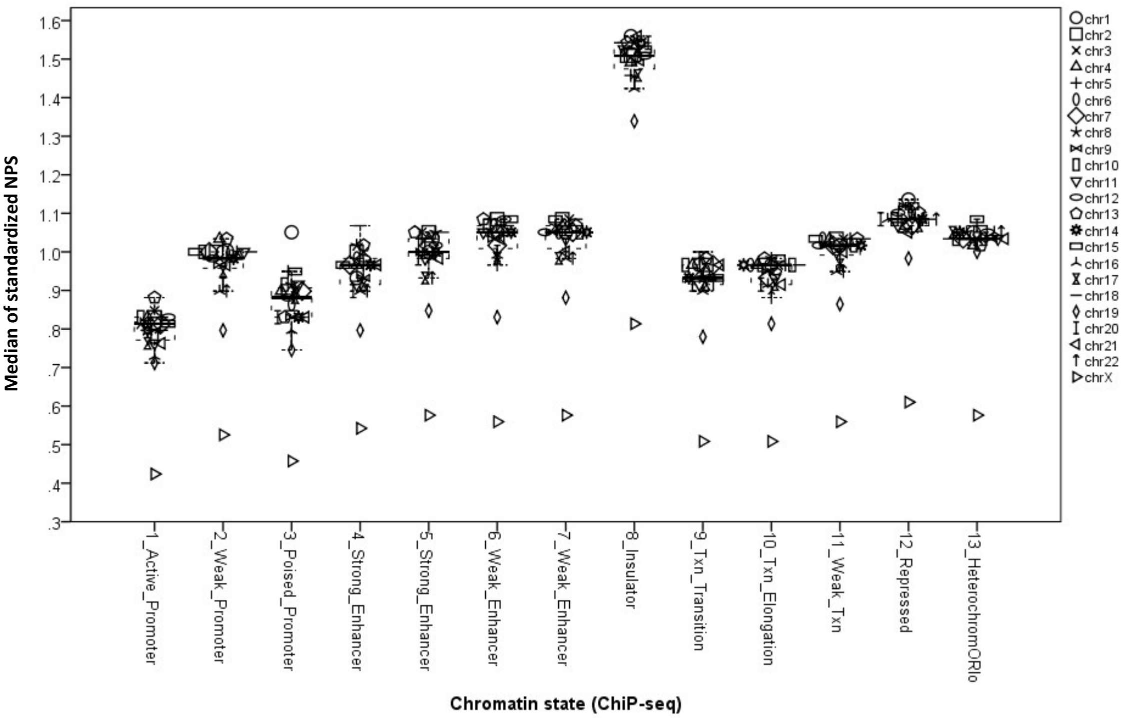


**Supplementary Figure S19. NPS medians by chromosome and chromatin state in CH01 sample pool.** To account for differences in sequencing depths between samples, standardized normalized prominence score (NPS) was calculated by dividing the NPS of a nucleosome protection peak (NNP) by the median NPS of all NPPs called for a sample. The bottom line of each box represents the 25th percentile, top line the 75th percentile, and thick middle line the median. Whiskers extend up to a maximum of 1.5 times the height of the box. Any values that fall outside this range are classified as outliers.

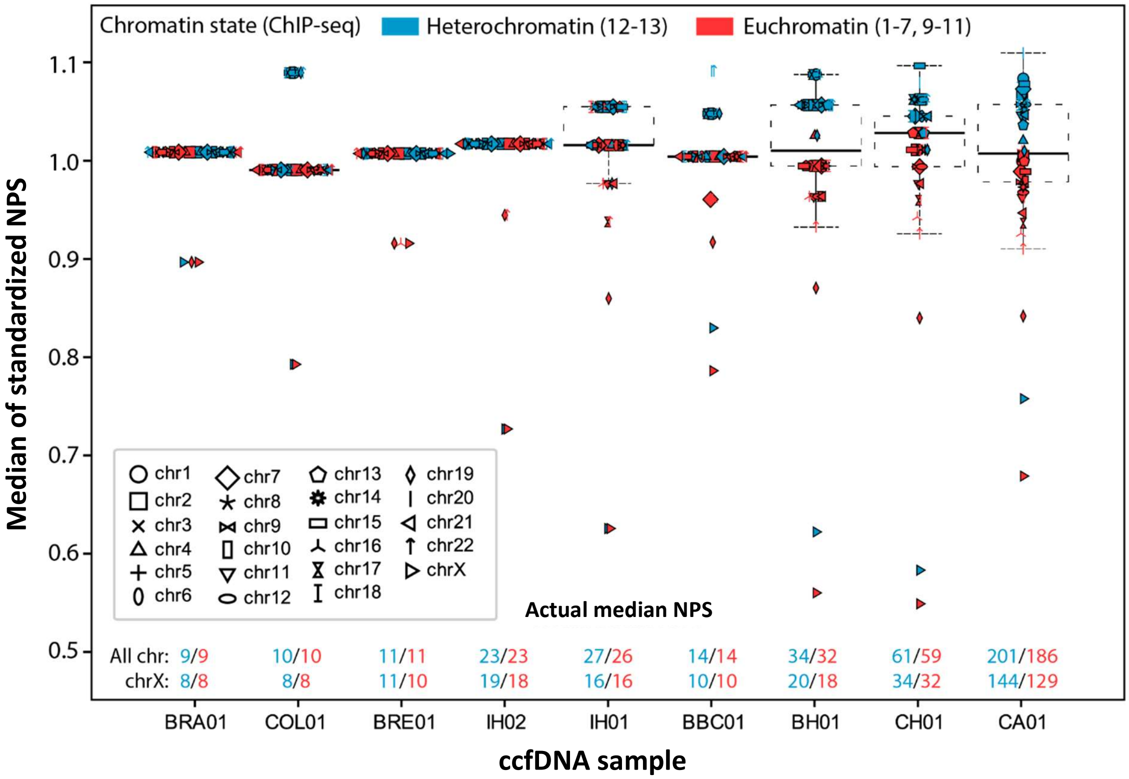


**Supplementary Figure S20. Box and whisker plots of NPS medians for NPPs occupying heterochromatin or euchromatin for each chromosome.** To account for differences in sequencing depths between samples, standardized normalized prominence score (NPS) was calculated by dividing the NPS of a nucleosome protection peak (NNP) by the median NPS of all NPPs called for a sample. The actual median NPS of heterochromatin (blue) and euchromatin (red) is also display for the autosomes and chrX of each sample. The bottom line of each box represents the 25th percentile, top line the 75th percentile, and thick middle line the median. Whiskers extend up to a maximum of 1.5 times the height of the box. Any values that fall outside this range are classified as outliers.

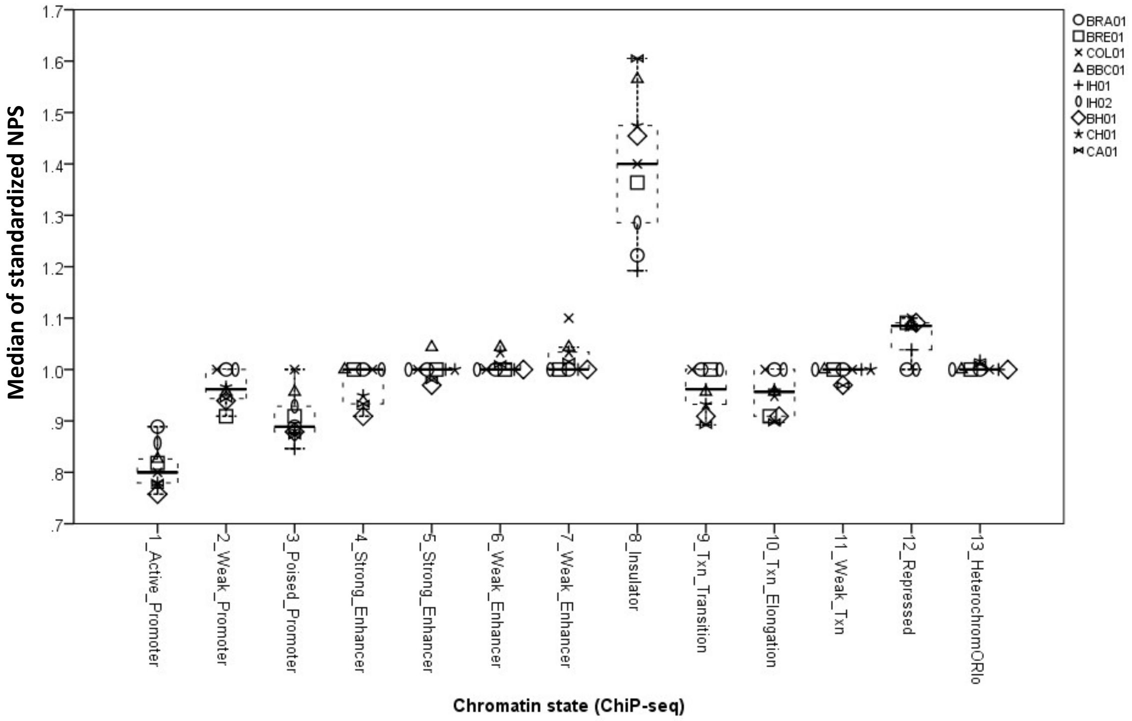


**Supplementary Figure S21. NPS medians by sample and chromatin state.** To account for differences in sequencing depths between samples, standardized normalized prominence score (NPS) was calculated by dividing the NPS of a nucleosome protection peak (NNP) by the median NPS of all NPPs called for a sample. The bottom line of each box represents the 25th percentile, top line the 75th percentile, and thick middle line the median. Whiskers extend up to a maximum of 1.5 times the height of the box. Any values that fall outside this range are classified as outliers.

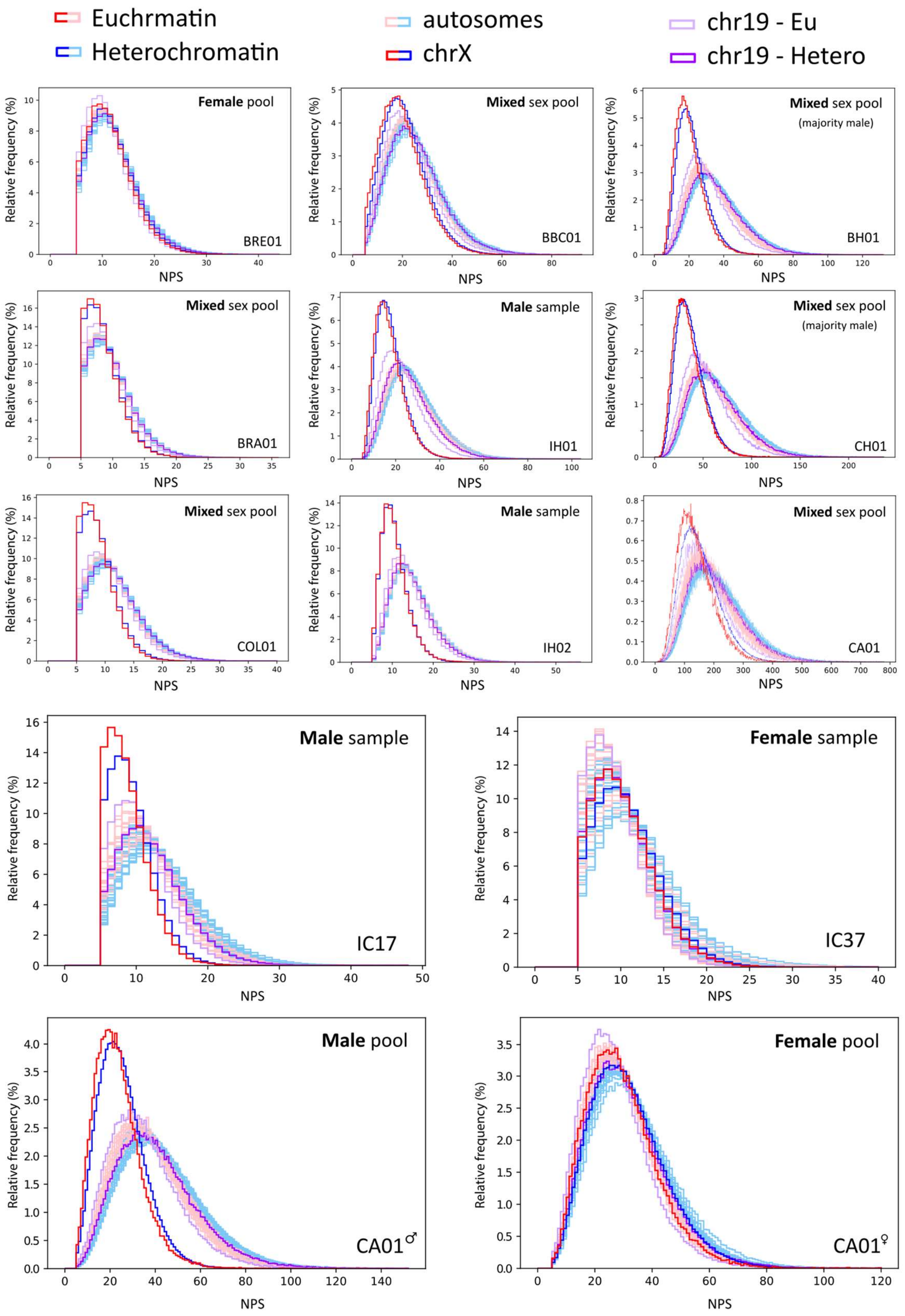


**Supplementary Figure S22. Nucleosome protection peak NPS histograms.** Histograms of normalized prominence score (NPS) of all nucleosome protection peaks (NPPs) genome-wide for each sample, separated into "Heterochromatin" composed of ChromHMM states 12 and 13 and "Euchromatin" composed of states 1-7, and 9-11 of the GM12878 cell line. The sex of a patient for individual samples, or sexes contained within sample pools, are displayed. For the BH01, CH01 and CA01 sample pools, the sexes displayed are based on NPS differences between chrX and autosomes, as the exact percentage breakdown of sexes contained in these pools is unknown.

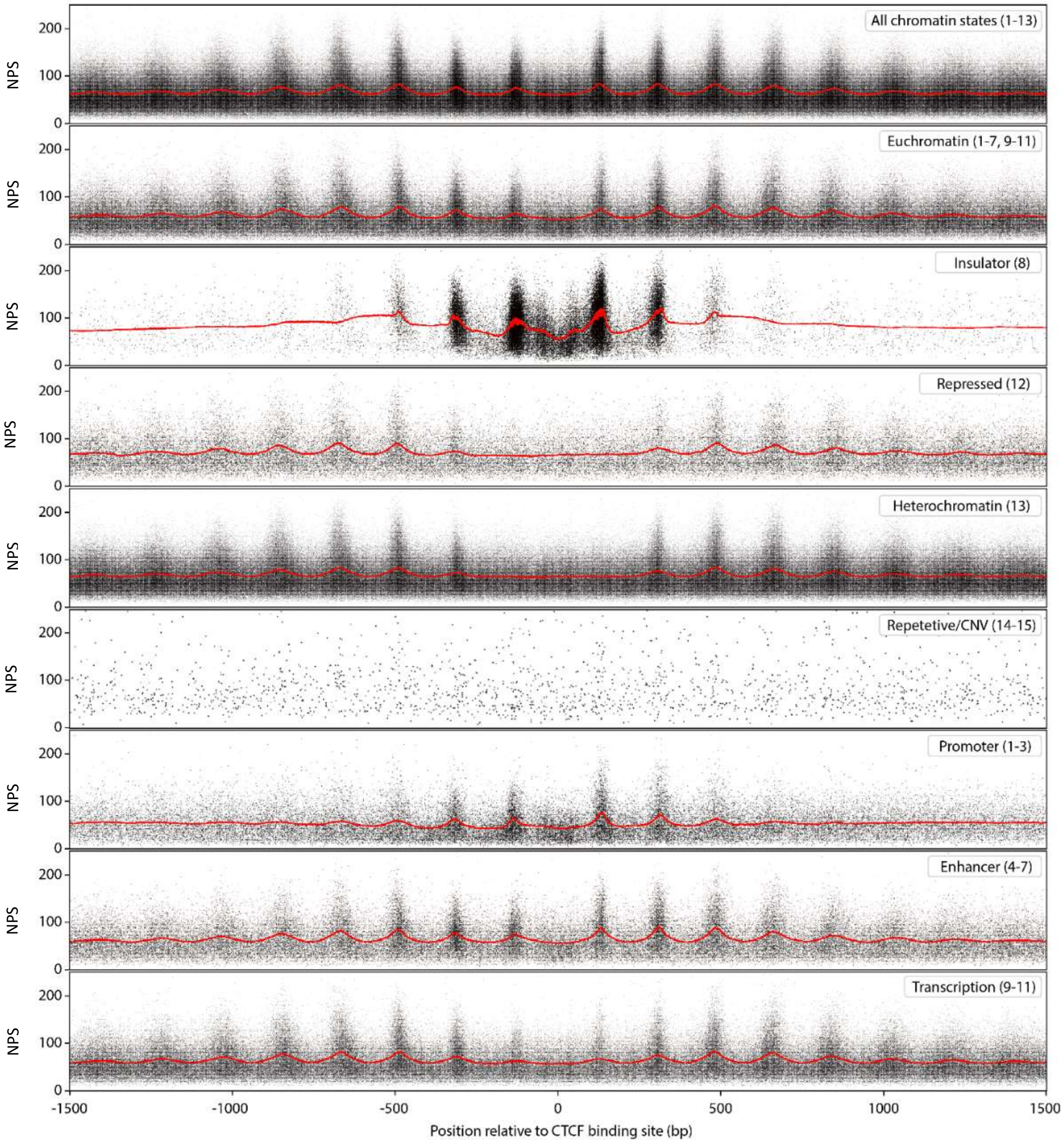


**Supplementary Figure S23. Strengths of nucleosome protection peaks (NPPs) relative to distance from CTCF binding sites**. CTCF binding sites are predictions by FIMO (v5.1.0) in the GRCh37 genome[4]. Chromatin states are those of the GM12878 B-lymphoblastoid cell line from Ernst et al. (2011)[2] and nucleosome protection peaks are those of the CH01 healthy sample pool from Snyder et. al. (2016)[3]. Scatter plot point sizes are varied for each plot relative to overall number of points to improve visualization and reduce saturation. Orange lines are moving average smoothed data to highlight peaks in normalized prominence score (NPS). Center peaks appear stronger in promoters (1-3) and enhancers (4-7) than transcription-associated chromatin states (9-11). As the ~250 bp spacing between these peaks remains, these are likely CTCF bound within enhancers or promoters[5], or insulators misclassified as euchromatin. Furthermore, these two center peaks do not appear in plots for heterochromatin states (states 12 and 13), suggesting that insulators are rarely misclassified as heterochromatin.

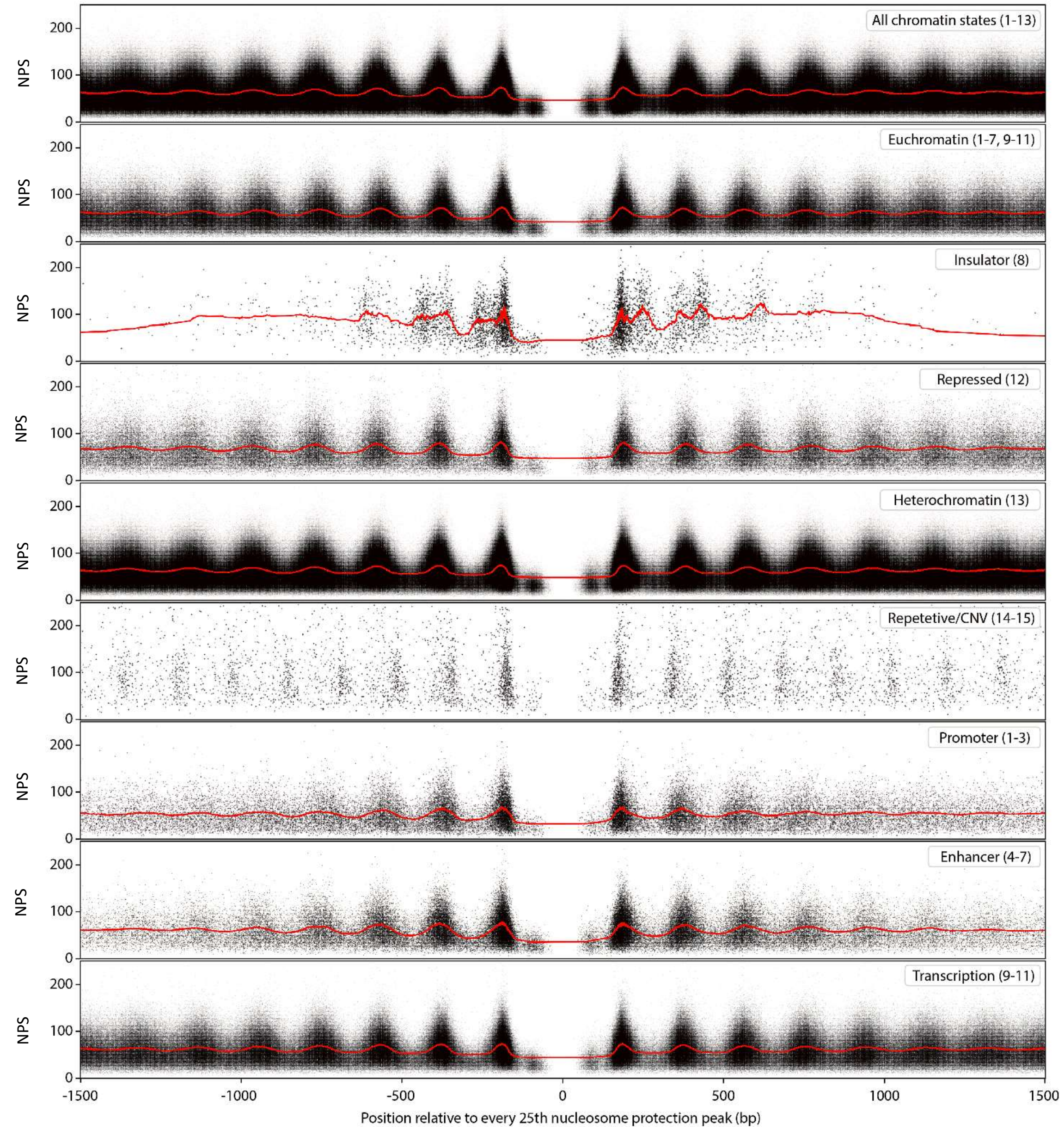


**Supplementary Figure S24. Strengths of nucleosome protection peaks (NPPs) relative to distance from every 25th NPP**. Chromatin states are those of the GM12878 B-lymphoblastoid cell line from Ernst et al. (2011) and nucleosome protection peaks are those of the CH01 healthy sample pool from Snyder et. al. (2016). Scatter plot point sizes are varied relative to overall number of points to improve visualization and reduce saturation. Orange lines are moving average smoothed data to highlight peaks in normalized prominence score (NPS). The distributions of these nucleosome protection meta-peaks are broader for NPP alignments than those observed for CTCF, presumably because nucleosome position calls are less accurate than sequence-specific motif positioning. Furthermore, unlike the CTCF alignments, the distances of the two peaks adjacent to the center are ≈185 bp rather than the ≈125 bp observed for CTCF binding sites. However, in addition to the ≈185 bp spacing, the insulator plot contains peaks spaced at ≈250 bp, corresponding to the distance created by CTCF binding between nucleosomes.

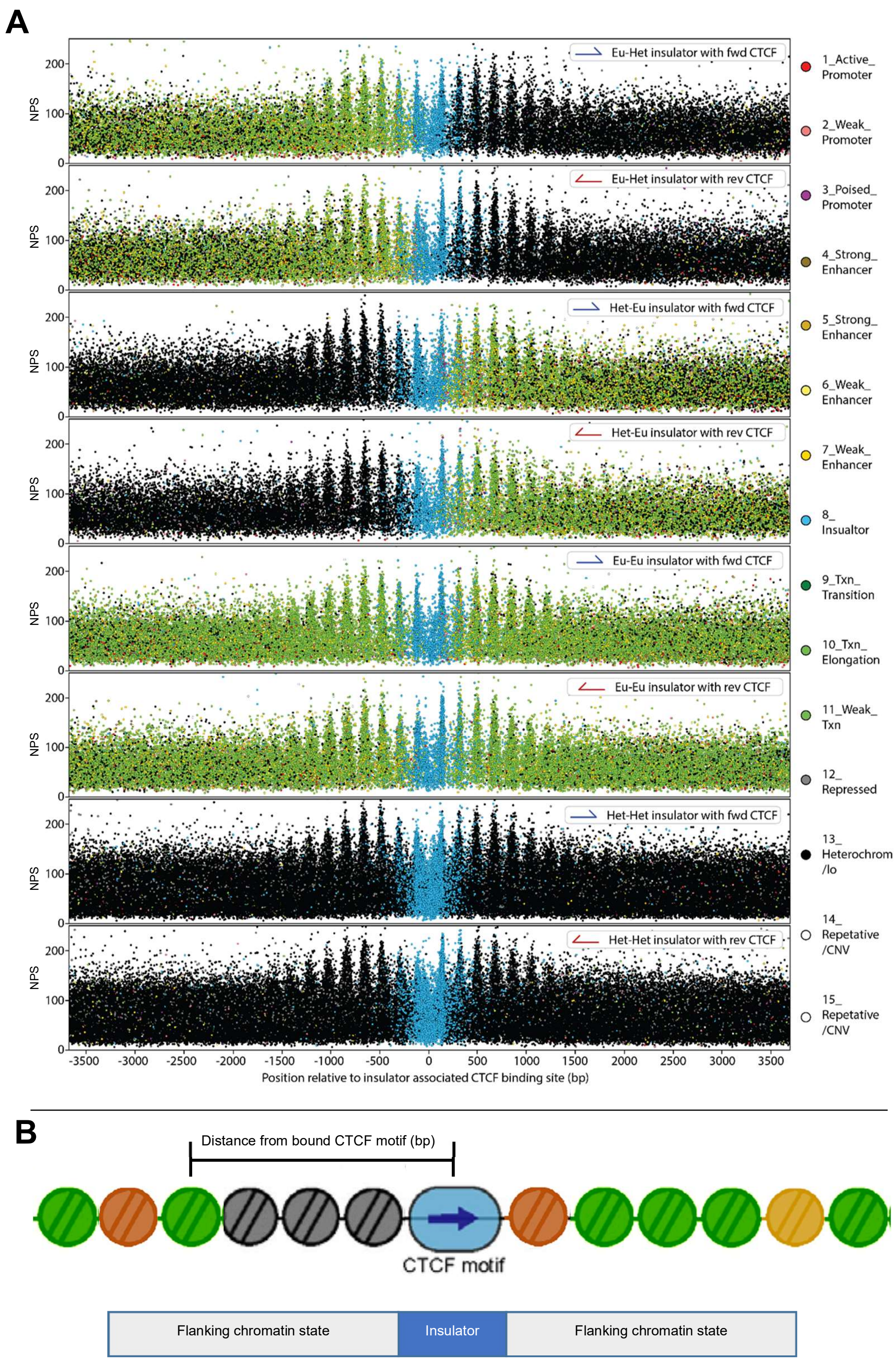


**Supplementary Figure S25. Strengths and chromatin states of nucleosome protection peaks (NPPs) relative to distance from CTCF/insulators.** (**A**) Insulators were categorized based on the state of the two flanking chromatin regions, as well as the direction of the CTCF binding site. Eu = euchromatin, Het = heterochromatin. Chromatin states are those of the GM12878 B-lymphoblastoid cell line from Ernst et al. (2011)[2] and nucleosome protection peaks are those of the CH01 healthy sample pool from Snyder et. al. (2016)[3]. NPS = normalized prominence score. Scatter plot point are coloured based on the chromatin state of each NPP. (**B**) Example: Heterochromatin-Euchromatin insulator with forward CTCF-binding motif

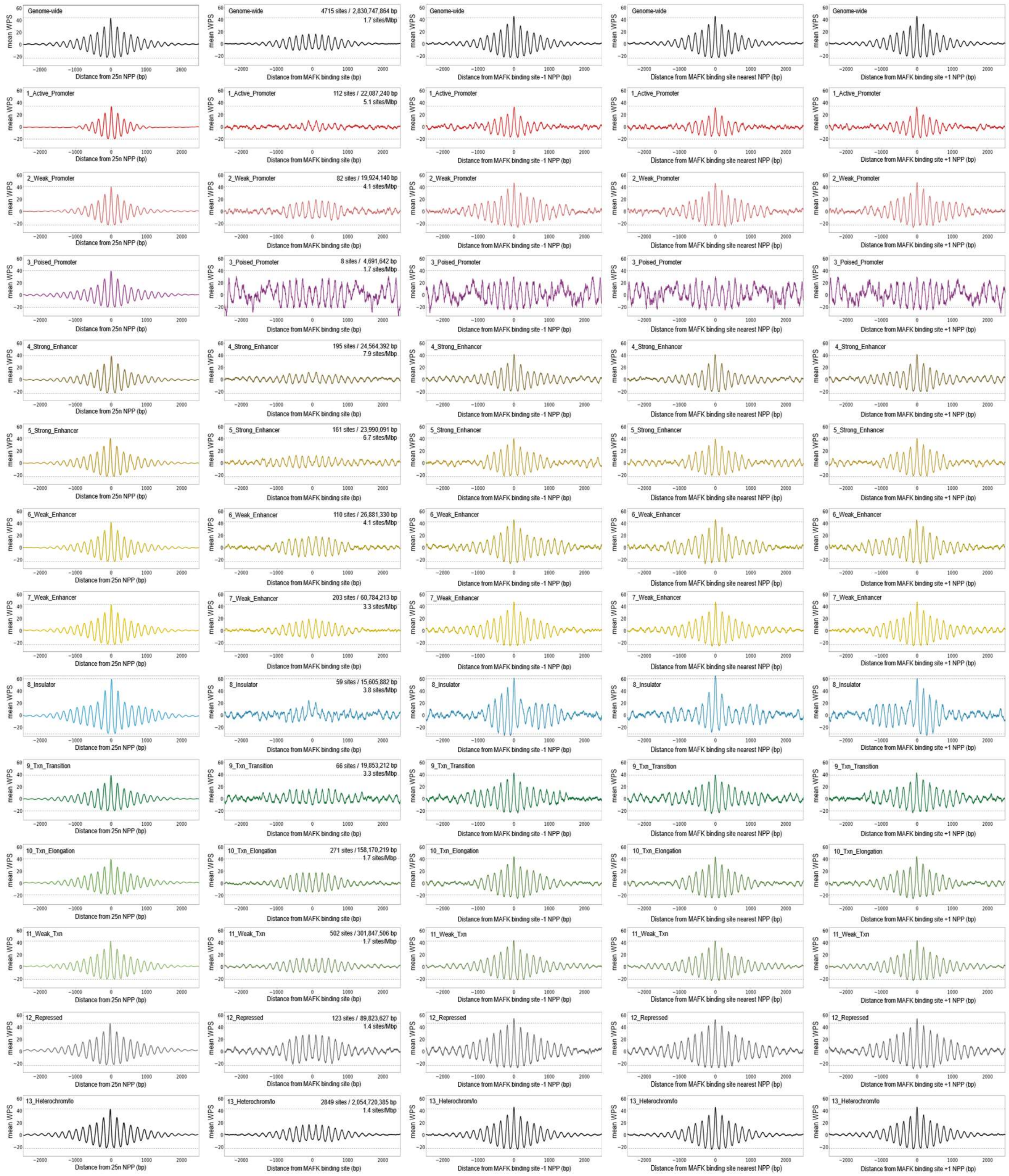


**Supplementary Figure S26. Mean WPS relative to distance from MAFK bindings sites**. Alignments of mean WPS relative to distance from MAFK binding sites within ENCODE ChIP-seq peaks and the NPPs flanking these genomic features. Binding sites are predicted by FIMO (v5.3.3) in the GRCh37 genome[25]. All sites are aligned at the 0 position after adjusting for binding site orientation. NPPs are those of the CH01 healthy sample pool from Snyder et. al. (2016)[10]. Chromatin states are those of Ernst et al. (2011)[20]. ENCODE ChIP-seq data are from the GM12878 B-lymphoblastoid cell line.

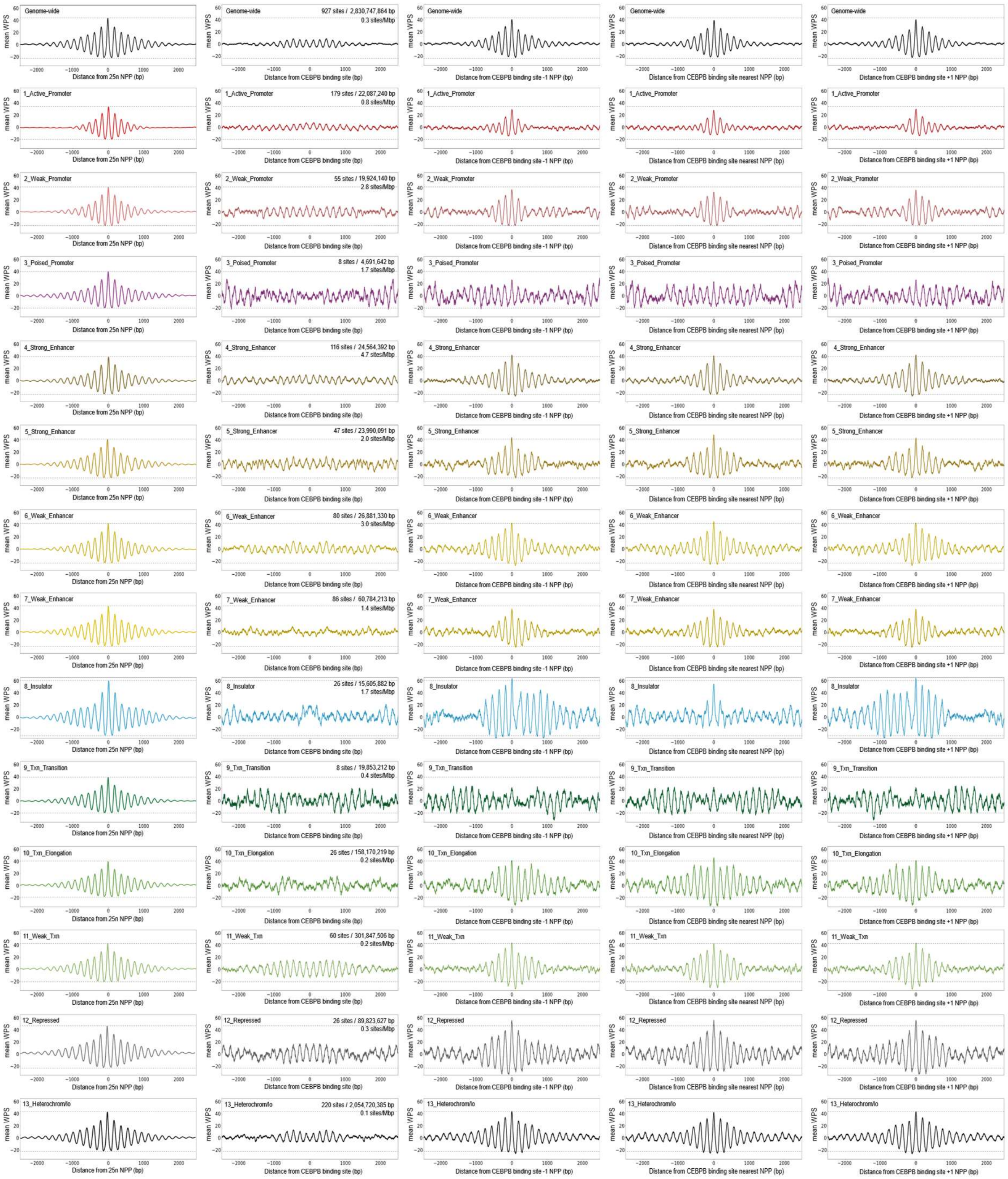


**Supplementary Figure S27. Mean WPS relative to distance from CEBPB bindings sites**. Alignments of mean WPS relative to distance from CEBPB binding sites within ENCODE ChIP-seq peaks and the NPPs flanking these genomic features. Binding sites are predicted by FIMO (v5.3.3) in the GRCh37 genome[25]. All sites are aligned at the 0 position after adjusting for binding site orientation. NPPs are those of the CH01 healthy sample pool from Snyder et. al. (2016)[10]. Chromatin states are those of Ernst et al. (2011)[20]. ENCODE ChIP-seq data are from the GM12878 B-lymphoblastoid cell line.

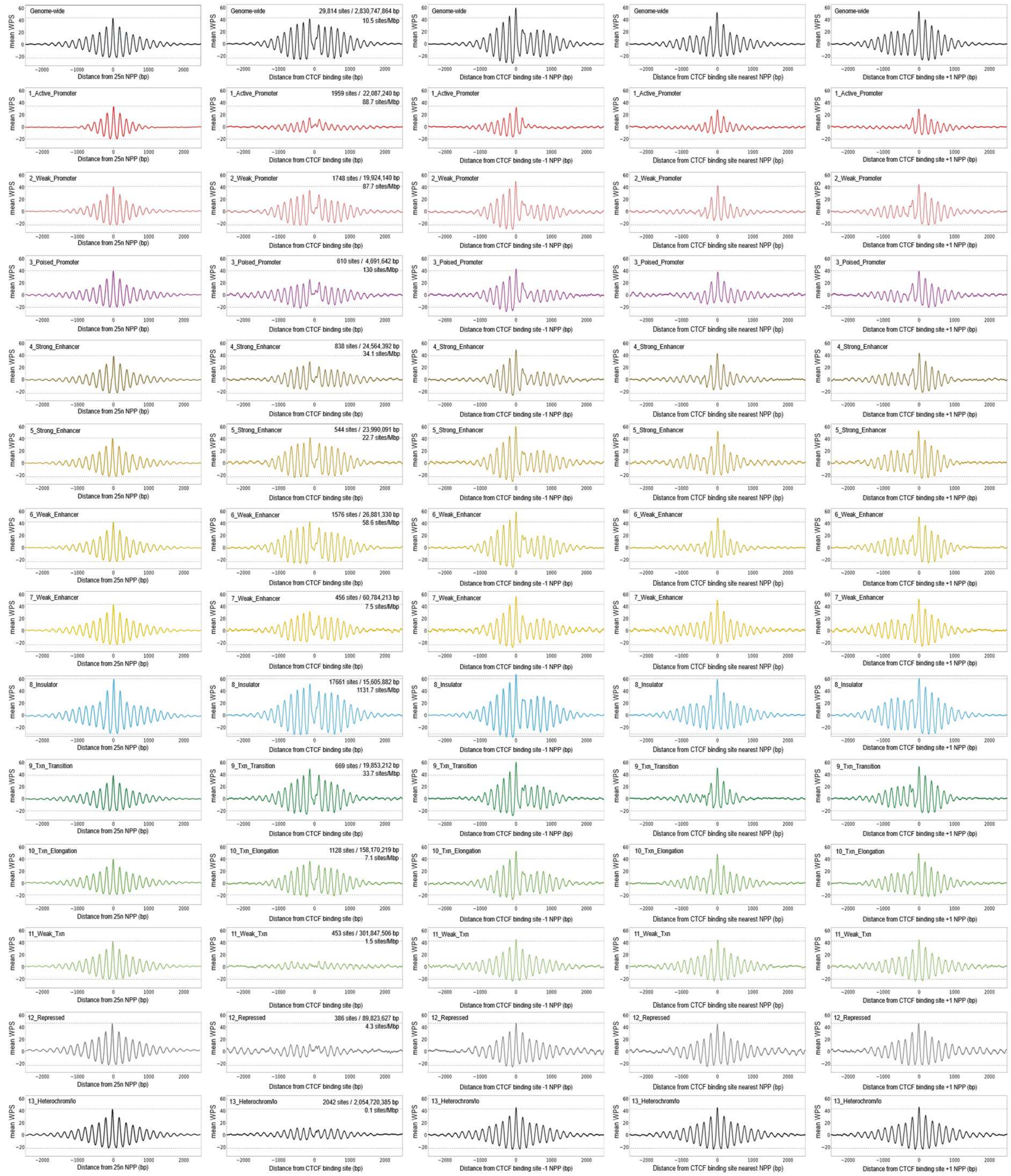


**Supplementary Figure S28. Mean WPS relative to distance from CTCF bindings sites**. Alignments of mean WPS relative to distance from CTCF binding sites within ENCODE ChIP-seq peaks and the NPPs flanking these genomic features. Binding sites are predicted by FIMO (v5.3.3) in the GRCh37 genome[25]. All sites are aligned at the 0 position after adjusting for binding site orientation. NPPs are those of the CH01 healthy sample pool from Snyder et. al. (2016)[10]. Chromatin states are those of Ernst et al. (2011)[20]. ENCODE ChIP-seq data are from the GM12878 B-lymphoblastoid cell line.

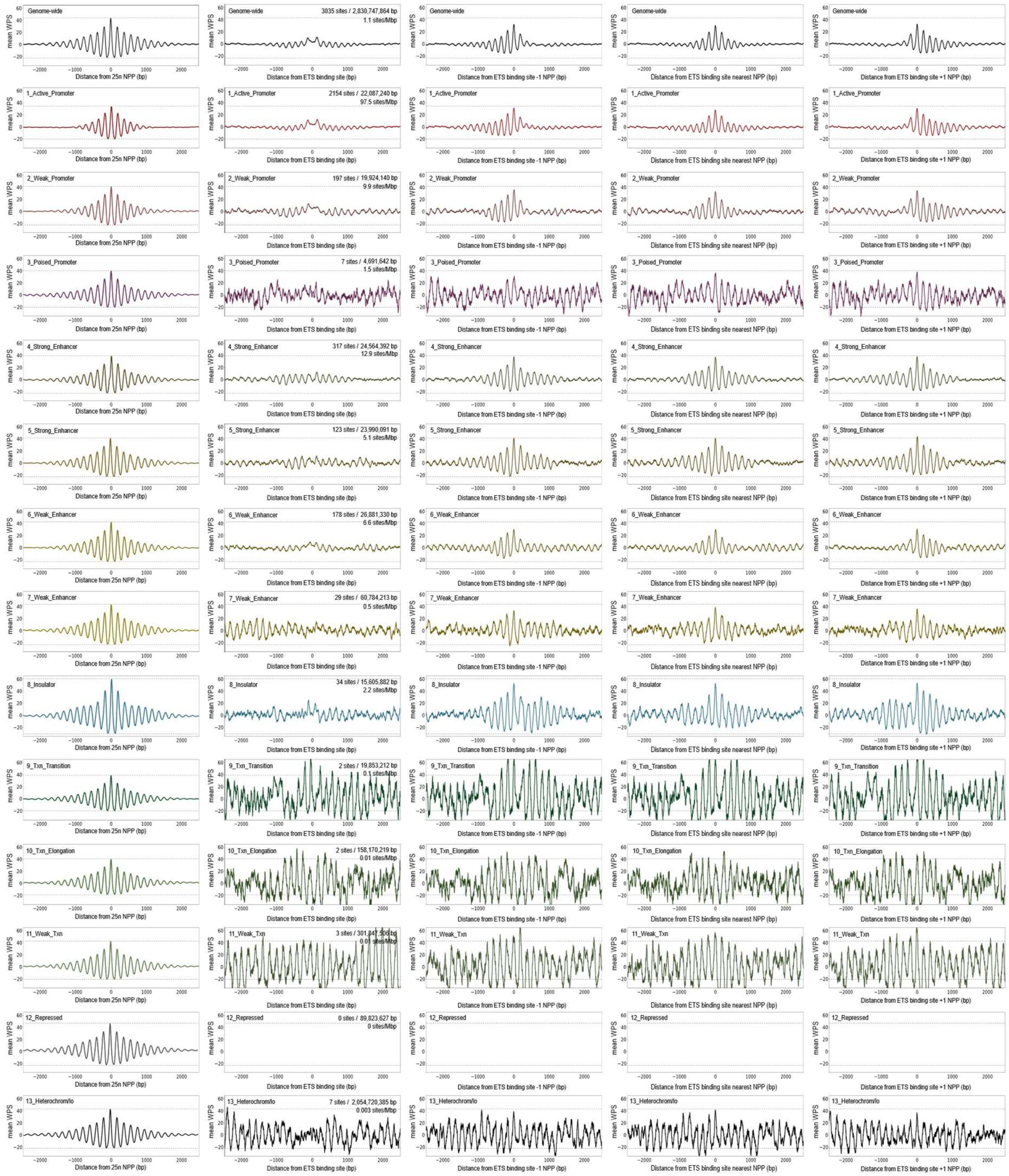

**Supplementary Figure S29. Mean WPS relative to distance from ETS bindings sites**. Alignments of mean WPS relative to distance from ETS binding sites within ENCODE ChIP-seq peaks and the NPPs flanking these genomic features. Binding sites are predicted by FIMO (v5.3.3) in the GRCh37 genome[25]. All sites are aligned at the 0 position after adjusting for binding site orientation. NPPs are those of the CH01 healthy sample pool from Snyder et. al. (2016)[10]. Chromatin states are those of Ernst et al. (2011)[20]. ENCODE ChIP-seq data are from the GM12878 B-lymphoblastoid cell line.

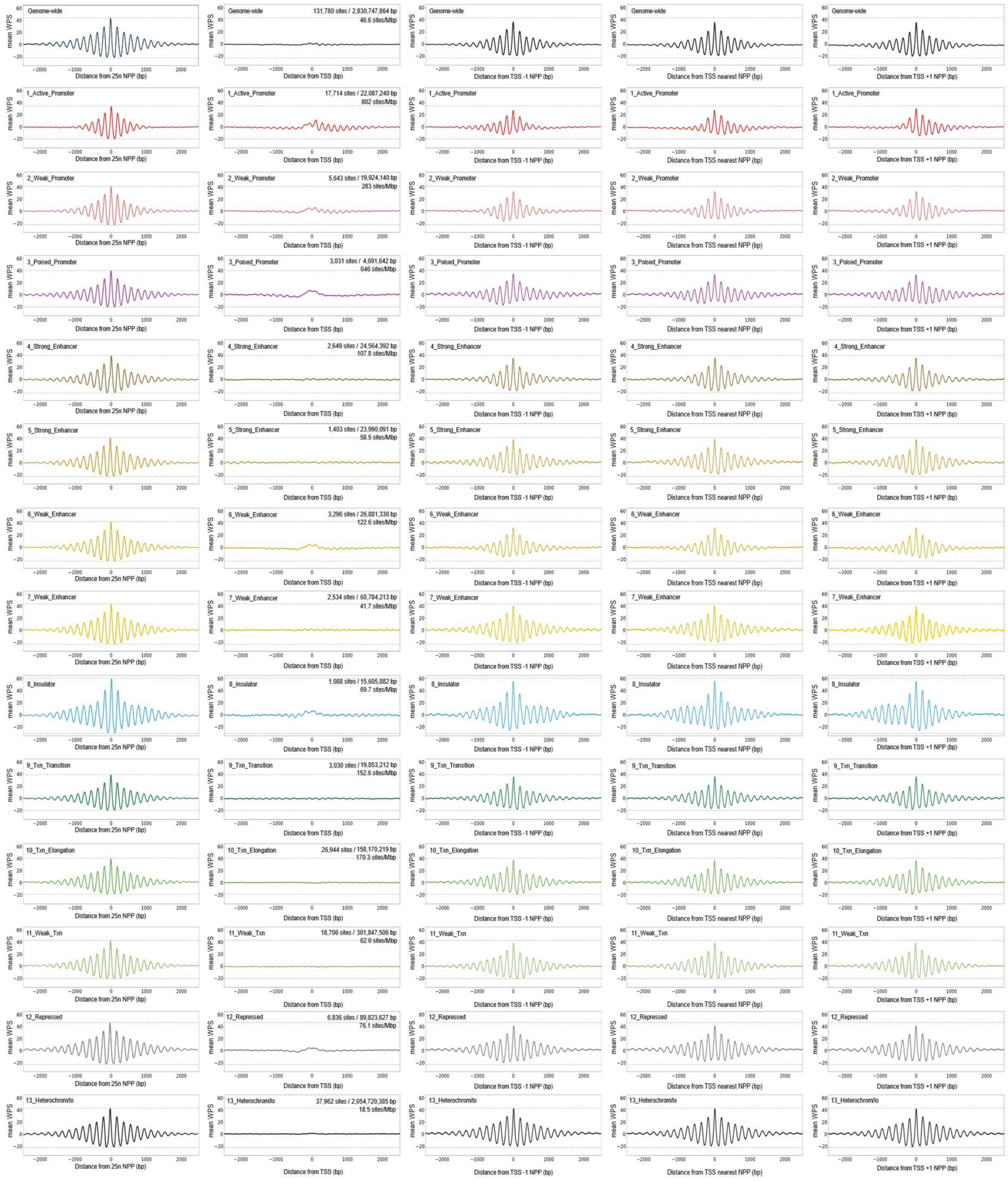


**Supplementary Figure S30. Mean WPS relative to distance from transcription start sites**. Alignments of mean WPS relative to distance from transcription start sites (TSSs) and the NPPs flanking these genomic features. TSSs are aligned at the 0 position after adjusting for transcription orientation. NPPs are those of the CH01 healthy sample pool from Snyder et. al. (2016)[10]. Chromatin states are those of Ernst et al. (2011)[20] from the GM12878 B-lymphoblastoid cell line.

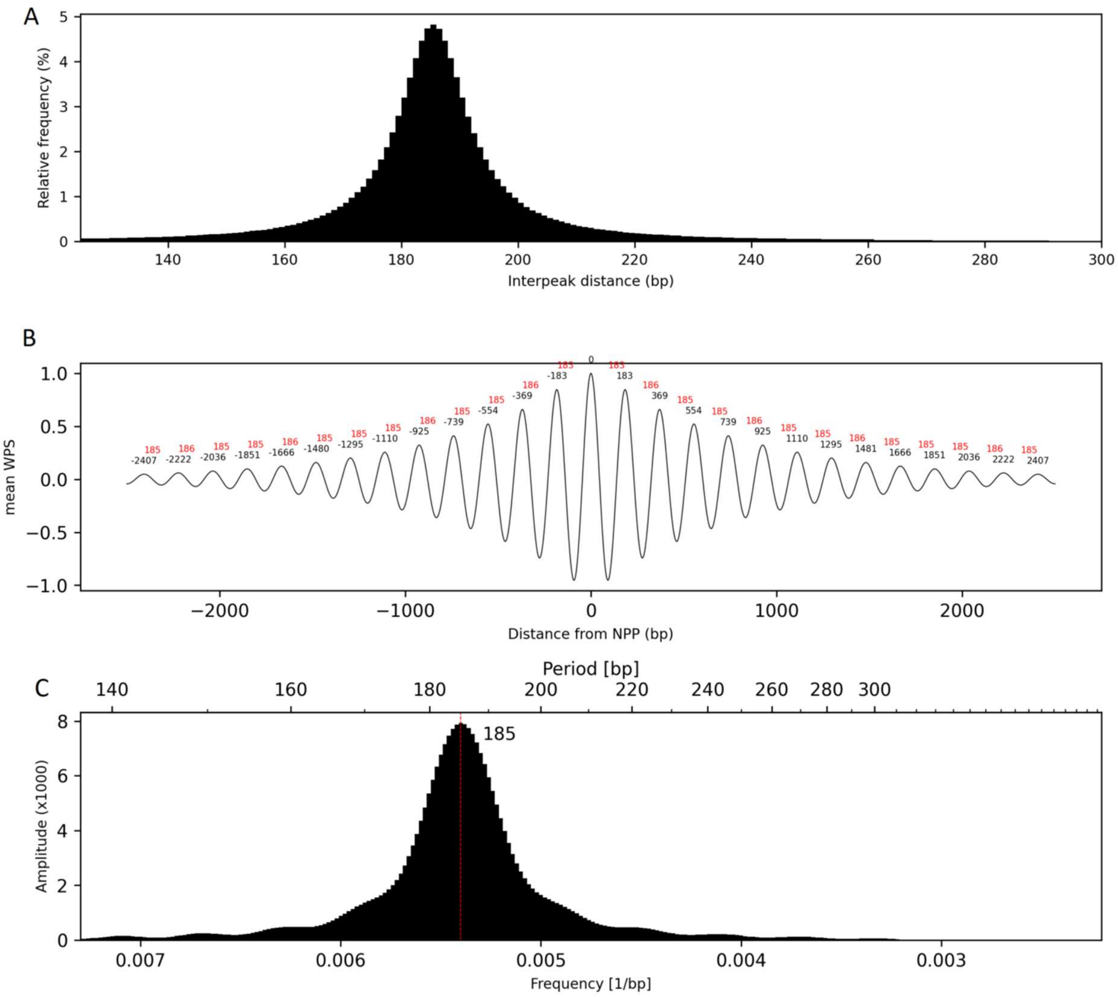


**Supplementary Figure S31. Sine wave phasing simulation using Cauchy distribution of interpeak distances.** (**A**) Cauchy distribution of interpeak distances with mode of 185 bp. (**B**) Averaged aggregate of all sine waves with interpeak distances from plot A. (**C**) Fourier transform of plot B.

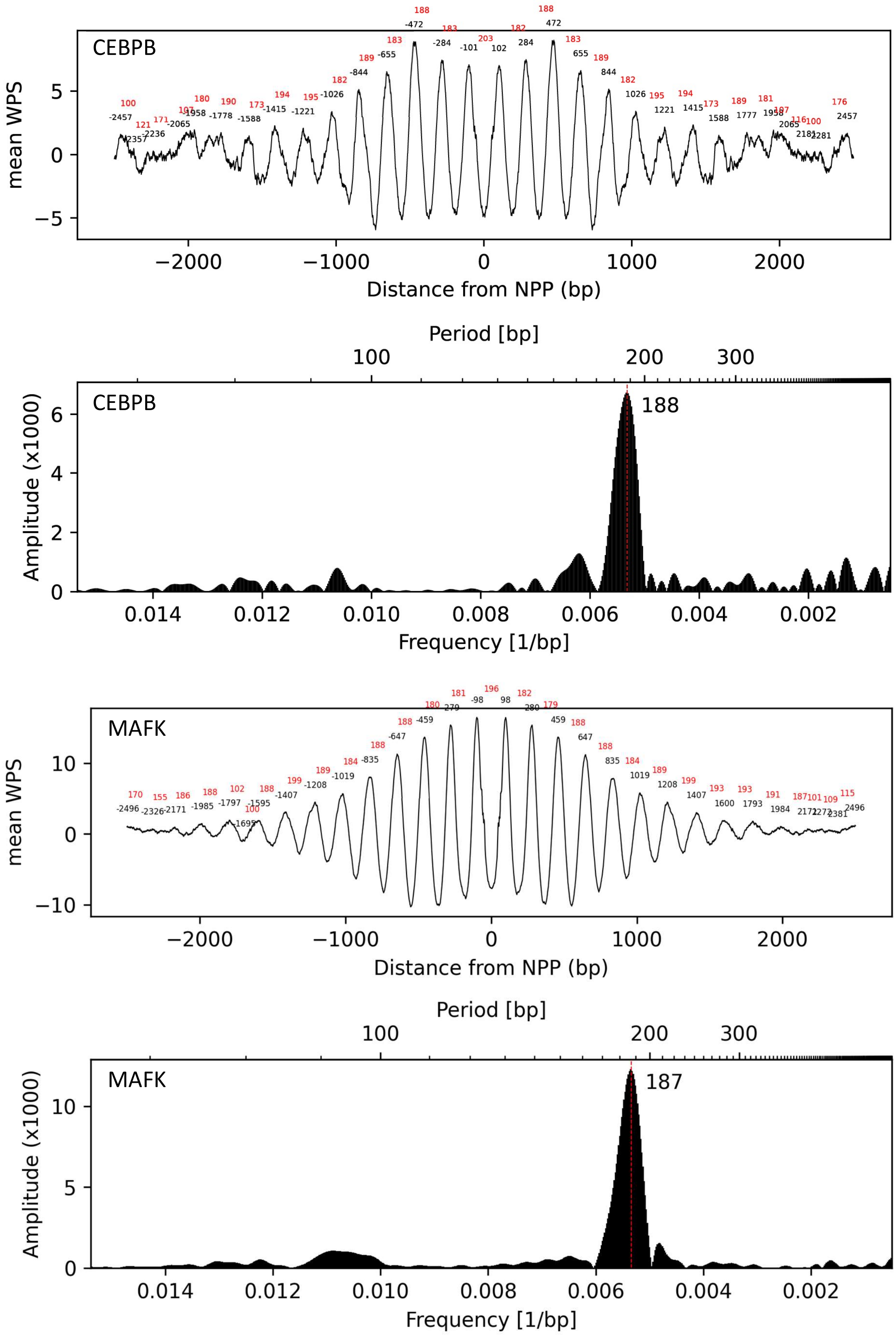
CEBPB
mean WPS
Distance from NPP (bp)
Period [bp]
Amplitude (x1000)
188
Frequency [1/bp]
MAFK
mean WPS
Distance from NPP (bp)
Period [bp]
Amplitude (x1000)
187
Frequency [1/bp]

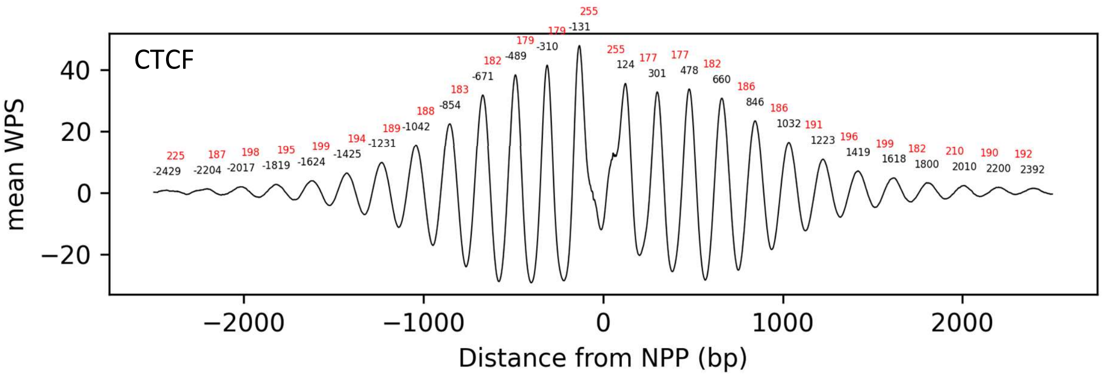
CTCF
mean WPS
40
20
0
−20
−2000
−1000
0
1000
2000
Distance from NPP (bp)

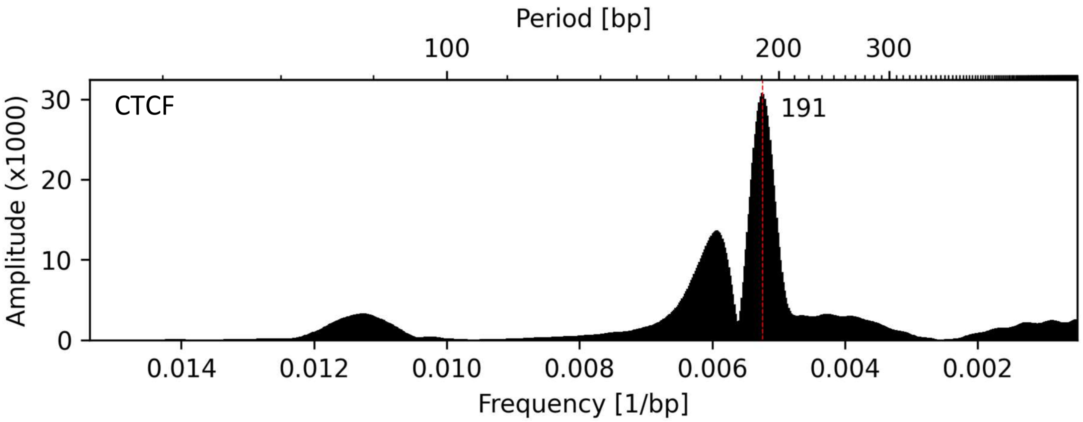
Period [bp]
100
200
300
CTCF
191
Amplitude (x1000)
30
20
10
0
0.014
0.012
0.010
0.008
0.006
0.004
0.002
Frequency [1/bp]

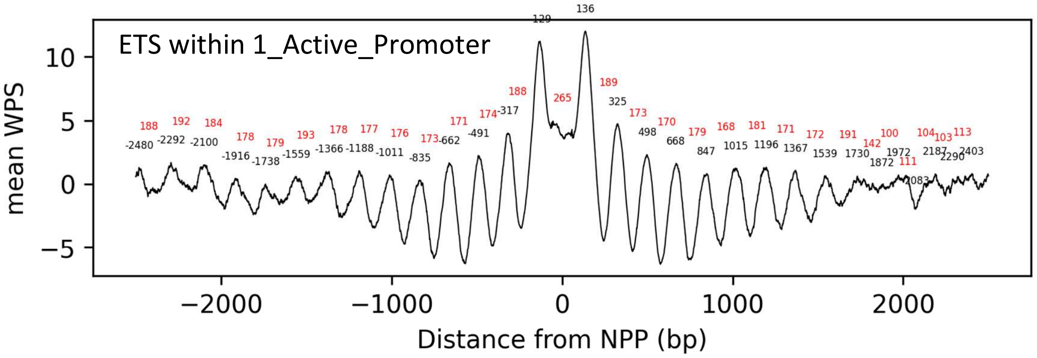
ETS within 1_Active_Promoter
mean WPS
10
5
0
−5
−2000
−1000
0
1000
2000
Distance from NPP (bp)

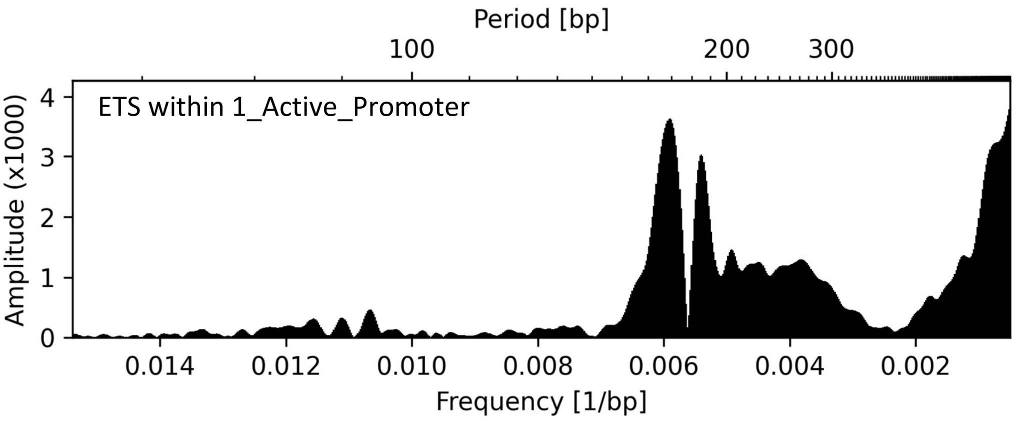
Period [bp]
100
200
300
ETS within 1_Active_Promoter
Amplitude (x1000)
4
3
2
1
0
0.014
0.012
0.010
0.008
0.006
0.004
0.002
Frequency [1/bp]

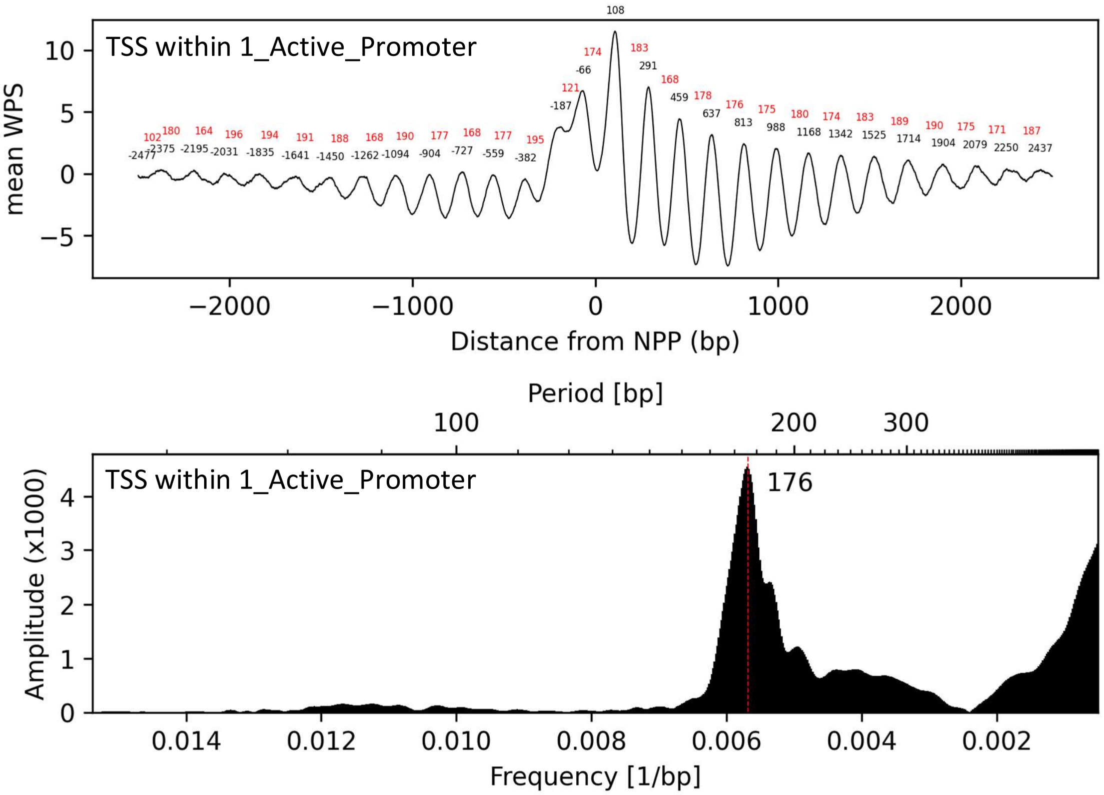


**Supplementary Figure S32. Fourier transforms of mean WPS aligned binding sites within ENCODE ChIP-seq peaks and ChromHMM chromatin states**. Binding sites are predicted by FIMO (v5.3.3) in the GRCh37 genome. All sites are aligned at the 0 position after adjusting for binding site orientation. Distances of each peak from position 0 are in black and distances between peaks are in red. The mode interpeak distance (period) are displayed for each Fourier transform. NPPs are those of the CH01 healthy sample pool from Snyder et. al. (2016). Chromatin states are those of Ernst et al. (2011). ENCODE ChIP-seq data are from the GM12878 B-lymphoblastoid cell line.

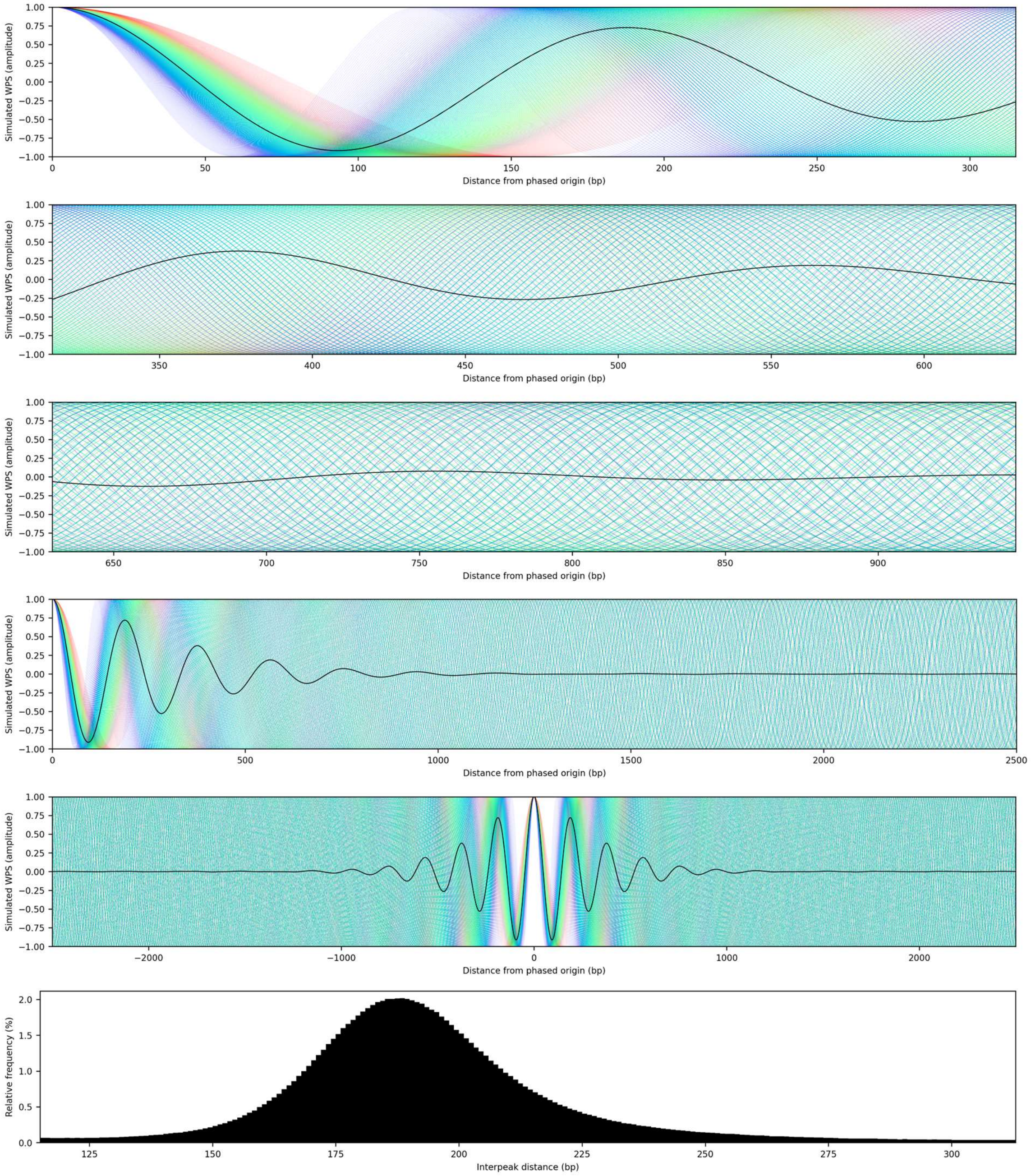


**Supplementary Figure S33. Sine wave phasing simulation using inter-NPP distance proportions measured in CH01 sample.** Rainbows are composed of sine waves at different frequencies. Each coloured line is a sine wave with an interpeak distance between 115 bp [violet] and 315 bp [red]), based on the major peak in nucleosome spacing histograms. The black line is an averaged aggregate of all the NRL-based sine waves depicted in rainbow. Sine waves for each interpeak distance were included multiple times in this wave summation based on the relative frequency with which these inter-NPP distances appeared genome-wide in the CH01 sample (see interpeak distance histogram). For example, waves with 187 bp interpeak distances were included 201 times [relative frequency = 2.01%] whereas waves with 295 bp interpeak distances were included 4 times [relative frequency = 0.4%]). Multiple plots at different scales are included for greater clarity of wave dispersion and fading of aggerate signal.

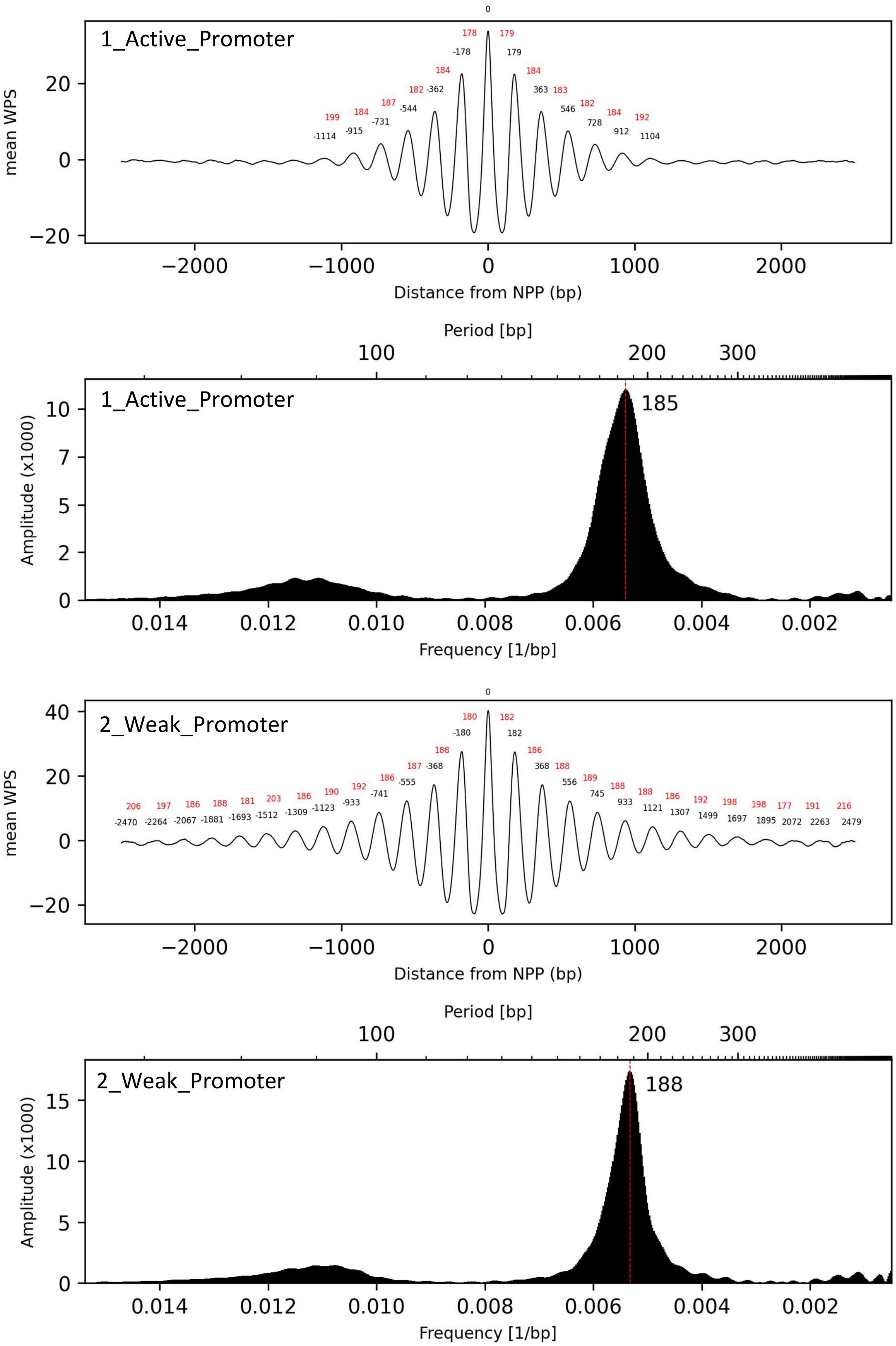
1_Active_Promoter
mean WPS
Distance from NPP (bp)
0
178
179
-178
179
184
184
182 -362
363 183
187
-544
546 182
184
-731
728
184
199
-915
912
192
-1114
1104
20
0
−20
−2000
−1000
0
1000
2000
Period [bp]
100
200
300
1_Active_Promoter
185
Amplitude (x1000)
10
7
5
2
0
0.014
0.012
0.010
0.008
0.006
0.004
0.002
Frequency [1/bp]
2_Weak_Promoter
mean WPS
0
180
182
-180
182
188
186
187 -368
368 188
186
-555
556
189
192
-741
745
188
190
-933
933
188
206
197
186
188
181
203
186
186
192
198
198
177
191
216
-2470
-2264
-2067
-1881
-1693
-1512
-1309
-1123
1121
1307
1499
1697
1895
2072
2263
2479
40
20
0
−20
−2000
−1000
0
1000
2000
Distance from NPP (bp)
Period [bp]
100
200
300
2_Weak_Promoter
188
Amplitude (x1000)
15
10
5
0
0.014
0.012
0.010
0.008
0.006
0.004
0.002
Frequency [1/bp]

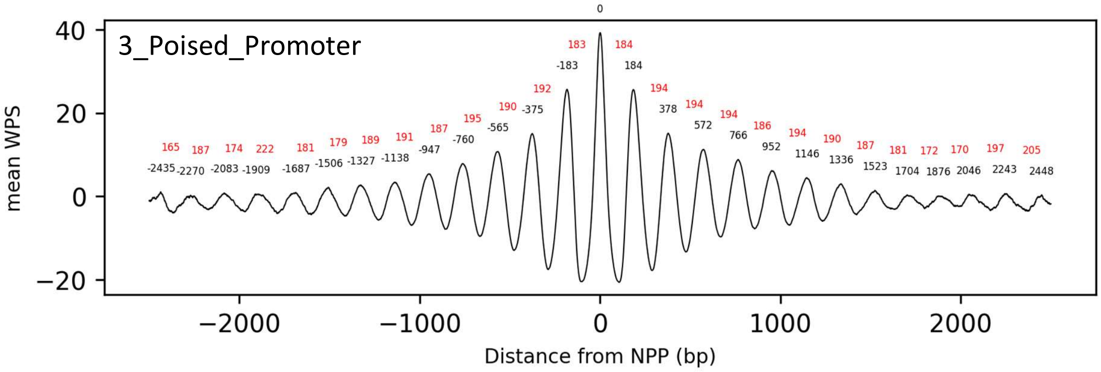
3_Poised_Promoter
mean WPS
Distance from NPP (bp)
0
183 184
-183 184
192 194
190 -375 378 194
195 -565 572 194
187 -760 766 186
165 187 174 222 181 179 189 191 -947 952 194 190 187 181 172 170 197 205
-2435 -2270 -2083 -1909 -1687 -1506 -1327 -1138 1146 1336 1523 1704 1876 2046 2243 2448

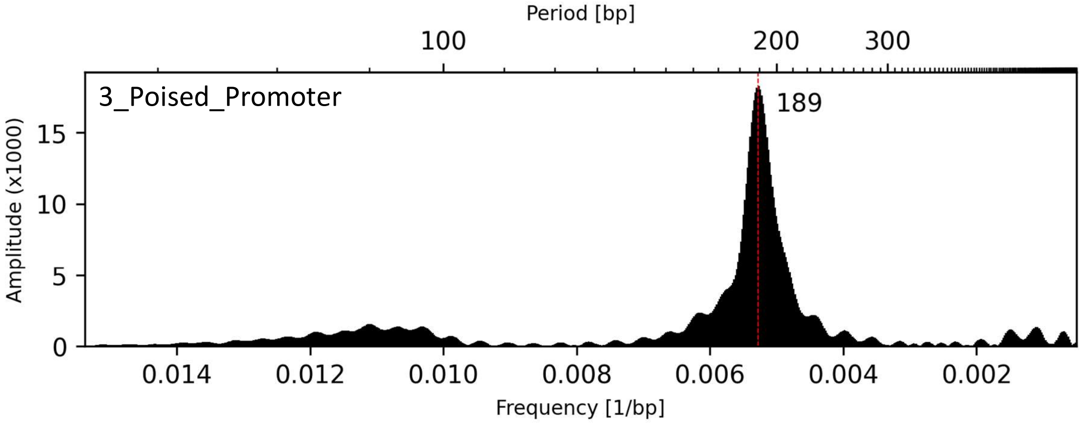
Period [bp]
100
200
300
3_Poised_Promoter
189
Amplitude (x1000)
Frequency [1/bp]

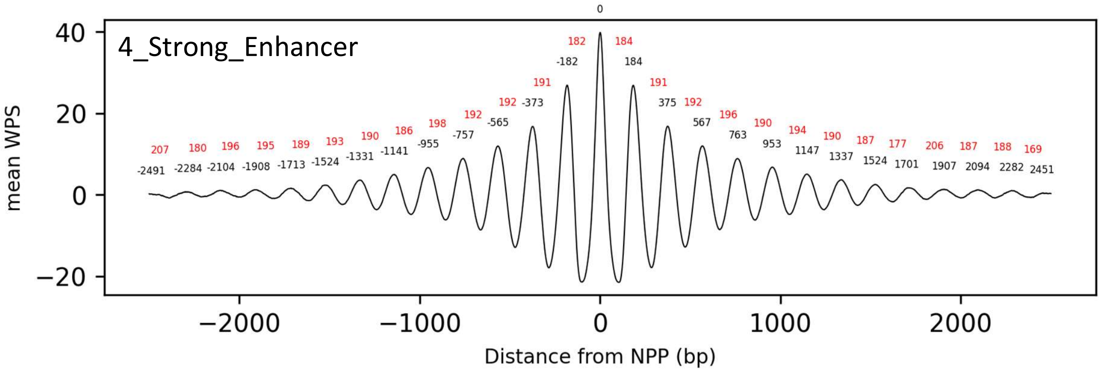
4_Strong_Enhancer
mean WPS
Distance from NPP (bp)
0
182 184
-182 184
191 191
192 -373 375 192
192 -565 567 196
198 -757 763 190
207 180 196 195 189 193 190 186 -955 953 194 190 187 177 206 187 188 169
-2491 -2284 -2104 -1908 -1713 -1524 -1331 -1141 1147 1337 1524 1701 1907 2094 2282 2451

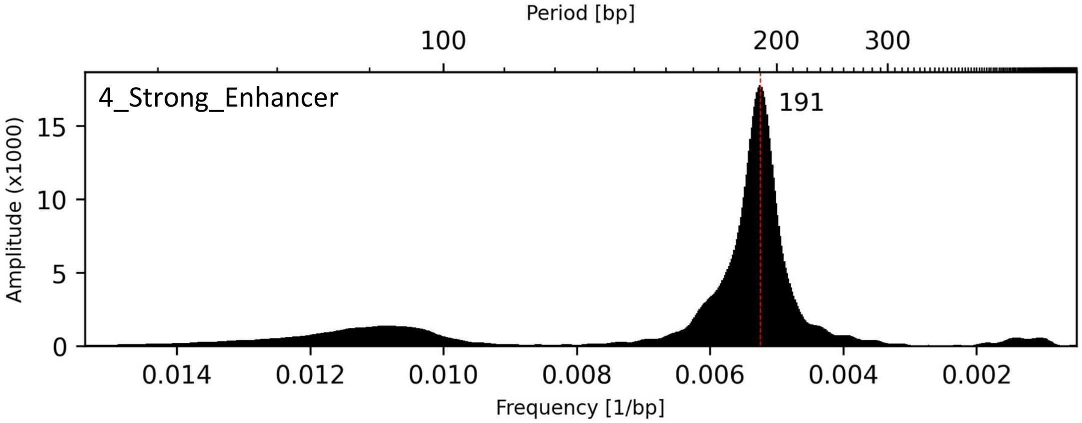
Period [bp]
100
200
300
4_Strong_Enhancer
191
Amplitude (x1000)
Frequency [1/bp]

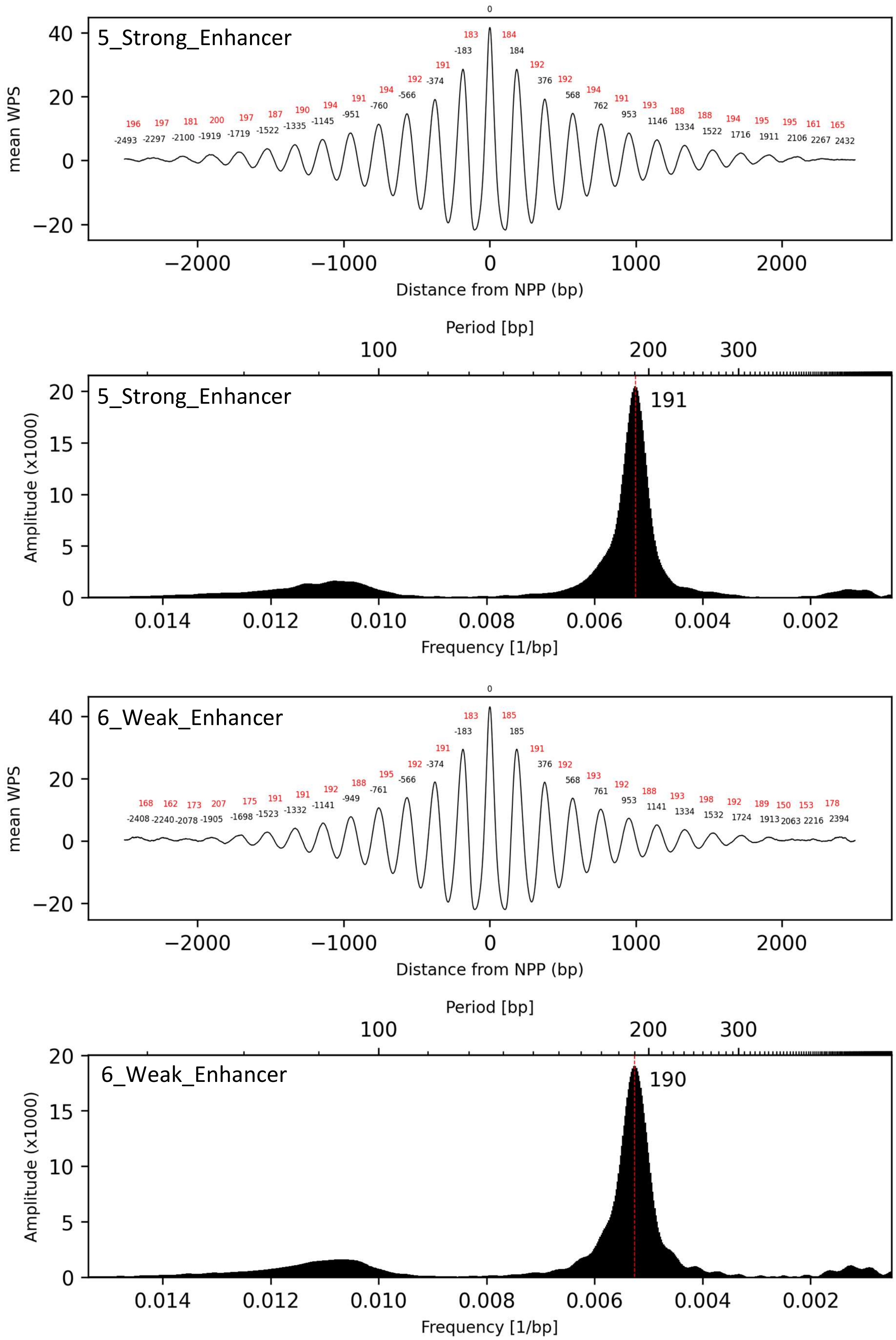
5_Strong_Enhancer
mean WPS
Distance from NPP (bp)
0
183 184
-183 184
191 192
-374 376
192 192
-566 568
194 194
-760 762
191 191
-951 953
194 193
-1145 1146
190 188
-1335 1334
187 188
-1522 1522
197 194
-1719 1716
200 195
-1919 1911
181 195
-2100 2106
197 161
-2297 2267
196 165
-2493 2432
Period [bp]
100
200
300
191
Amplitude (x1000)
Frequency [1/bp]
0.014
0.012
0.010
0.008
0.006
0.004
0.002
6_Weak_Enhancer
0
183 185
-183 185
191 191
-374 376
192 192
-566 568
195 193
-761 761
188 192
-949 953
192 188
-1141 1141
191 193
-1332 1334
191 198
-1523 1532
175 192
-1698 1724
207 189
-1905 1913
173 150
-2078 2063
162 153
-2240 2216
168 178
-2408 2394
190

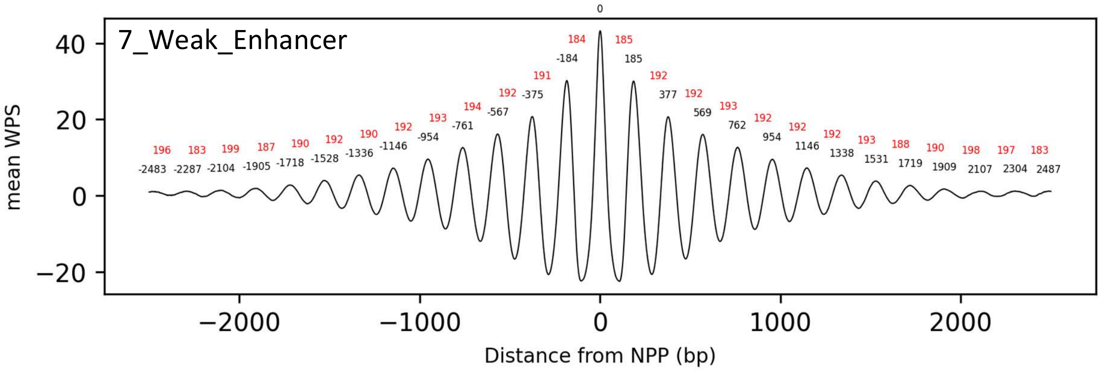
7_Weak_Enhancer
mean WPS
Distance from NPP (bp)

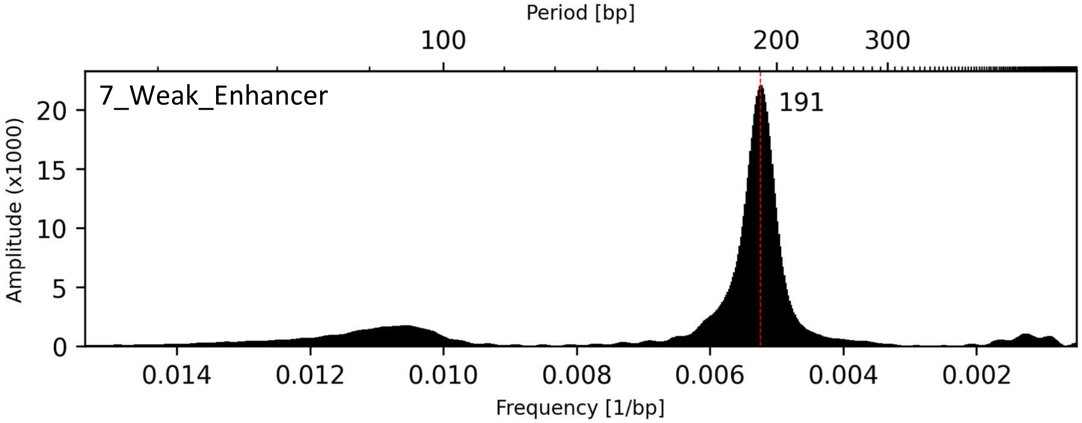
Period [bp]
7_Weak_Enhancer
191
Amplitude (x1000)
Frequency [1/bp]

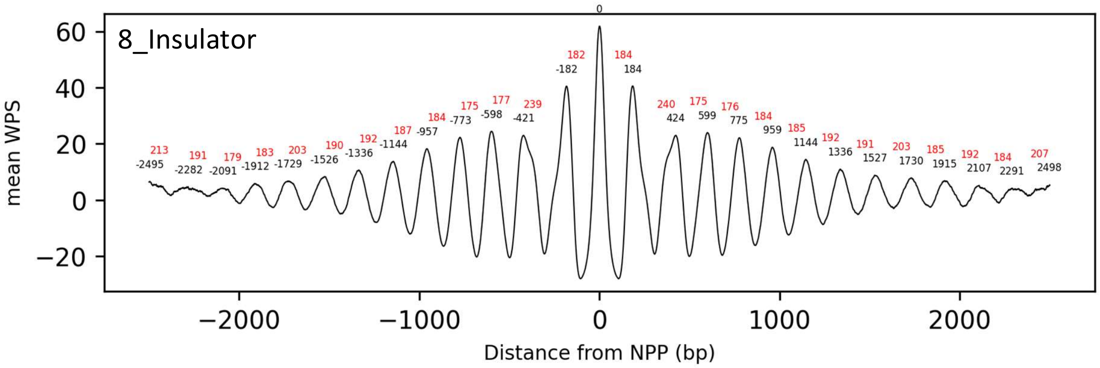
8_Insulator
mean WPS
Distance from NPP (bp)

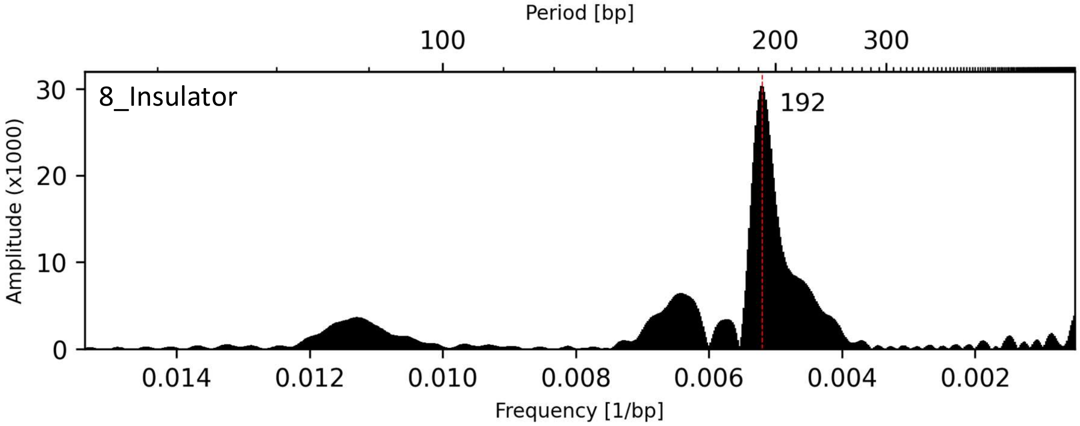
Period [bp]
8_Insulator
192
Amplitude (x1000)
Frequency [1/bp]

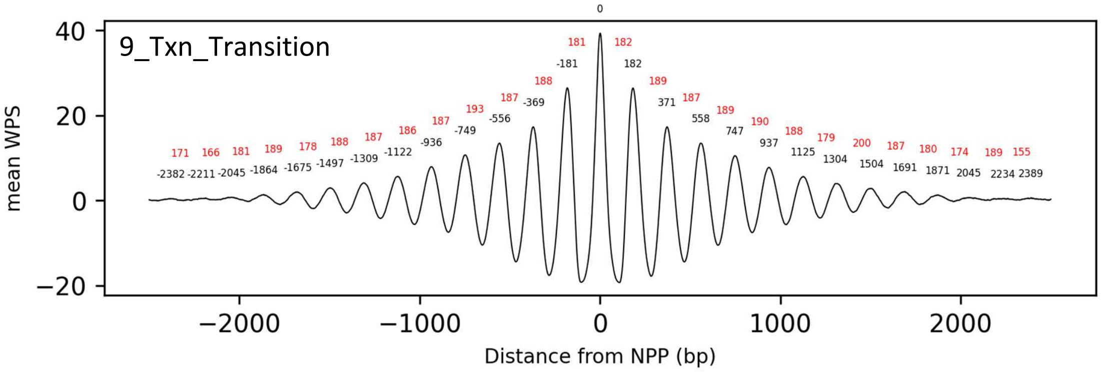
9_Txn_Transition
mean WPS
Distance from NPP (bp)
0
181 182
-181 182
188 189
187 -369 371 187
193 -556 558 189
187 -749 747 190
186 -936 937 188
187 -1122 1125 179
188 -1309 1304 200
178 -1497 1504 187
189 -1675 1691 180
181 -1864 1871 174
166 -2045 2045 189
171 -2211 2234 155
-2382 2389

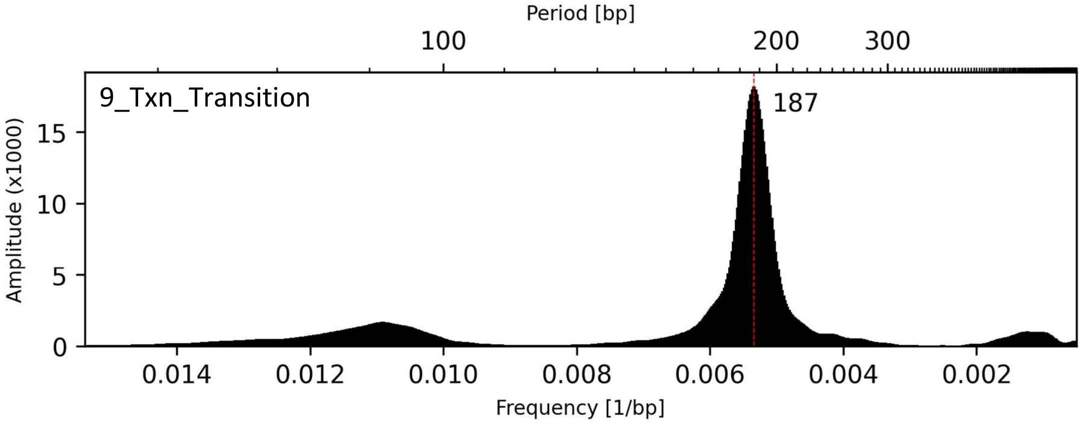
Period [bp]
100
200
300
9_Txn_Transition
187
Amplitude (x1000)
Frequency [1/bp]

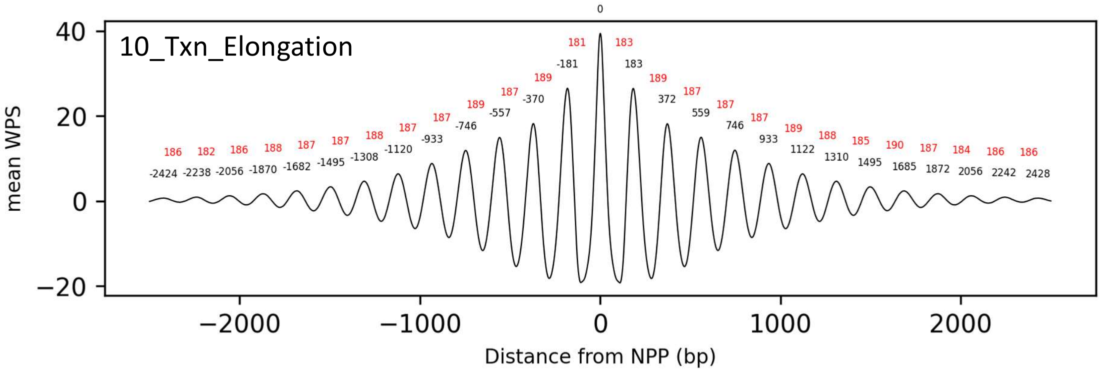
10_Txn_Elongation
mean WPS
Distance from NPP (bp)
0
181 183
-181 183
189 189
187 -370 372 187
189 -557 559 187
187 -746 746 187
187 -933 933 189
188 -1120 1122 188
187 -1308 1310 185
187 -1495 1495 190
188 -1682 1685 187
186 -1870 1872 184
182 -2056 2056 186
186 -2238 2242 186
-2424 2428

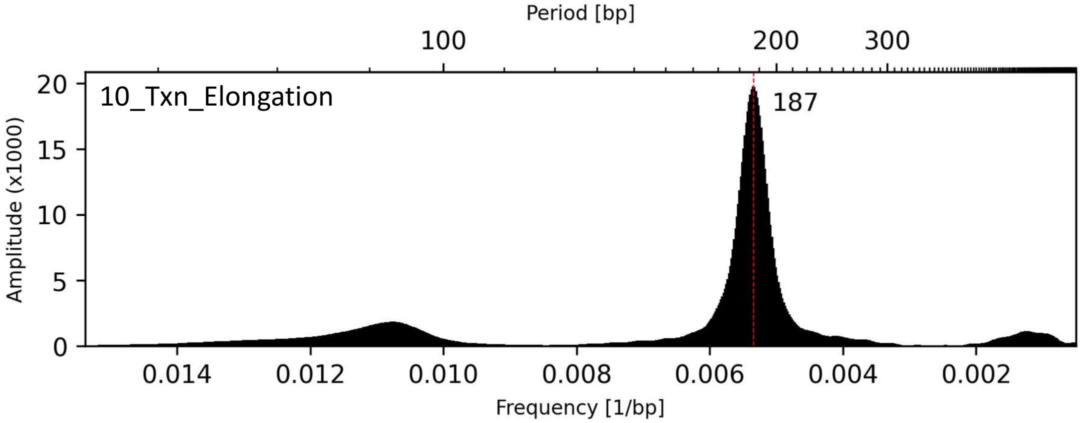
Period [bp]
100
200
300
10_Txn_Elongation
187
Amplitude (x1000)
Frequency [1/bp]

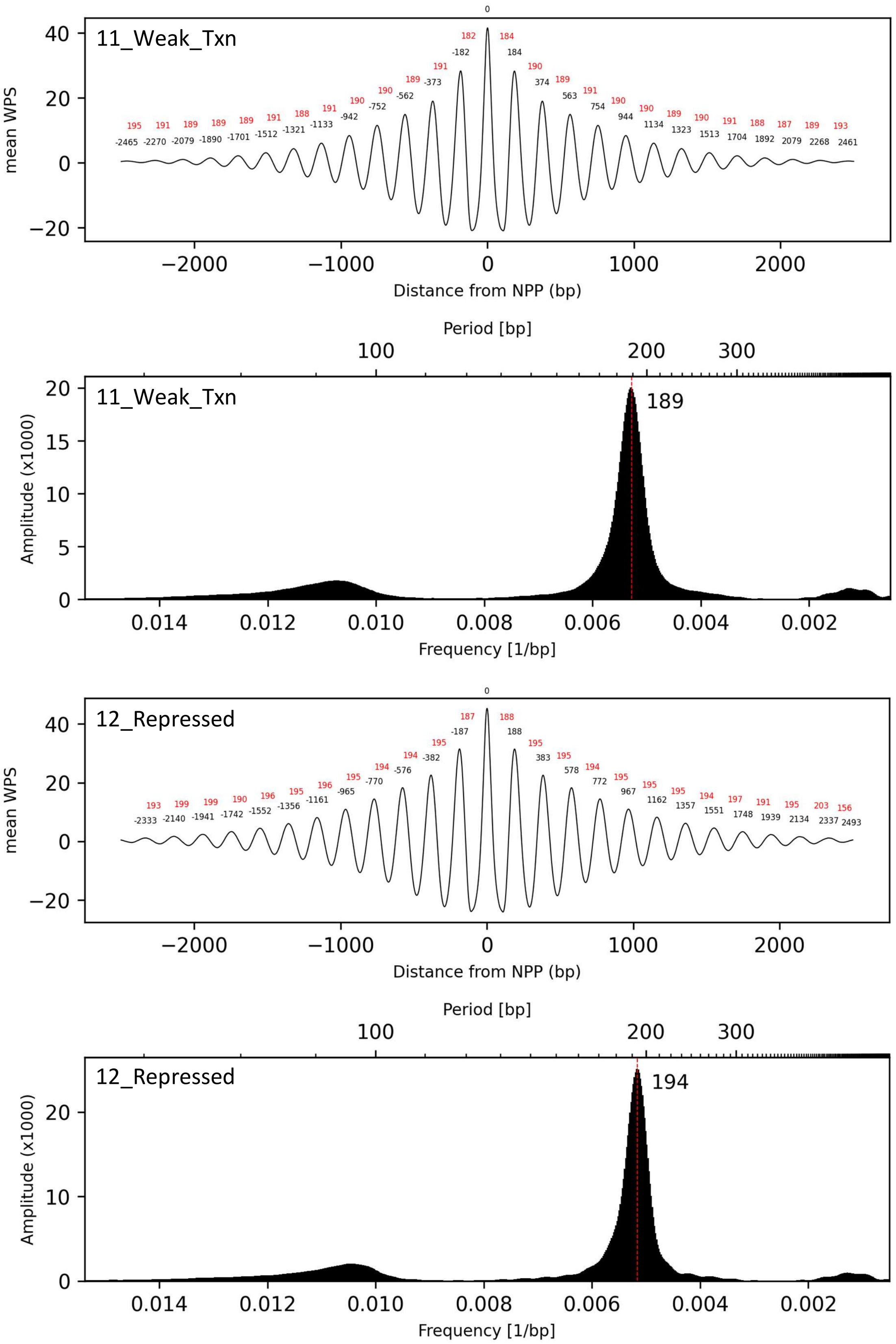
11_Weak_Txn
mean WPS
Distance from NPP (bp)
0
182 184
-182 184
191 190
-373 374
189 189
-562 563
190 191
-752 754
190 190
-942 944
191 190
-1133 1134
188 189
-1321 1323
191 190
-1512 1513
189 191
-1701 1704
189 188
-1890 1892
189 187
-2079 2079
191 189
-2270 2268
195 193
-2465 2461
Period [bp]
Amplitude (x1000)
Frequency [1/bp]
189
12_Repressed
mean WPS
Distance from NPP (bp)
0
187 188
-187 188
195 195
-382 383
194 195
-576 578
194 194
-770 772
195 195
-965 967
196 195
-1161 1162
195 195
-1356 1357
196 194
-1552 1551
190 197
-1742 1748
199 191
-1941 1939
199 195
-2140 2134
193 203
-2333 2337
156
2493
Period [bp]
Amplitude (x1000)
Frequency [1/bp]
194

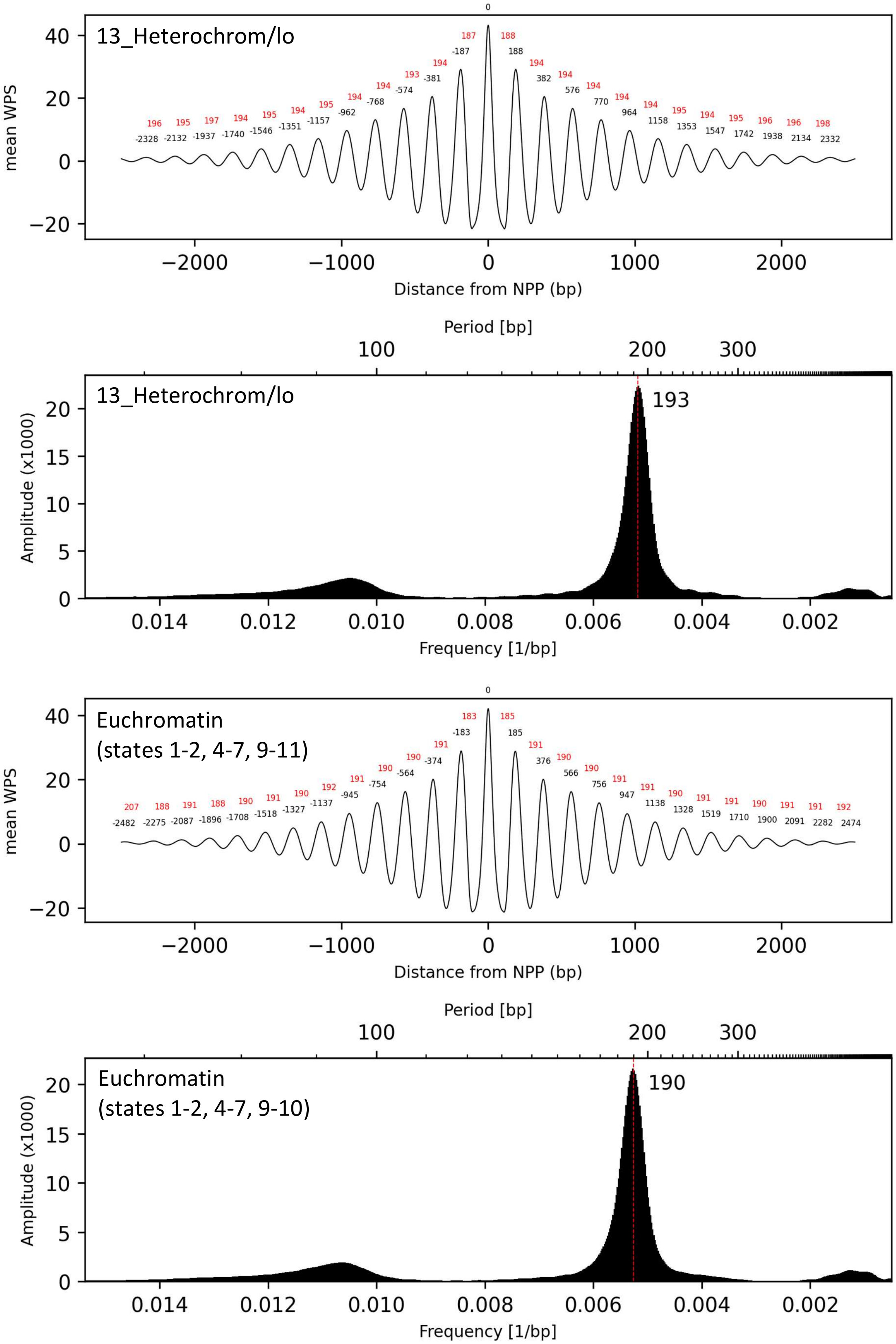
13_Heterochrom/lo
mean WPS
Distance from NPP (bp)
196 195 197 194 195 194 195 194 194 193 194 187 188 194 194 194 194 194 195 194 195 196 196 198
-2328 -2132 -1937 -1740 -1546 -1351 -1157 -962 -768 -574 -381 -187 0 188 382 576 770 964 1158 1353 1547 1742 1938 2134 2332
Period [bp]
13_Heterochrom/lo
193
Amplitude (x1000)
Frequency [1/bp]
Euchromatin
(states 1-2, 4-7, 9-11)
mean WPS
Distance from NPP (bp)
207 188 191 188 190 191 190 192 191 190 190 191 183 185 191 190 190 191 191 190 191 191 190 191 191 192
-2482 -2275 -2087 -1896 -1708 -1518 -1327 -1137 -945 -754 -564 -374 -183 0 185 376 566 756 947 1138 1328 1519 1710 1900 2091 2282 2474
Period [bp]
Euchromatin
(states 1-2, 4-7, 9-10)
190
Amplitude (x1000)
Frequency [1/bp]

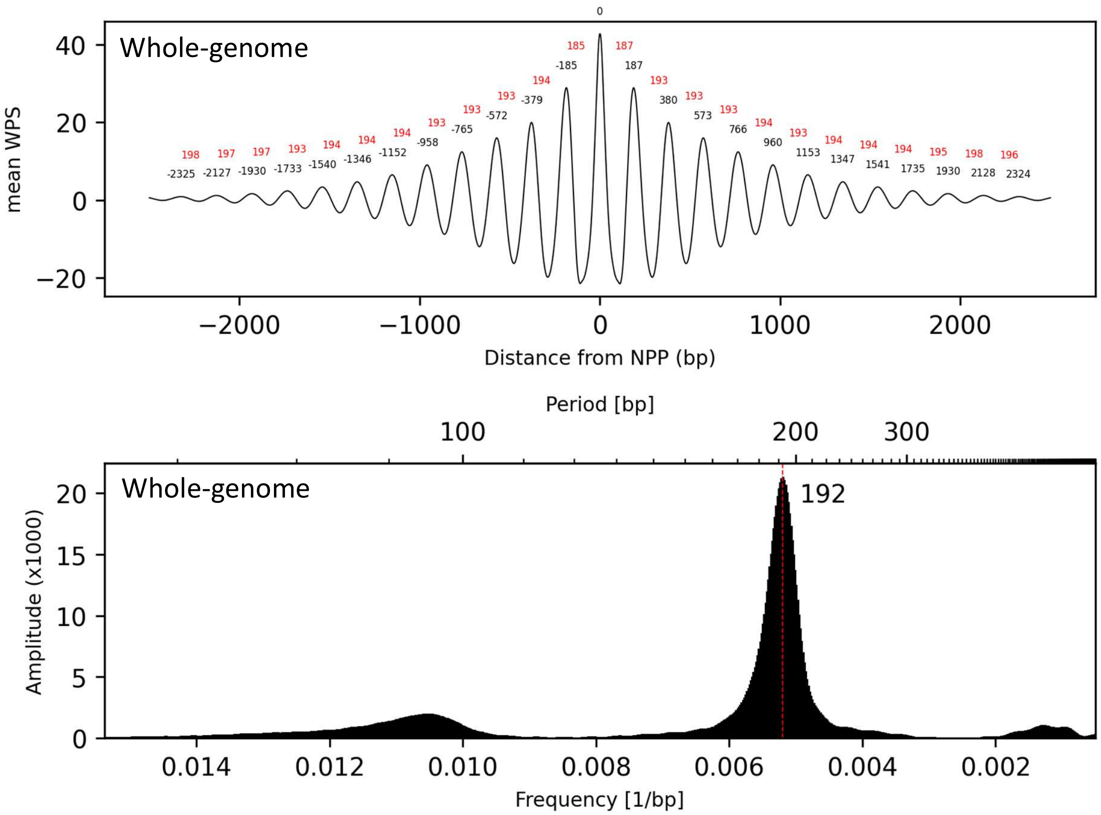


**Supplementary Figure S34. Fourier transforms of mean WPS aligned to NPPs within ChromHMM chromatin states**. Distances of each peak from position 0 are in black and distances between peaks are in red. The mode interpeak distance (period) are displayed for each Fourier transform. NPPs are those of the CH01 healthy sample pool from Snyder et. al. (2016)[10]. Chromatin states are those of Ernst et al. (2011)[20].
