## Supplementary Figures and Tables for "Unraveling cfDNA Fragmentation Patterns: Linking Chromatin Architecture to Cancer Diagnostic Sensitivity"

**Supplementary Table S1.** Sequenced sample information

| Sample/Pool | Cancer Type | No. of samples | Sex | Mean Coverage | Nuc peaks called |
|---|---|---|---|---|---|
| BRA01 | Brain | 13 | Mixed | 17 | 10,170,903 |
| COL01 | Colorectal | 19 | Mixed | 20 | 10,580,603 |
| IH02 | Healthy | 1 | Male | 21 | 9,573,485 |
| BRE01 | Breast | 16 | Female | 23 | 11,040,968 |
| IC17 | Liver | 1 | Male | 28 | 9,916,086 |
| IC37 | Colorectal | 1 | Female | 31 | 9,399,740 |
| BBC01 | BRE01, BRA01, COL01 | 48 | Mixed | 62 | 12,296,532 |
| BH01 | Healthy | Unknown | Mixed | 97 | 12,494,723 |
| IH01 | Healthy | 1 | Male | 104 | 11,894,427 |
| CH01 | IH01, IH02, BH01 | 2 + BH01 | Mixed | 231 | 12,811,998 |
| CA01 | CH01, Mix | 61 + BH01 | Mixed | 698 | 12,840,565 |

**Supplementary Table S2.1. ddPCR assays targeting loci with strong nucleosome protection peaks.**

| Midpoint (GRCh37) | Quadraplex | Position to nuc peak | Fluorophore | Amplitude |
|---|---|---|---|---|
| chr2:36639194 | 2 | Upstream | FAM | Low |
| chr2:36639274 | 3 | Centre | FAM | High |
| chr2:36639354 | 1 | Downsteam | FAM | High |
| chr3:104760321 | 3 | Upstream | FAM | Low |
| chr3:104760401 | 1 | Centre | FAM | Low |
| chr3:104760489 | 2 | Downsteam | FAM | High |
| chr5:10112285 | 1 | Upstream | HEX | Low |
| chr5:10112365 | 2 | Centre | HEX | High |
| chr5:10112445 | 3 | Downsteam | HEX | Low |
| chr6:112350854 | 2 | Upstream | HEX | Low |
| chr6:112350934 | 1 | Centre | HEX | High |
| chr6:112351014 | 3 | Downsteam | HEX | High |

| Name | Oligo | Sequence |
|---|---|---|
| chr2:36639194 Upstream | Forward | CTAGTGAAGCTGACCCTATTGAAT |
| | Reverse | GCTTTCTGACTTGTTGCTTTAGAA |
| | Probe | AGACAGTTTTCGACTTTATCTCTCATATTT |
| chr2:36639274 Centre | Forward | AAAGCAACAAGTCAGAAAGCTTGA |
| | Reverse | TTGGGCGTCATCACAGTTACTA |
| | Probe | CTAATCATGACCCGTAGCAGCTCT |
| chr2:36639354 Down | Forward | GTAACTGTGATGACGCCCAAC |
| | Reverse | GAAACTGCAAGTTAACAGCTACTC |
| | Probe | AAACATTCTCAGTAAATAACCAAGAACTCC |
| chr3:104760321 Upstream | Forward | TGGACTAGAAAATGACACCGAATCTG |
| | Reverse | TAGATGGCTTGTACACAGCAGAC |
| | Probe | TTCTGGAGGCTGAGAAGCCCAC |
| chr3:104760401 Centre | Forward | TGCTGTGTACAAGCCATCTAGT |
| | Reverse | TGAAGCCTCTGGAGATTAGATCAG |
| | Probe | ATGGTATTCTGTTACAGTAGCCTGAAAAG |
| chr3:104760489 Down | Forward | CCAGAGGCTTCAAATTGGTG |
| | Reverse | GATTAGGTAAATTGGAGACTAACTACATTG |
| | Probe | CTGAGTAAAGTCATGCAAACTCAATTATAG |
| chr5:10112285 Upstream | Forward | TCCCGCGTAATATCATCTCCAAC |
| | Reverse | TAGGCATGTAGCAAACCCAGG |
| | Probe | CCCATGGAATCATTTCCTTTATTGTAAAAT |
| chr5:10112365 Centre | Forward | CTGGGTTTGCTACATGCCTAATAC |
| | Reverse | AATGTTTTCTTCCAGCTGGTACTA |
| | Probe | AAGCCACTGGCTGGACAGAATG |
| chr5:10112445 Down | Forward | ACCAGCTGGAAGAAAACATTCA |
| | Reverse | GATGACACCATAGGTTGACCAT |
| | Probe | CCCGCCTTAAAGTTTAGGGATTAAACC |
| chr6:112350854 Upstream | Forward | TGCAAGGTGCCTGTAGTTCT |
| | Reverse | GAAGAATGTGCTGGTTTAAGGAGG |
| | Probe | TCTTCTGCTGAAGACTTTGCTTAGGA |
| chr6:112350934 Centre | Forward | CTTAAACCAGCACATTCTTCATCT |
| | Reverse | ACATAGAACGGGTAGAACATTTGC |

| | | |
|---|---|---|
| | Probe | TTAGCCGGTTTCTGAATCTTAAACAGTA |
| chr6:112351014 Down | Forward | ATGTTCTACCCGTTCTATGTCTTT |
| | Reverse | ACCTTAAATGCCATGATTCCTTG |
| | Probe | CCTAACTCTCAGATGCTTTCTAGTCTATC |

**Supplementary Table S2.2 ddPCR relative concentrations of positions relative to nucleosome peaks -ANOVA.**

| Sample | Mean relative frequencies ± standard deviation (%) | | | One-way ANOVA |
|---|---|---|---|---|
| | Upstream | Centre | Downstream | (p) |
| cfDNA - Breast Cancer | 6.56 ± 1.03 | 12.14 ± 1.61 | 6.3 ± 0.84 | < 0.001 |
| cfDNA - Colorectal Cancer | 6.72 ± 1.16 | 12.14 ± 1.75 | 6.14 ± 1.02 | < 0.001 |
| gDNA- Healthy | 8.38 ± 0.28 | 8.32 ± 0.29 | 8.3 ± 0.28 | 0.052 |
| gDNA -137 bp sonicated | 8.41 ± 0.88 | 8.69 ± 1.11 | 7.9 ± 0.87 | 0.139 |
| gDNA -162 bp sonicated | 8.31 ± 0.72 | 9.04 ± 0.92 | 7.65 ± 1.18 | 0.005 |
| gDNA - 186 bp sonicated | 8.6 ± 1.23 | 8.88 ± 1.24 | 7.52 ± 1.17 | 0.024 |
| gDNA - 213 bp sonicated | 8.22 ± 1.03 | 8.6 ± 0.83 | 8.19 ± 0.58 | 0.421 |

**Supplementary Table S2.3 ddPCR relative concentrations of quadraplexes - ANOVA.**

| Sample | Quadraplex 1 (2 centre assays) | Quadraplex 2 (2 upstream assays) | Quadraplex 3 (2 downstream assays) | One-way ANOVA (p) |
|---|---|---|---|---|
| cfDNA - Breast Cancer | 9.74 ± 3.19 | 7.72 ± 3.02 | 7.55 ± 1.99 | < 0.001 |
| cfDNA - Colorectal Cancer | 9.78 ± 3.25 | 7.69 ± 3.05 | 7.53 ± 2.11 | < 0.001 |
| gDNA- Healthy | 8.33 ± 0.29 | 8.47 ± 0.26 | 8.2 ± 0.23 | < 0.001 |
| gDNA -137 bp sonicated | 8.16 ± 0.74 | 8.48 ± 1.38 | 8.37 ± 0.78 | 0.73 |
| gDNA -162 bp sonicated | 8.4 ± 0.76 | 8.34 ± 1.60 | 8.26 ± 0.81 | 0.95 |
| gDNA - 186 bp sonicated | 8.12 ± 0.39 | 8.1 ± 1.96 | 8.78 ± 1.11 | 0.37 |
| gDNA - 213 bp sonicated | 8.35 ± 0.63 | 7.98 ± 1.09 | 8.67 ± 0.60 | 0.12 |

**Supplementary Table S3.** Twenty most common mutation hotspots in cancer.

| Gene | Residue | % of samples Chang (2018)[1] | Genomic location (GRCh37) | Nearest 5' nucleosome peak | | Nearest 3' nucleosome peak | |
|---|---|---|---|---|---|---|---|
| | | | | bp | WPS | bp | WPS |
| KRAS* | G12 | 8.84 | 12:25398284 | 77 | 41 | 85 | 57 |
| BRAF* | V600 | 3.65 | 7:140453136 | 122 | 45 | 54 | 22 |
| IDH1* | R132 | 3.11 | 2:209113112 | 115 | 83 | 77 | 107 |
| PIK3CA | H1047 | 2.63 | 3:178952084 | 264 | 57 | 135 | 54 |
| PIK3CA | E545 | 2.57 | 3:178936091 | 46 | 54 | 135 | 49 |
| TP53 | R273 | 2.48 | 17:7577120 | 71 | 78 | 110 | 63 |
| TP53 | R248 | 2.28 | 17:7577538 | 308 | 63 | 26 | 33 |
| NRAS* | Q61 | 1.72 | 1:115256528 | 95 | 70 | 104 | 89 |
| TP53 | R175 | 1.69 | 17:7578406 | 22 | 13 | 330 | 33 |
| PIK3CA | E542 | 1.51 | 3:178936082 | 37 | 54 | 144 | 49 |
| KRAS | G13 | 1.07 | 11:534286 | 99 | 21 | 636 | 18 |
| TP53 | R213 | 1.04 | 17:7578211 | 347 | 43 | 173 | 13 |
| TP53 | G245 | 0.98 | 17:7577547 | 317 | 63 | 17 | 33 |
| TP53 | R282 | 0.89 | 17:7577093 | 44 | 78 | 137 | 63 |
| KRAS | Q61 | 0.77 | 12:25380275 | 51 | 120 | 138 | 117 |
| PTEN | R130 | 0.68 | 10:89692904 | 204 | 20 | 7 | 59 |
| TP53 | Y220 | 0.62 | 17:7578189 | 325 | 43 | 195 | 13 |
| AKT1 | E17 | 0.61 | 14:105246551 | 68 | 40 | 113 | 22 |
| TP53 | H179 | 0.61 | 17:7578393 | 9 | 13 | 343 | 33 |
| EGFR | L858 | 0.59 | 7:55259514 | 28 | 72 | 163 | 72 |

* Hotspots used in assays

**Supplementary Table S4.** Cancer mutation hotspot ddPCR quadruplex assays.

| Hotspot | WPS of nearest protection peak | Max distance to peak (bp) | Assay Midpoint (GRCh37) | Quadruplex | Fluorophore | Amplitude |
|---|---|---|---|---|---|---|
| IDH1 (R132) | 107 | 148 | chr2:209113092 | 1 | HEX | Low |
| | | 50 | chr2:209113191 | 2 | HEX | High |
| NRAS (Q61) | 70 | 148 | chr1:115256533 | 2 | FAM | Low |
| | | 50 | chr1:115256434 | 1 | FAM | Low |
| KRAS (G12) | 41 | 130 | chr12:25398290 | 2 | HEX | Low |
| | | 50 | chr12:25398209 | 1 | HEX | High |
| BRAF (V600) | 22 | 116 | chr7:140453124 | 2 | FAM | High |
| | | 50 | chr7:140453191 | 1 | FAM | High |

| Name | Oligo | Sequence |
|---|---|---|
| BRAF (V600) Mutant | Forward | CATCCACAAAATGGATCCAGA |
| | Reverse | CCTCACAGTAAAAATAGGTGATTTTGG |
| | Probe | TTCAAACTGATGGGACCCACTCC |
| BRAF (V600) Nucleosome | Forward | GCTAGACCAAAATCACCTATTTTTACTG |
| | Reverse | TGATAGGAAAATGAGATCTACTGTTTTC |
| | Probe | TCTTCATGAAGAAATATATCTGAGGTGTAG |
| IDH1 (R132) Mutant | Forward | GGCCATGAAAAAAAAAACATGC |
| | Reverse | TGGATGGGTAAAACCTATCATCATA |
| | Probe | AACATGACTTACTTGATCCCCATAAGCAT |
| IDH1 (R132) Nucleosome | Forward | ACTCACAAGCCGGGGGATAT |
| | Reverse | ACAAATGTGGAAATCACCAAATG |
| | Probe | CTCTGAAGACCGTGCCACCCA |
| KRAS (G12) Mutant | Forward | GATTCTGAATTAGCTGTATCGTCAAG |
| | Reverse | TTATTATAAGGCCTGCTGAAAATG |
| | Probe | CTGAATATAAACTTGTGGTAGTTGGAGC |
| KRAS (G12) Nucleosome | Forward | TCAAAGAATGGTCCTGCACC |
| | Reverse | ATACAGCTAATTCAGAATCATTTTGTG |
| | Probe | TTTACCTCTATTGTTGGATCATATTCG |
| NRAS (Q61) Mutant | Forward | TTCGCCTGTCCTCATGTATTG |
| | Reverse | AGTGGTTATAGATGGTGAAACCTGTT |
| | Probe | CCAGCTGTATCCAGTATGTCCAAC |
| NRAS (Q61) Nucleosome | Forward | AAGATCATCCTTTCAGAGAAAATAATGC |
| | Reverse | AGGCTTCCTCTGTGTATTTGCC |
| | Probe | CTGTAGAGGTTAATATCCGCAAATGACTTG |

**Supplementary Table S5.** Difference in median WPS nucleosome peak spacing between ChromHMM heterochromatin and euchromatin

| Chromosome | Median spacing | | Difference | U-statistic | p-value |
|---|---|---|---|---|---|
| | 13_Heterochrom/lo | Euchromatin (1-2, 4-7, 9-11) | | | |
| chr1 | 192 | 188 | 4 | 88391036236 | 0 |
| chr2 | 192 | 188 | 4 | 88857144793 | 0 |
| chr3 | 192 | 189 | 3 | 59916559116 | 2.29E-310 |
| chr4 | 192 | 189 | 3 | 42085307496 | 6.17E-173 |
| chr5 | 192 | 189 | 3 | 46134621368 | 1.87E-248 |
| chr6 | 192 | 189 | 3 | 44393135901 | 5.55E-253 |
| chr7 | 192 | 188 | 4 | 37430812422 | 1.26E-222 |
| chr8 | 192 | 189 | 3 | 27678170243 | 4.55E-199 |
| chr9 | 193 | 188 | 5 | 22072789980 | 1.78E-275 |
| chr10 | 192 | 188 | 4 | 27130775138 | 3.53E-247 |
| chr11 | 192 | 188 | 4 | 26641253473 | 5.05E-277 |
| chr12 | 192 | 188 | 4 | 29212237723 | 7.29E-263 |
| chr13 | 192 | 189 | 3 | 11921354515 | 3.31E-86 |
| chr14 | 192 | 188 | 4 | 13569174218 | 2.91E-205 |
| chr15 | 192 | 188 | 4 | 12143779663 | 1.94E-217 |
| chr16 | 192 | 187 | 5 | 11779384128 | 3.48E-241 |
| chr17 | 191 | 187 | 4 | 12235093762 | 2.46E-228 |
| chr18 | 192 | 189 | 3 | 7378917671 | 1.19E-94 |
| chr19 | 191 | 186 | 5 | 6401963385 | 2.77E-164 |
| chr20 | 192 | 188 | 4 | 6061849628 | 4.68E-155 |
| chr21 | 192 | 188 | 4 | 1661910629 | 4.14E-51 |
| chr22 | 192 | 188 | 4 | 2403931285 | 3.16E-127 |
| chrX | 193 | 189 | 4 | 18483262562 | 5.01E-90 |
| whole-genome | 192 | 188 | 4 | 1.16419E+13 | 0 |

**Supplementary Table S6.1: Lower agreement of nucleosome position calls between sexes for euchromatin regions of chrX**

| Mann-Whitney U Tests | | | | | | |
|---|---|---|---|---|---|---|
| chromatin_state | median distance between nearest neighbor nucleosome peaks (bp) - male versus female | | difference | p_value | space occupied by chromatin state (Mb) | |
| | chrX | chr7 | | | chrX | chr7 |
| 1_Active_Promoter | 18 | 11 | 7 | 2.15E-75 | 0.72 | 1.14 |
| 2_Weak_Promoter | 16 | 11 | 5 | 1.11E-39 | 0.54 | 1.02 |
| 3_Poised_Promoter | 17 | 12 | 5 | 3.96E-06 | 0.05 | 0.21 |
| 4_Strong_Enhancer | 16 | 14 | 2 | 6.94E-05 | 0.38 | 1.06 |
| 5_Strong_Enhancer | 15 | 13 | 2 | 3.30E-06 | 0.65 | 1.18 |
| 6_Weak_Enhancer | 16 | 12 | 4 | 3.32E-27 | 0.76 | 1.42 |
| 7_Weak_Enhancer | 15 | 13 | 2 | 8.82E-16 | 1.48 | 2.97 |
| 8_Insulator | 8 | 7 | 1 | 2.88E-03 | 0.62 | 0.78 |
| 9_Txn_Transition | 18 | 14 | 4 | 2.50E-06 | 0.16 | 0.96 |
| 10_Txn_Elongation | 19 | 16 | 3 | 1.77E-35 | 1.52 | 7.74 |
| 11_Weak_Txn | 17 | 14 | 3 | 9.48E-132 | 9.66 | 17.28 |
| **12_Repressed** | **14** | **13** | **1** | **2.77E-06** | **1.08** | **3.97** |
| **13_Heterochrom/lo** | **15** | **14** | **1** | **3.51E-196** | **133.23** | **114.73** |
| **Euchromatin (1-2,4-7,9-11)** | **17** | **14** | **3** | **4.48E-242** | **15.88** | **34.77** |

**Supplementary Table S6.2: Difference of Spearman's rank-order correlations between males and females in average distance between nucleosomes flanking nearest-neighbor nucleosome peaks.**

| | Spearman's correlation (rho) | | Fisher's z-test | |
|---|---|---|---|---|
| **chromatin_state** | **chr7** | **chrX** | **z_diff** | **p_value** |
| 1_Active_Promoter | 0.40 | 0.27 | 6.13 | 9.06E-10 |
| 2_Weak_Promoter | 0.37 | 0.28 | 4.03 | 5.63E-05 |
| 3_Poised_Promoter | 0.43 | 0.18 | 3.42 | 6.19E-04 |
| 4_Strong_Enhancer | 0.33 | 0.29 | 1.43 | 1.52E-01 |
| 5_Strong_Enhancer | 0.32 | 0.29 | 1.53 | 1.27E-01 |
| 6_Weak_Enhancer | 0.36 | 0.28 | 3.64 | 2.71E-04 |
| 7_Weak_Enhancer | 0.34 | 0.31 | 2.26 | 2.41E-02 |
| 8_Insulator | 0.57 | 0.51 | 3.21 | 1.34E-03 |
| 9_Txn_Transition | 0.31 | 0.26 | 1.40 | 1.63E-01 |
| 10_Txn_Elongation | 0.26 | 0.22 | 3.23 | 1.25E-03 |
| 11_Weak_Txn | 0.29 | 0.24 | 8.18 | 2.22E-16 |
| 12_Repressed | 0.34 | 0.29 | 3.53 | 4.11E-04 |
| 13_Heterochrom/lo | 0.23 | 0.22 | 6.45 | 1.10E-10 |
| 14_Repetitive/CNV | 0.46 | 0.44 | 0.58 | 5.59E-01 |
| 15_Repetitive/CNV | 0.35 | 0.68 | -9.10 | 0.00E+00 |
| Euchromatin | 0.30 | 0.25 | 10.79 | 0.00E+00 |
| None | 0.40 | 0.28 | 1.25 | 2.12E-01 |

**Supplementary Table S6.3: Spearman's rank-order correlations between males and females in average distance between nucleosomes flanking nearest-neighbor nucleosome peaks.**

| chromosome | chromatin_state | correlation | p_value | n |
|---|---|---|---|---|
| chr7 | 1_Active_Promoter | 0.40 | 3.00E-186 | 4761 |
| | 2_Weak_Promoter | 0.37 | 1.23E-145 | 4402 |
| | 3_Poised_Promoter | 0.43 | 1.87E-42 | 921 |
| | 4_Strong_Enhancer | 0.33 | 5.20E-113 | 4501 |
| | 5_Strong_Enhancer | 0.32 | 2.79E-121 | 4937 |
| | 6_Weak_Enhancer | 0.36 | 1.14E-174 | 5799 |
| | 7_Weak_Enhancer | 0.34 | 0.00E+00 | 12286 |
| | 8_Insulator | 0.57 | 4.84E-295 | 3425 |
| | 9_Txn_Transition | 0.31 | 4.41E-93 | 4125 |
| | 10_Txn_Elongation | 0.26 | 0.00E+00 | 32893 |
| | 11_Weak_Txn | 0.29 | 0.00E+00 | 74663 |
| | 12_Repressed | 0.34 | 0.00E+00 | 16725 |
| | 13_Heterochrom/lo | 0.23 | 0.00E+00 | 610767 |
| | 14_Repetitive/CNV | 0.46 | 4.90E-80 | 1526 |

| | | | | |
|---|---|---|---|---|
| | 15_Repetitive/CNV | 0.35 | 1.12E-24 | 799 |
| | Euchromatin | 0.30 | 0.00E+00 | 148367 |
| | None | 0.40 | 1.18E-111 | 2864 |
| | 1_Active_Promoter | 0.27 | 1.35E-48 | 2748 |
| | 2_Weak_Promoter | 0.28 | 7.58E-40 | 2159 |
| | 3_Poised_Promoter | 0.18 | 1.17E-02 | 191 |
| | 4_Strong_Enhancer | 0.29 | 1.31E-30 | 1519 |
| | 5_Strong_Enhancer | 0.29 | 1.21E-52 | 2638 |
| | 6_Weak_Enhancer | 0.28 | 6.07E-55 | 2910 |
| | 7_Weak_Enhancer | 0.31 | 1.95E-127 | 5887 |
| | 9_Txn_Transition | 0.26 | 4.19E-11 | 643 |
| chrX | 8_Insulator | 0.51 | 2.18E-177 | 2656 |
| | 10_Txn_Elongation | 0.22 | 1.11E-67 | 6022 |
| | 11_Weak_Txn | 0.24 | 0.00E+00 | 40830 |
| | 12_Repressed | 0.29 | 2.84E-87 | 4446 |
| | 13_Heterochrom/lo | 0.22 | 0.00E+00 | 684121 |
| | 14_Repetitive/CNV | 0.44 | 4.47E-55 | 1139 |
| | 15_Repetitive/CNV | 0.68 | 1.73E-110 | 822 |
| | Euchromatin | 0.25 | 0.00E+00 | 65356 |
| | None | 0.28 | 6.15E-03 | 92 |

**Supplementary Table S7.1.** Difference in WPS median nucleosome peak spacing between ChromHMM hetero- and eu- chromatin (§contiguous)

| Sample | Sex | Δ median nucleosome peak spacing (Hetero – Eu) of primary peak (115-315bp) across chromosomes | | | | | | | | | | | | | | | | | | | | | | | |
|---|---|---|---|---|---|---|---|---|---|---|---|---|---|---|---|---|---|---|---|---|---|---|---|---|---|
| | | X | 7 | 8 | 9 | 6 | 10 | 11 | 12 | 5 | 4 | 13 | 3 | 14 | 15 | 16 | 17 | 18 | 2 | 20 | 1 | 19 | 21 | 22 | 1-22 |
| †BRA01 | f/m | 2 | 3 | 3 | 3 | 3 | 3 | 3 | 3 | 3 | 3 | 3 | 3 | 3 | 4 | 4 | 3 | 3 | 3 | 4 | 3 | 4 | 3 | 4 | 3 |
| †COL01 | f/m | 2 | 2 | 2 | 2 | 2 | 2 | 3 | 3 | 2 | 2 | 2 | 2 | 3 | 3 | 3 | 3 | 3 | 2 | 3 | 3 | 3 | 2 | 3 | 3 |
| †BRE01 | f | 1* | 2 | 3 | 2 | 2 | 2 | 3 | 3 | 2 | 2 | 2 | 2 | 3 | 3 | 3 | 4 | 3 | 2 | 3 | 3 | 3 | 3 | 3 | 3 |
| IC17 | m | 3 | 4 | 3 | 3 | 3 | 3 | 3 | 3 | 3 | 3 | 2 | 3 | 3 | 3 | 4 | 3 | 2 | 3 | 4 | 3 | 3 | 3 | 2 | 3 |
| IC37 | f | 2* | 3 | 3 | 3 | 3 | 2 | 3 | 3 | 2 | 2 | 3 | 2 | 3 | 3 | 3 | 3 | 2 | 3 | 3 | 3 | 3 | 2 | 3 | 3 |
| IH02 | m | 2 | 3 | 3 | 3 | 3 | 3 | 3 | 4 | 3 | 3 | 2 | 3 | 3 | 3 | 4 | 4 | 2 | 3 | 4 | 3 | 4 | 2 | 3 | 3 |
| †BBC01 | f/m | 3 | 3 | 3 | 4 | 3 | 4 | 4 | 3 | 3 | 3 | 2 | 4 | 4 | 4 | 5 | 4 | 3 | 3 | 5 | 3 | 4 | 3 | 3 | 4 |
| †BH01 | f/m | 4 | 4 | 3 | 4 | 3 | 4 | 4 | 4 | 3 | 3 | 3 | 3 | 4 | 4 | 5 | 4 | 3 | 4 | 5 | 4 | 5 | 4 | 5 | 4 |
| IH01 | m | 3 | 3 | 3 | 4 | 2 | 3 | 3 | 3 | 3 | 2 | 2 | 3 | 4 | 4 | 5 | 4 | 3 | 4 | 5 | 3 | 4 | 3 | 4 | 3 |
| †CA01♀ | f | 2* | 4 | 4 | 4 | 4 | 4 | 4 | 4 | 4 | 3 | 3 | 4 | 4 | 4 | 5 | 4 | 3 | 4 | 5 | 4 | 5 | 4 | 4 | 4 |
| †CA01♂ | m | 4 | 3 | 3 | 3 | 3 | 3 | 4 | 4 | 3 | 4 | 3 | 3 | 4 | 4 | 4 | 5 | 4 | 3 | 4 | 4 | 5 | 3 | 4 | 3 |
| †CH01 | f/m | 4 | 4 | 4 | 4 | 4 | 4 | 5 | 4 | 4 | 3 | 3 | 4 | 4 | 4 | 5 | 4 | 3 | 4 | 5 | 4 | 5 | 4 | 5 | 4 |
| †CA01 | f/m | 4 | 4 | 4 | 4 | 4 | 4 | 4 | 4 | 4 | 3 | 3 | 4 | 4 | 4 | 5 | 5 | 3 | 4 | 5 | 4 | 5 | 4 | 5 | 4 |

"Heterochromatin" is composed of ChromHMM states 12 and 13 and "Euchromatin" of states 1-3, 4-7, and 9-11 of the GM12878 cell line.

§Adjoining "euchromatin" states are treated as contiguous for NPP spacing calculations. Chromosomes are ordered by ascending difference in length from chrX.

*NRL difference between euchromatin and heterochromatin diminishes in female chrX. †Pool of samples

**Supplementary Table S7.2.** Difference in WPS median peak spacing between ChromHMM hetero- and eu- chromatin (excluding states 3 and 11, §contiguous)

| Sample | Sex | Δ median nucleosome peak spacing (Hetero – Eu) of primary peak (115-315bp) across chromosomes | | | | | | | | | | | | | | | | | | | | | | | |
|---|---|---|---|---|---|---|---|---|---|---|---|---|---|---|---|---|---|---|---|---|---|---|---|---|---|
| | | X | 7 | 8 | 9 | 6 | 10 | 11 | 12 | 5 | 4 | 13 | 3 | 14 | 15 | 16 | 17 | 18 | 2 | 20 | 1 | 19 | 21 | 22 | 1-22 |
| †BRA01 | f/m | 2 | 4 | 3 | 4 | 3 | 3 | 4 | 3 | 4 | 3 | 3 | 3 | 4 | 4 | 4 | 4 | 4 | 4 | 4 | 4 | 5 | 4 | 4 | 4 |
| †COL01 | f/m | 2 | 3 | 3 | 3 | 3 | 3 | 3 | 3 | 3 | 2 | 2 | 3 | 3 | 3 | 3 | 4 | 3 | 3 | 3 | 3 | 4 | 3 | 3 | 3 |
| †BRE01 | f | 1* | 3 | 4 | 3 | 3 | 3 | 4 | 3 | 2 | 3 | 2 | 3 | 3 | 3 | 3 | 4 | 3 | 3 | 4 | 3 | 4 | 3 | 4 | 3 |
| IC17 | m | 3 | 4 | 3 | 4 | 3 | 4 | 4 | 4 | 3 | 3 | 3 | 3 | 3 | 4 | 4 | 4 | 3 | 3 | 4 | 3 | 4 | 3 | 3 | 4 |
| IC37 | f | 1* | 3.5 | 3 | 3 | 3 | 3 | 3 | 3 | 3 | 3 | 3 | 3 | 3 | 3 | 4 | 4 | 3 | 3 | 4 | 3 | 3 | 2 | 4 | 3 |
| IH02 | m | 3 | 4 | 3 | 4 | 3 | 3 | 4 | 4 | 4 | 3 | 2 | 4 | 4 | 4 | 4 | 5 | 3 | 4 | 4 | 4 | 4 | 3 | 4 | 4 |
| †BBC01 | f/m | 3 | 4 | 4 | 4 | 4 | 4 | 5 | 3 | 4 | 3 | 3 | 4 | 4 | 5 | 5 | 5 | 4 | 4 | 5 | 4 | 5 | 3 | 4 | 4 |
| †BH01 | f/m | 4 | 4 | 4 | 4 | 4 | 4 | 5 | 5 | 4 | 4 | 4 | 4 | 5 | 5 | 6 | 5 | 3 | 4 | 5 | 5 | 6 | 4 | 6 | 5 |
| IH01 | m | 3 | 3 | 4 | 4 | 3 | 4 | 4 | 4 | 3 | 3 | 3 | 3 | 5 | 5 | 5 | 5 | 3 | 4 | 5 | 4 | 5 | 3 | 5 | 4 |
| †CA01♀ | f | 2* | 5 | 4 | 4 | 4 | 4 | 5 | 5 | 4 | 4 | 4 | 4 | 5 | 5 | 6 | 5 | 3 | 4 | 5 | 5 | 6 | 4 | 4 | 5 |
| †CA01♂ | m | 4 | 4 | 3 | 4 | 4 | 4 | 4 | 4 | 4 | 4 | 3 | 3 | 4 | 4 | 5 | 5 | 4 | 4 | 5 | 4 | 6 | 4 | 5 | 4 |
| †CH01 | f/m | 4 | 5 | 4 | 5 | 4 | 4 | 5 | 5 | 4 | 4 | 4 | 4 | 5 | 5 | 6 | 5 | 3 | 4 | 5 | 5 | 6 | 4 | 6 | 5 |
| †CA01 | f/m | 4 | 5 | 4 | 5 | 4 | 4 | 5 | 5 | 4 | 4 | 4 | 4 | 5 | 5 | 6 | 5 | 3 | 4 | 6 | 5 | 6 | 5 | 6 | 5 |

"Heterochromatin" is composed of ChromHMM states 12 and 13 and "Euchromatin" of states 1-2, 4-7, and 9-10 of the GM12878 cell line.

§Adjoining "euchromatin" states are treated as contiguous for NPP spacing calculations. Chromosomes are ordered by ascending difference in length from chrX.

*NRL difference between euchromatin and heterochromatin diminishes in female chrX. †Pool of samples

**Supplementary Table S7.3.** Difference in WPS median nucleosome peak spacing between ChromHMM hetero- and eu- chromatin (§separate)

| | | Δ median nucleosome peak spacing (Hetero – Eu) of primary peak (115-315bp) across chromosomes | | | | | | | | | | | | | | | | | | | | | | | |
|---|---|---|---|---|---|---|---|---|---|---|---|---|---|---|---|---|---|---|---|---|---|---|---|---|---|
| **Sample** | **Sex** | **X** | **7** | **8** | **9** | **6** | **10** | **11** | **12** | **5** | **4** | **13** | **3** | **14** | **15** | **16** | **17** | **18** | **2** | **20** | **1** | **19** | **21** | **22** | **1-22** |
| †BRA01 | f/m | 3 | 4 | 3 | 3 | 4 | 4 | 4 | 3 | 3 | 3 | 3 | 3 | 4 | 4 | 4 | 4 | 4 | 4 | 5 | 4 | 4 | 4 | 4 | 4 |
| †COL01 | f/m | 3 | 3 | 3 | 3 | 3 | 3 | 3 | 3 | 3 | 2 | 3 | 3 | 3 | 3 | 3 | 4 | 3 | 3 | 3 | 3 | 3 | 3 | 3 | 3 |
| †BRE01 | f | 1* | 3 | 4 | 3 | 3 | 3 | 4 | 3 | 2 | 3 | 2 | 3 | 3 | 3 | 4 | 4 | 4 | 3 | 4 | 3 | 4 | 3 | 3 | 3 |
| IC17 | m | 4 | 4 | 4 | 4 | 3 | 4 | 4 | 4 | 3 | 3 | 3 | 3 | 3 | 4 | 4 | 3 | 3 | 4 | 4 | 3 | 4 | 3 | 3 | 4 |
| IC37 | f | 2* | 4 | 4 | 3 | 3 | 3 | 3 | 3 | 3 | 3 | 3 | 3 | 3 | 3 | 4 | 4 | 3 | 3 | 4 | 3 | 3 | 2 | 4 | 3 |
| IH02 | m | 3 | 3 | 4 | 4 | 3 | 4 | 4 | 4 | 3 | 3 | 2 | 3 | 4 | 4 | 4 | 3 | 3 | 3 | 4 | 4 | 4 | 3 | 4 | 4 |
| †BBC01 | f/m | 3 | 3 | 4 | 4 | 4 | 4 | 4 | 3 | 4 | 3 | 3 | 4 | 4 | 4 | 5 | 4 | 3 | 4 | 5 | 3 | 5 | 3 | 4 | 4 |
| †BH01 | f/m | 4 | 4 | 4 | 4 | 4 | 4 | 4 | 4 | 4 | 3 | 3 | 4 | 4 | 4 | 5 | 4 | 3 | 4 | 5 | 4 | 5 | 4 | 5 | 4 |
| IH01 | m | 3 | 3 | 4 | 4 | 3 | 4 | 4 | 3 | 3 | 2 | 3 | 3 | 4 | 4 | 5 | 4 | 3 | 3 | 5 | 3 | 4 | 3 | 5 | 3 |
| †CA01♀ | f | 3* | 4 | 4 | 4 | 4 | 4 | 5 | 5 | 4 | 4 | 4 | 4 | 4 | 4 | 5 | 4 | 4 | 4 | 5 | 5 | 6 | 4 | 4 | 4 |
| †CA01♂ | m | 5 | 4 | 3 | 4 | 3 | 4 | 4 | 4 | 3 | 4 | 3 | 3 | 4 | 4 | 5 | 5 | 4 | 3 | 5 | 4 | 5 | 4 | 4 | 4 |
| †CH01 | f/m | 4 | 4 | 4 | 4 | 4 | 4 | 5 | 4 | 4 | 3 | 3 | 4 | 5 | 4 | 5 | 5 | 3 | 4 | 5 | 4 | 5 | 4 | 5 | 4 |
| †CA01 | f/m | 4 | 4 | 4 | 4 | 4 | 4 | 5 | 4 | 4 | 4 | 3 | 4 | 5 | 5 | 6 | 5 | 3 | 4 | 5 | 5 | 5 | 4 | 5 | 4 |

"Heterochromatin" is composed of ChromHMM states 12 and 13 and "Euchromatin" of states 1-3, 4-7, and 9-11 of the GM12878 cell line.

§Adjoining "euchromatin" states are treated separately for NPP spacing calculations. Chromosomes are ordered by ascending difference in length from chrX.

*NRL difference between euchromatin and heterochromatin diminishes in female chrX. †Pool of samples

**Supplementary Table S7.4.** Difference in WPS median peak spacing between ChromHMM hetero- and eu- chromatin (excluding states 3 and 11, §separate)

| | | Δ median nucleosome peak spacing (Hetero – Eu) of primary peak (115-315bp) across chromosomes | | | | | | | | | | | | | | | | | | | | | | | |
|---|---|---|---|---|---|---|---|---|---|---|---|---|---|---|---|---|---|---|---|---|---|---|---|---|---|
| **Sample** | **Sex** | **X** | **7** | **8** | **9** | **6** | **10** | **11** | **12** | **5** | **4** | **13** | **3** | **14** | **15** | **16** | **17** | **18** | **2** | **20** | **1** | **19** | **21** | **22** | **1-22** |
| †BRA01 | f/m | 3 | 4 | 4 | 4 | 4 | 4 | 5 | 4 | 4 | 4 | 4 | 4 | 5 | 5 | 5 | 4 | 4 | 4 | 5 | 5 | 5 | 4 | 5 | 4 |
| †COL01 | f/m | 3 | 4 | 4 | 3 | 3 | 3 | 4 | 4 | 3 | 3 | 3 | 3 | 4 | 4 | 4 | 5 | 4 | 3 | 4 | 4 | 4 | 3 | 4 | 4 |
| †BRE01 | f | 2* | 3 | 5 | 3 | 3 | 3 | 4 | 3 | 3 | 3 | 3 | 3 | 4 | 3 | 4 | 4 | 4 | 3 | 4 | 4 | 4 | 4 | 4 | 4 |
| IC17 | m | 4 | 5 | 5 | 5 | 4 | 4 | 5 | 4 | 4 | 4 | 4 | 4 | 3 | 4 | 5 | 4 | 4 | 4 | 5 | 4 | 5 | 4 | 3 | 4 |
| IC37 | f | 2* | 4 | 4 | 4 | 4 | 3 | 4 | 4 | 4 | 4 | 4 | 4 | 4 | 4 | 4 | 4 | 4 | 4 | 4 | 4 | 4 | 3 | 5 | 4 |
| IH02 | m | 4 | 4 | 4 | 4 | 4 | 4 | 5 | 5 | 4 | 4 | 3 | 4 | 5 | 5 | 5 | 4 | 4 | 4 | 5 | 4 | 5 | 3 | 5 | 4 |
| †BBC01 | f/m | 3 | 4 | 5 | 5 | 4 | 5 | 5 | 4 | 4 | 4 | 3 | 4 | 5 | 5 | 6 | 5 | 4 | 4 | 5 | 4 | 5 | 4 | 4 | 5 |
| †BH01 | f/m | 4 | 5 | 5 | 5 | 4 | 5 | 5 | 5 | 5 | 4 | 4 | 4 | 5 | 5 | 6 | 5 | 4 | 5 | 6 | 5 | 6 | 5 | 6 | 5 |
| IH01 | m | 4 | 4 | 5 | 5 | 4 | 4 | 4 | 4 | 4 | 3 | 4 | 4 | 5 | 5 | 6 | 5 | 4 | 4 | 6 | 4 | 5 | 4 | 6 | 4 |
| †CA01♀ | f | 3* | 5 | 5 | 5 | 5 | 5 | 5 | 5 | 5 | 4 | 4 | 5 | 5 | 5 | 6 | 5 | 4 | 5 | 6 | 5 | 6 | 5 | 5 | 5 |
| †CA01♂ | m | 5 | 4 | 4 | 4 | 4 | 4 | 5 | 4 | 4 | 4 | 4 | 4 | 4 | 5 | 5 | 6 | 4 | 4 | 5 | 5 | 6 | 4 | 5 | 4 |
| †CH01 | f/m | 5 | 5 | 5 | 5 | 5 | 5 | 6 | 5 | 5 | 4 | 4 | 5 | 5 | 5 | 6 | 5 | 4 | 5 | 6 | 5 | 6 | 5 | 6 | 5 |
| †CA01 | f/m | 5 | 5 | 5 | 5 | 5 | 5 | 5 | 5 | 5 | 4 | 4 | 4 | 5 | 5 | 6 | 6 | 4 | 5 | 6 | 5 | 6 | 5 | 6 | 5 |

"Heterochromatin" is composed of ChromHMM states 12 and 13 and "Euchromatin" of states 1-2, 4-7, and 9-10 of the GM12878 cell line.

§Adjoining "euchromatin" states are treated separately for NPP spacing calculations. Chromosomes are ordered by ascending difference in length from chrX.

*NRL difference between euchromatin and heterochromatin diminishes in female chrX. †Pool of samples

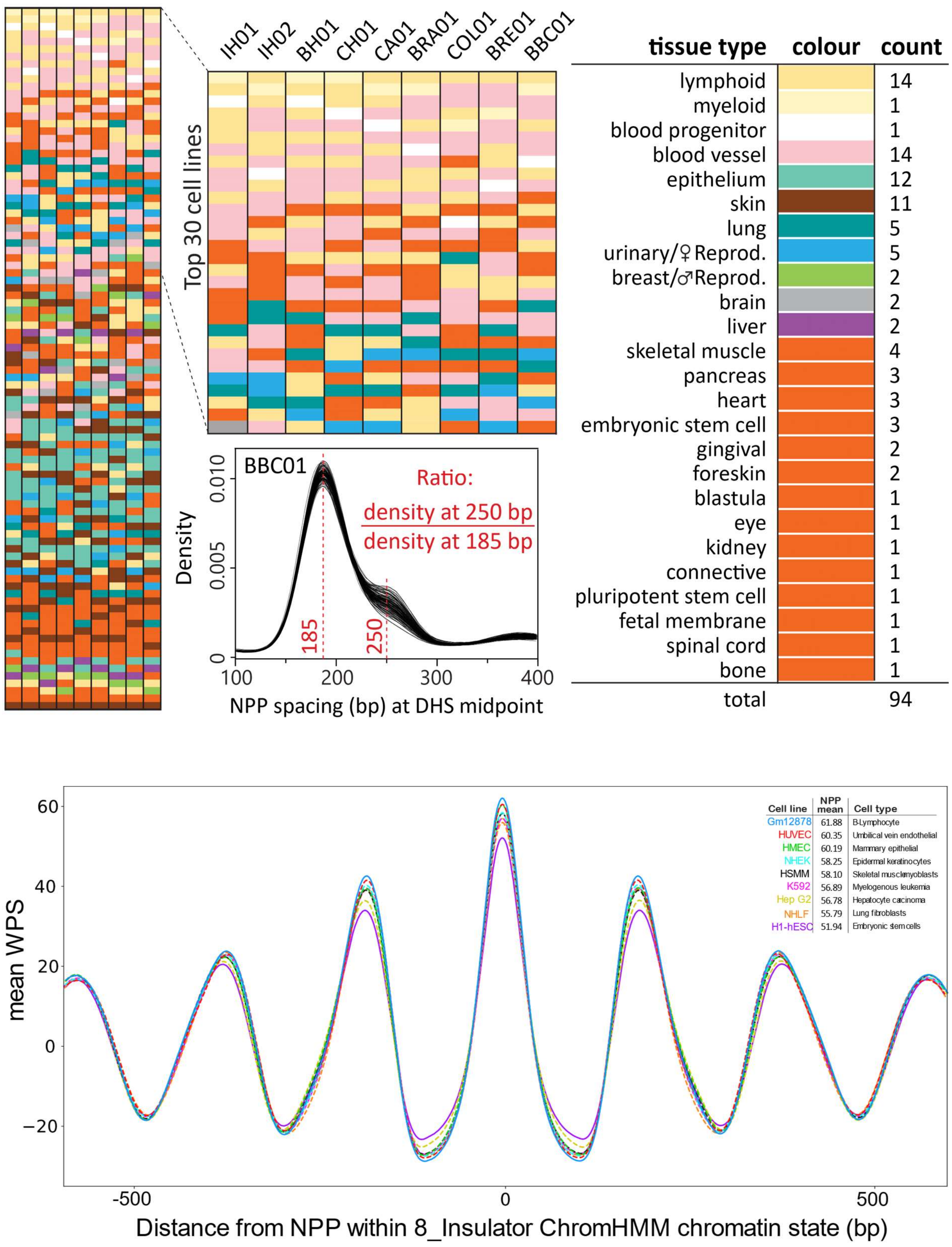


**Supplementary Figure S1. Use of increased nucleosome spacing to infer tissues of origin for cells contributing to cfDNA of samples.**

**(A)** Ratios of DHSs with 250 bp nucleosome spacing to DHSs with 185 bp spacing are ordered from highest to lowest for each sample. These ratios were determined using Savitzky-Golay smoothed (window size = 25, polynomial of order = 3) spacing distributions of nucleosome protection peaks (NPPs) flanking DHS midpoints for 94 cell lines, as depicted in the BBC01 example. A higher ratio indicates that the nucleosome spacing pattern in the cfDNA more closely resembles that of the cell line's DHSs, suggesting that the cfDNA likely originates from cells of that specific lineage. DHSs have a strong correlation with TF binding at regulatory regions, encompassing promoters, enhancers, and insulators. TF binding at these loci is linked to changes in nucleosome positioning, causing a shift from the baseline mode spacing of ≈185 bp to a mode of ≈250 bp. We used the DHS data from 94 cell lines not derived from cancer, sourced from The Encyclopedia of DNA Elements (ENCODE)[2,3]. Our findings revealed that the DHS nucleosome spacing footprints of all samples predominantly aligned with blood-related cell lines.

**(B)** Mean windowed protection score (WPS) relative to distance from nucleosome protection peaks (NPPs) within 8_Insulator chromatin states of 9 ENCODE cell lines**.** NPPs are those of the CH01 healthy sample pool from Snyder et. al. (2016)[4]. 8_Insulator chromatin states are those of Ernst et al. (2011)[5]. Stronger windowed protection scores within CTCF-bound insulators indicates a greater correlation of a cell line with the cells contributing cfDNA to the CH01 sample pool.

**A** Ten most common mutation hotspots in cancer

| Gene | Residue | % of samples Chang (2018) | Genomic location (GRCh37) | Nearest nucleosome peak Distance (bp) 5′ | Nearest nucleosome peak Distance (bp) 3′ | Nearest nucleosome peak Prominence score 5′ | Nearest nucleosome peak Prominence score 3′ |
|---|---|---|---|---|---|---|---|
| **KRAS*** | G12 | 8.84 | **12:25398284** | **77** | 85 | **41** | 57 |
| **BRAF*** | V600 | 3.65 | **7:140453136** | 122 | **54** | 45 | **22** |
| **IDH1*** | R132 | 3.11 | **2:209113112** | 115 | **77** | 83 | **107** |
| PIK3CA | H1047 | 2.63 | 3:178952084 | 264 | 135 | 57 | 54 |
| PIK3CA | E545 | 2.57 | 3:178936091 | 46 | 135 | 54 | 49 |
| TP53 | R273 | 2.48 | 17:7577120 | 71 | 110 | 78 | 63 |
| TP53 | R248 | 2.28 | 17:7577538 | 308 | 26 | 63 | 33 |
| **NRAS*** | Q61 | 1.72 | **1:115256528** | **95** | 104 | **70** | 89 |
| TP53 | R175 | 1.69 | 17:7578406 | 22 | 330 | 13 | 33 |
| PIK3CA | E542 | 1.51 | 3:178936082 | 37 | 144 | 54 | 49 |

* Hotspots and nearest peaks used as assay targets are in **bold**

Colour scale: Distance 0 to ≥ 90; Prominence score 0 to ≥ 110

**B**

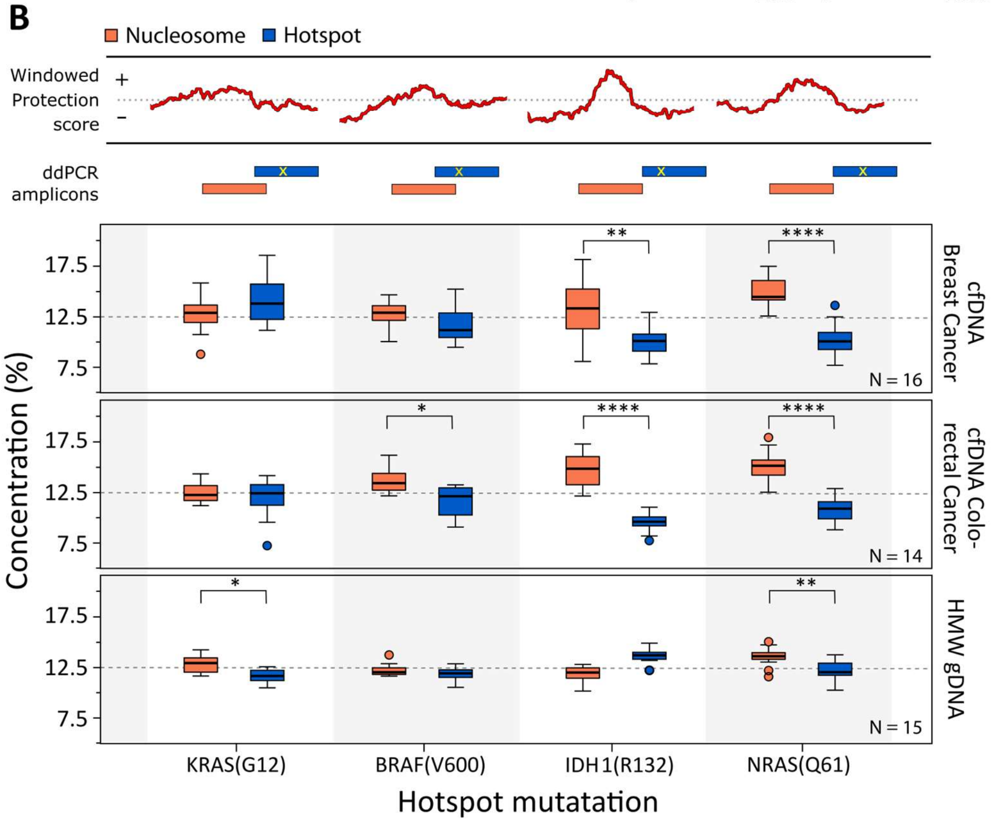

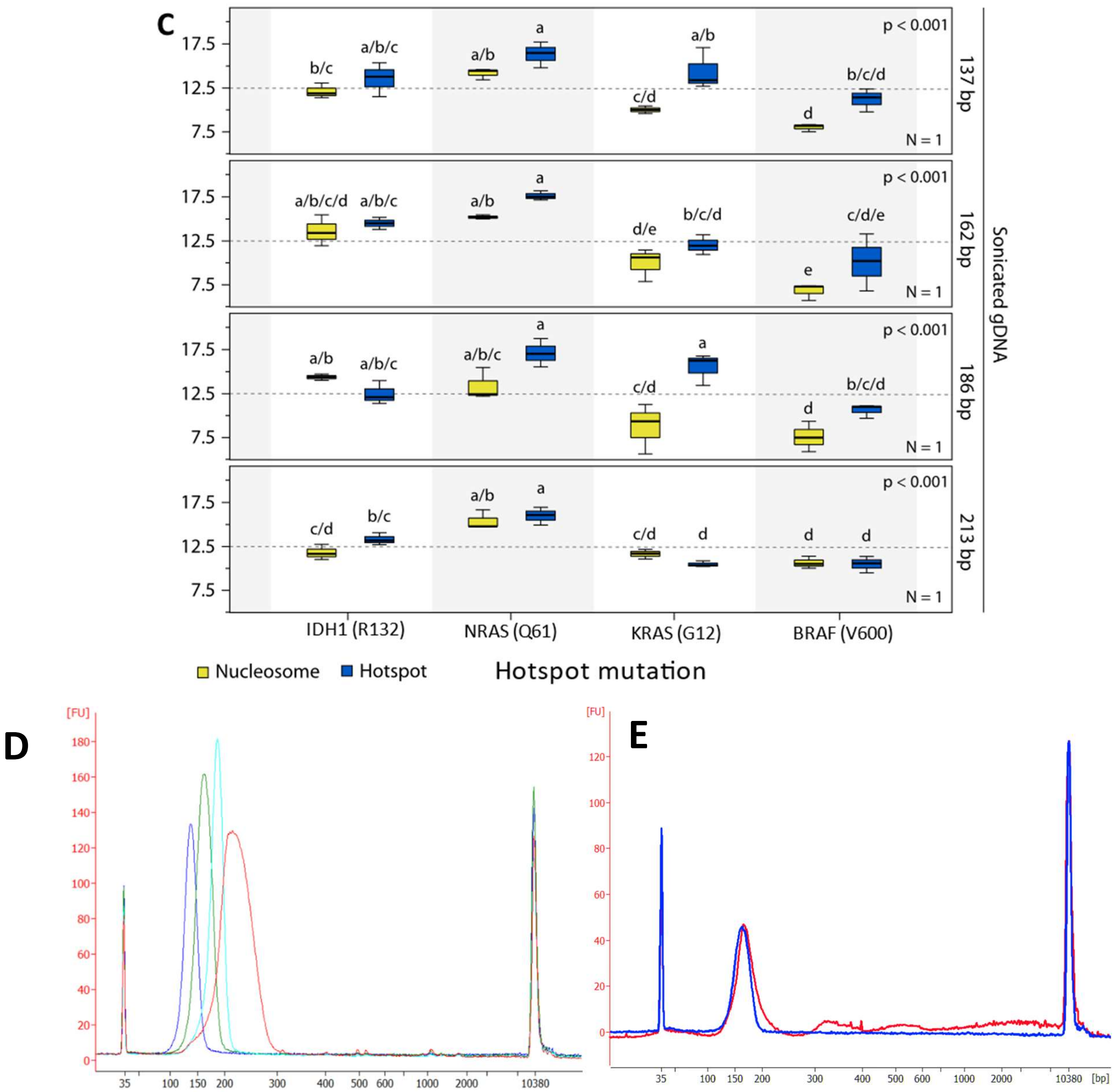


**Figure S2. PCR assays for somatic mutation analysis have reduced sensitivity near positionally-conserved nucleosome breakpoints.** (**A**) The top 10 most common somatic mutations among 24,592 cancers, including the distances to and strengths of the nearest upstream and downstream nucleosome positions. (**B**) Multiplexed ddPCR assays were designed to target two somatic mutations with low nucleosome protection scores (KRAS and BRAF), and two with high nucleosome protection scores (IDH1 and NRAS), together with a second proximal ddPCR assay centered on the predicted nucleosome binding site. n=30 cfDNA samples from breast and colorectal cancer were analyzed. Plots show the DNA concentrations determined for each sample as a percentage across all 8 assays, which controls for differences in sample input amounts, with the expected concentration under the null hypothesis indicated at 12.5% (1:8). Outliers are represented by circles. Statistical significance for each region within the samples was determined using one-sided Mann-Whitney U tests, with levels of significance denoted by *p < 0.05, **p < 0.01, ***p < 0.001, ****p < 0.0001. Control experiments with high molecular weight genomic DNA (HMW gDNA) stochastically fragmented to various sizes failed to reach the level of significance observed for patient cfDNA samples, except for the KRAs control. (**C**) gDNA samples sonicated, sized selected, and gel purified to create size distributions similar to cfDNA sample primary peaks, with mode fragment lengths displayed for each. These fragmented samples were run in three technical replicates. Letters above or below box and whisker plots represent homogenous subsets determined by post-hoc Tukey's HSD analyses (α=0.05) of one-way ANOVAs (p values on plots). (**D**) Bioanalyzer 2100 fragmentat sized profiles of sonicated gDNA sample size selected to produce primary distributions similar to cfDNA. Peaks at 137, 162, 186 and 213 bp. Electropherograms from 2100 Expert version B.02.10.SI764 running Agilent 2100 Bioanalyzer with a High Sensitivity DNA Chip. (**E**) Sonicated sample with 162 bp peak (blue) compared to typical cfDNA sample with 167 bp peak (red).

A

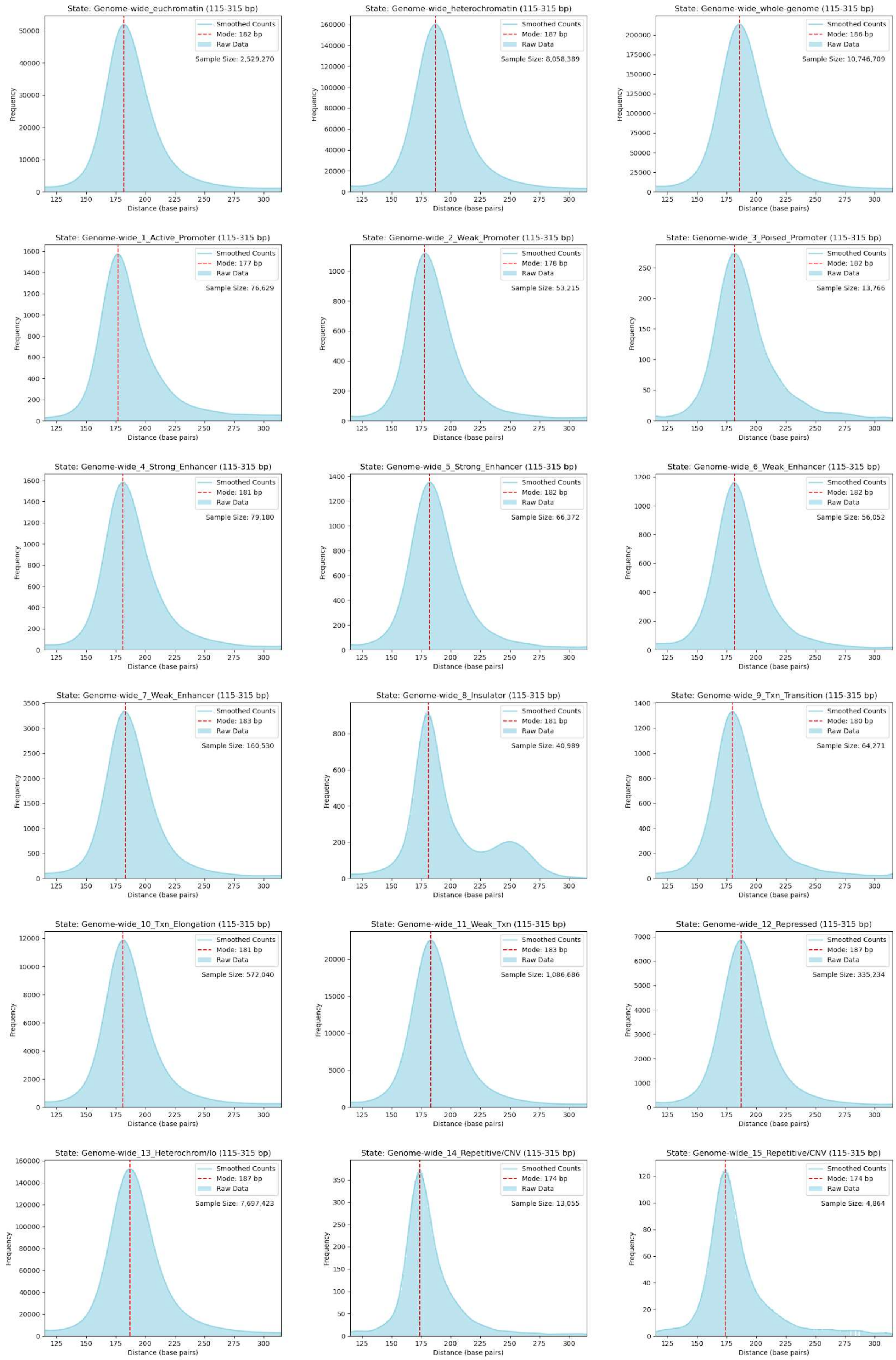
State: Genome-wide_euchromatin (115-315 bp)
Smoothed Counts
Mode: 182 bp
Raw Data
Sample Size: 2,529,270
State: Genome-wide_heterochromatin (115-315 bp)
Mode: 187 bp
Sample Size: 8,058,389
State: Genome-wide_whole-genome (115-315 bp)
Mode: 186 bp
Sample Size: 10,746,709
State: Genome-wide_1_Active_Promoter (115-315 bp)
Mode: 177 bp
Sample Size: 76,629
State: Genome-wide_2_Weak_Promoter (115-315 bp)
Mode: 178 bp
Sample Size: 53,215
State: Genome-wide_3_Poised_Promoter (115-315 bp)
Mode: 182 bp
Sample Size: 13,766
State: Genome-wide_4_Strong_Enhancer (115-315 bp)
Mode: 181 bp
Sample Size: 79,180
State: Genome-wide_5_Strong_Enhancer (115-315 bp)
Mode: 182 bp
Sample Size: 66,372
State: Genome-wide_6_Weak_Enhancer (115-315 bp)
Mode: 182 bp
Sample Size: 56,052
State: Genome-wide_7_Weak_Enhancer (115-315 bp)
Mode: 183 bp
Sample Size: 160,530
State: Genome-wide_8_Insulator (115-315 bp)
Mode: 181 bp
Sample Size: 40,989
State: Genome-wide_9_Txn_Transition (115-315 bp)
Mode: 180 bp
Sample Size: 64,271
State: Genome-wide_10_Txn_Elongation (115-315 bp)
Mode: 181 bp
Sample Size: 572,040
State: Genome-wide_11_Weak_Txn (115-315 bp)
Mode: 183 bp
Sample Size: 1,086,686
State: Genome-wide_12_Repressed (115-315 bp)
Mode: 187 bp
Sample Size: 335,234
State: Genome-wide_13_Heterochrom/lo (115-315 bp)
Mode: 187 bp
Sample Size: 7,697,423
State: Genome-wide_14_Repetitive/CNV (115-315 bp)
Mode: 174 bp
Sample Size: 13,055
State: Genome-wide_15_Repetitive/CNV (115-315 bp)
Mode: 174 bp
Sample Size: 4,864
Frequency
Distance (base pairs)

B

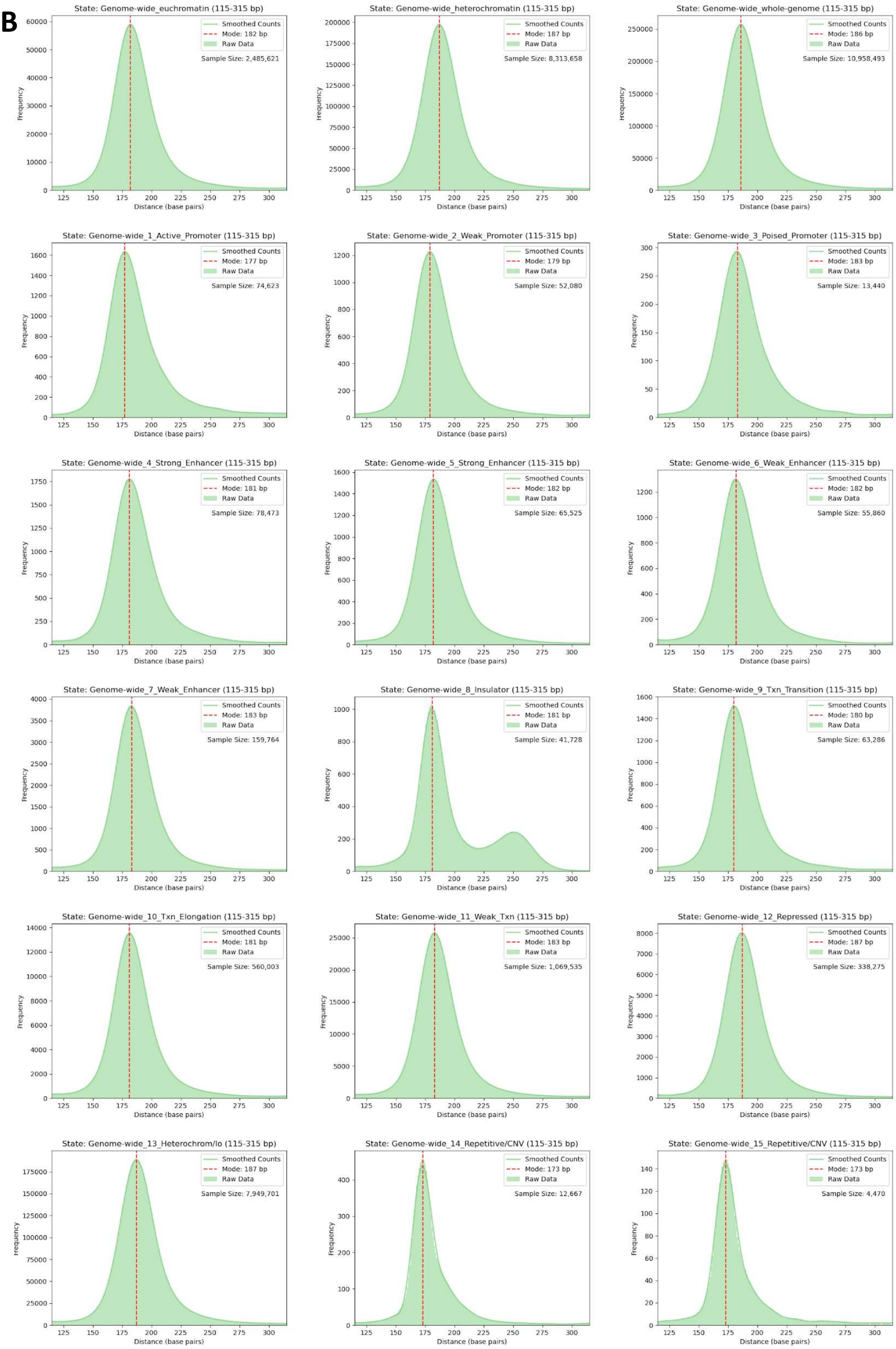
State: Genome-wide_euchromatin (115-315 bp)
Smoothed Counts
Mode: 182 bp
Raw Data
Sample Size: 2,485,621
State: Genome-wide_heterochromatin (115-315 bp)
Smoothed Counts
Mode: 187 bp
Raw Data
Sample Size: 8,313,658
State: Genome-wide_whole-genome (115-315 bp)
Smoothed Counts
Mode: 186 bp
Raw Data
Sample Size: 10,958,493
State: Genome-wide_1_Active_Promoter (115-315 bp)
Smoothed Counts
Mode: 177 bp
Raw Data
Sample Size: 74,623
State: Genome-wide_2_Weak_Promoter (115-315 bp)
Smoothed Counts
Mode: 179 bp
Raw Data
Sample Size: 52,080
State: Genome-wide_3_Poised_Promoter (115-315 bp)
Smoothed Counts
Mode: 183 bp
Raw Data
Sample Size: 13,440
State: Genome-wide_4_Strong_Enhancer (115-315 bp)
Smoothed Counts
Mode: 181 bp
Raw Data
Sample Size: 78,473
State: Genome-wide_5_Strong_Enhancer (115-315 bp)
Smoothed Counts
Mode: 182 bp
Raw Data
Sample Size: 65,525
State: Genome-wide_6_Weak_Enhancer (115-315 bp)
Smoothed Counts
Mode: 182 bp
Raw Data
Sample Size: 55,860
State: Genome-wide_7_Weak_Enhancer (115-315 bp)
Smoothed Counts
Mode: 183 bp
Raw Data
Sample Size: 159,764
State: Genome-wide_8_Insulator (115-315 bp)
Smoothed Counts
Mode: 181 bp
Raw Data
Sample Size: 41,728
State: Genome-wide_9_Txn_Transition (115-315 bp)
Smoothed Counts
Mode: 180 bp
Raw Data
Sample Size: 63,286
State: Genome-wide_10_Txn_Elongation (115-315 bp)
Smoothed Counts
Mode: 181 bp
Raw Data
Sample Size: 560,003
State: Genome-wide_11_Weak_Txn (115-315 bp)
Smoothed Counts
Mode: 183 bp
Raw Data
Sample Size: 1,069,535
State: Genome-wide_12_Repressed (115-315 bp)
Smoothed Counts
Mode: 187 bp
Raw Data
Sample Size: 338,275
State: Genome-wide_13_Heterochrom/lo (115-315 bp)
Smoothed Counts
Mode: 187 bp
Raw Data
Sample Size: 7,949,701
State: Genome-wide_14_Repetitive/CNV (115-315 bp)
Smoothed Counts
Mode: 173 bp
Raw Data
Sample Size: 12,667
State: Genome-wide_15_Repetitive/CNV (115-315 bp)
Smoothed Counts
Mode: 173 bp
Raw Data
Sample Size: 4,470
Frequency
Distance (base pairs)

**C**

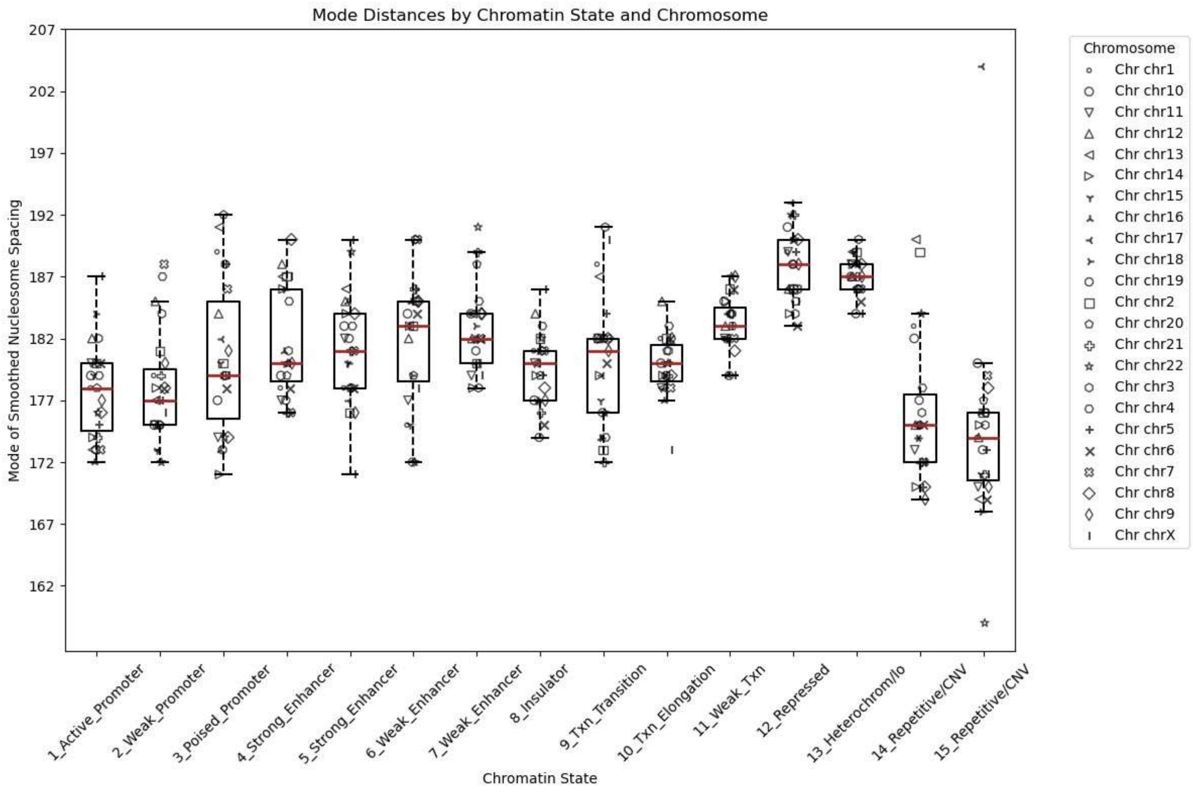
Mode Distances by Chromatin State and Chromosome
Mode of Smoothed Nucleosome Spacing
207
202
197
192
187
182
177
172
167
162
1_Active_Promoter
2_Weak_Promoter
3_Poised_Promoter
4_Strong_Enhancer
5_Strong_Enhancer
6_Weak_Enhancer
7_Weak_Enhancer
8_Insulator
9_Txn_Transition
10_Txn_Elongation
11_Weak_Txn
12_Repressed
13_Heterochrom/lo
14_Repetitive/CNV
15_Repetitive/CNV
Chromatin State
Chromosome
Chr chr1
Chr chr10
Chr chr11
Chr chr12
Chr chr13
Chr chr14
Chr chr15
Chr chr16
Chr chr17
Chr chr18
Chr chr19
Chr chr2
Chr chr20
Chr chr21
Chr chr22
Chr chr3
Chr chr4
Chr chr5
Chr chr6
Chr chr7
Chr chr8
Chr chr9
Chr chrX

**D**

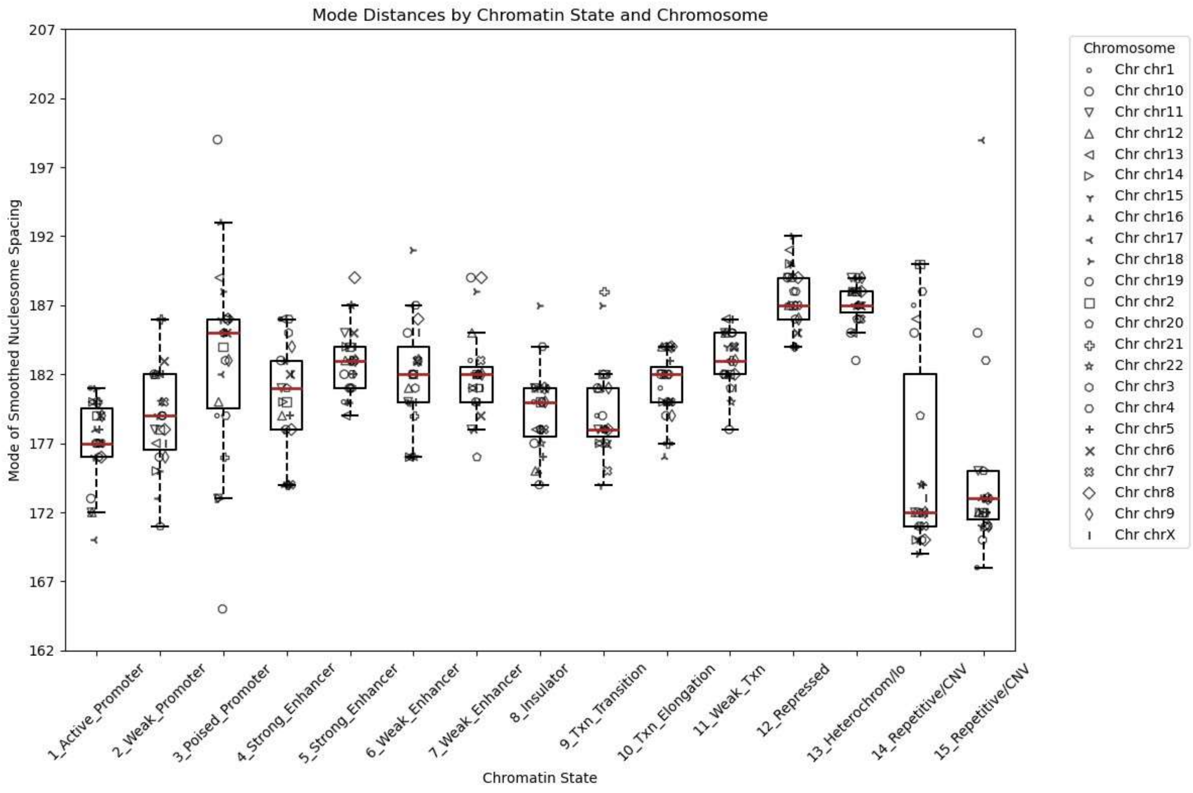
Mode Distances by Chromatin State and Chromosome
Mode of Smoothed Nucleosome Spacing
207
202
197
192
187
182
177
172
167
162
1_Active_Promoter
2_Weak_Promoter
3_Poised_Promoter
4_Strong_Enhancer
5_Strong_Enhancer
6_Weak_Enhancer
7_Weak_Enhancer
8_Insulator
9_Txn_Transition
10_Txn_Elongation
11_Weak_Txn
12_Repressed
13_Heterochrom/lo
14_Repetitive/CNV
15_Repetitive/CNV
Chromatin State
Chromosome
Chr chr1
Chr chr10
Chr chr11
Chr chr12
Chr chr13
Chr chr14
Chr chr15
Chr chr16
Chr chr17
Chr chr18
Chr chr19
Chr chr2
Chr chr20
Chr chr21
Chr chr22
Chr chr3
Chr chr4
Chr chr5
Chr chr6
Chr chr7
Chr chr8
Chr chr9
Chr chrX

**E**

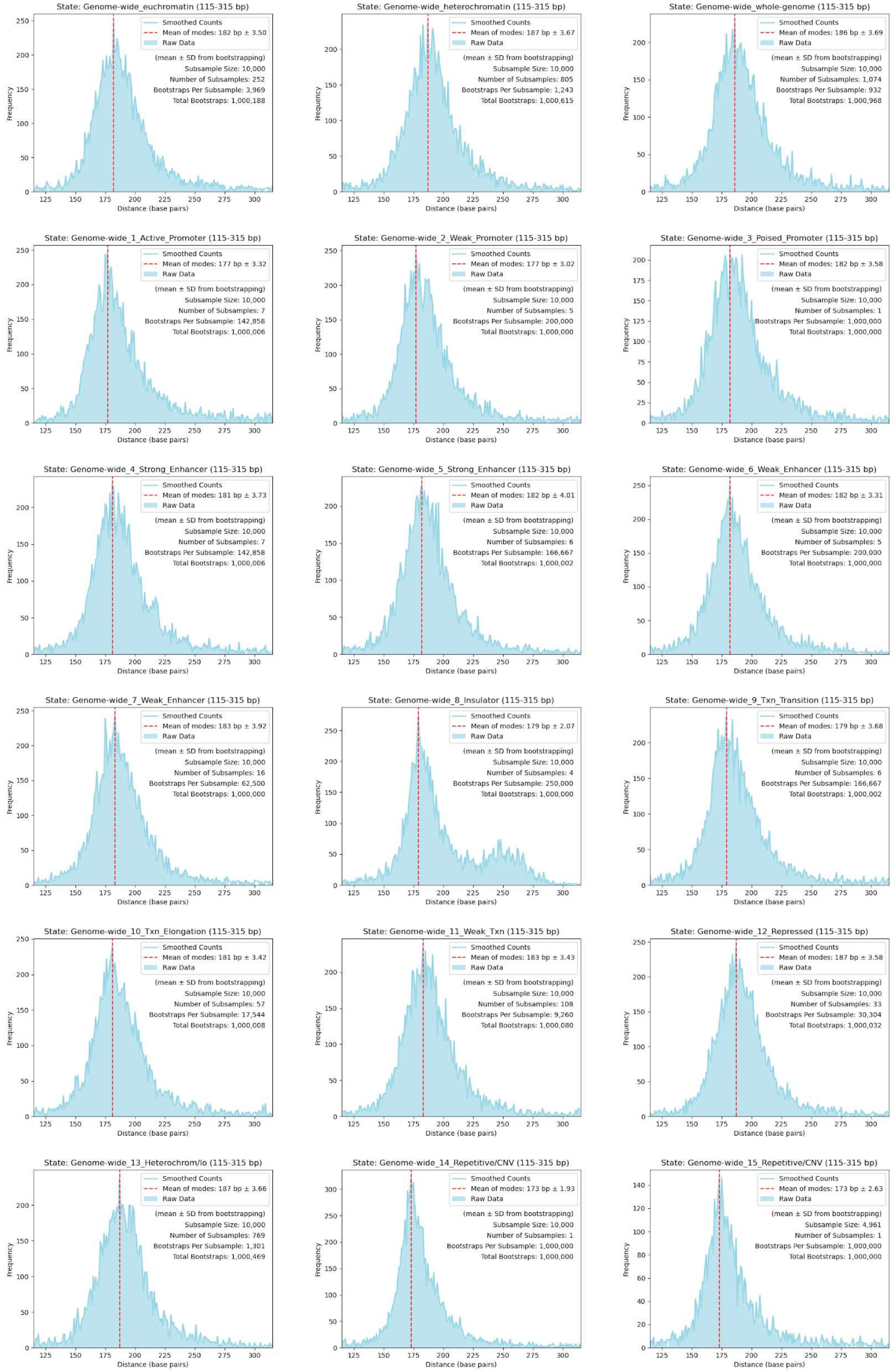
State: Genome-wide_euchromatin (115-315 bp)
Smoothed Counts
Mean of modes: 182 bp ± 3.50
Raw Data
(mean ± SD from bootstrapping)
Subsample Size: 10,000
Number of Subsamples: 252
Bootstraps Per Subsample: 3,969
Total Bootstraps: 1,000,188
State: Genome-wide_heterochromatin (115-315 bp)
Smoothed Counts
Mean of modes: 187 bp ± 3.67
Raw Data
(mean ± SD from bootstrapping)
Subsample Size: 10,000
Number of Subsamples: 805
Bootstraps Per Subsample: 1,243
Total Bootstraps: 1,000,615
State: Genome-wide_whole-genome (115-315 bp)
Smoothed Counts
Mean of modes: 186 bp ± 3.69
Raw Data
(mean ± SD from bootstrapping)
Subsample Size: 10,000
Number of Subsamples: 1,074
Bootstraps Per Subsample: 932
Total Bootstraps: 1,000,968
State: Genome-wide_1_Active_Promoter (115-315 bp)
Smoothed Counts
Mean of modes: 177 bp ± 3.32
Raw Data
(mean ± SD from bootstrapping)
Subsample Size: 10,000
Number of Subsamples: 7
Bootstraps Per Subsample: 142,858
Total Bootstraps: 1,000,006
State: Genome-wide_2_Weak_Promoter (115-315 bp)
Smoothed Counts
Mean of modes: 177 bp ± 3.02
Raw Data
(mean ± SD from bootstrapping)
Subsample Size: 10,000
Number of Subsamples: 5
Bootstraps Per Subsample: 200,000
Total Bootstraps: 1,000,000
State: Genome-wide_3_Poised_Promoter (115-315 bp)
Smoothed Counts
Mean of modes: 182 bp ± 3.58
Raw Data
(mean ± SD from bootstrapping)
Subsample Size: 10,000
Number of Subsamples: 1
Bootstraps Per Subsample: 1,000,000
Total Bootstraps: 1,000,000
State: Genome-wide_4_Strong_Enhancer (115-315 bp)
Smoothed Counts
Mean of modes: 181 bp ± 3.73
Raw Data
(mean ± SD from bootstrapping)
Subsample Size: 10,000
Number of Subsamples: 7
Bootstraps Per Subsample: 142,858
Total Bootstraps: 1,000,006
State: Genome-wide_5_Strong_Enhancer (115-315 bp)
Smoothed Counts
Mean of modes: 182 bp ± 4.01
Raw Data
(mean ± SD from bootstrapping)
Subsample Size: 10,000
Number of Subsamples: 6
Bootstraps Per Subsample: 166,667
Total Bootstraps: 1,000,002
State: Genome-wide_6_Weak_Enhancer (115-315 bp)
Smoothed Counts
Mean of modes: 182 bp ± 3.31
Raw Data
(mean ± SD from bootstrapping)
Subsample Size: 10,000
Number of Subsamples: 5
Bootstraps Per Subsample: 200,000
Total Bootstraps: 1,000,000
State: Genome-wide_7_Weak_Enhancer (115-315 bp)
Smoothed Counts
Mean of modes: 183 bp ± 3.92
Raw Data
(mean ± SD from bootstrapping)
Subsample Size: 10,000
Number of Subsamples: 16
Bootstraps Per Subsample: 62,500
Total Bootstraps: 1,000,000
State: Genome-wide_8_Insulator (115-315 bp)
Smoothed Counts
Mean of modes: 179 bp ± 2.07
Raw Data
(mean ± SD from bootstrapping)
Subsample Size: 10,000
Number of Subsamples: 4
Bootstraps Per Subsample: 250,000
Total Bootstraps: 1,000,000
State: Genome-wide_9_Txn_Transition (115-315 bp)
Smoothed Counts
Mean of modes: 179 bp ± 3.68
Raw Data
(mean ± SD from bootstrapping)
Subsample Size: 10,000
Number of Subsamples: 6
Bootstraps Per Subsample: 166,667
Total Bootstraps: 1,000,002
State: Genome-wide_10_Txn_Elongation (115-315 bp)
Smoothed Counts
Mean of modes: 181 bp ± 3.42
Raw Data
(mean ± SD from bootstrapping)
Subsample Size: 10,000
Number of Subsamples: 57
Bootstraps Per Subsample: 17,544
Total Bootstraps: 1,000,008
State: Genome-wide_11_Weak_Txn (115-315 bp)
Smoothed Counts
Mean of modes: 183 bp ± 3.43
Raw Data
(mean ± SD from bootstrapping)
Subsample Size: 10,000
Number of Subsamples: 108
Bootstraps Per Subsample: 9,260
Total Bootstraps: 1,000,080
State: Genome-wide_12_Repressed (115-315 bp)
Smoothed Counts
Mean of modes: 187 bp ± 3.58
Raw Data
(mean ± SD from bootstrapping)
Subsample Size: 10,000
Number of Subsamples: 33
Bootstraps Per Subsample: 30,304
Total Bootstraps: 1,000,032
State: Genome-wide_13_Heterochrom/lo (115-315 bp)
Smoothed Counts
Mean of modes: 187 bp ± 3.66
Raw Data
(mean ± SD from bootstrapping)
Subsample Size: 10,000
Number of Subsamples: 769
Bootstraps Per Subsample: 1,301
Total Bootstraps: 1,000,469
State: Genome-wide_14_Repetitive/CNV (115-315 bp)
Smoothed Counts
Mean of modes: 173 bp ± 1.93
Raw Data
(mean ± SD from bootstrapping)
Subsample Size: 10,000
Number of Subsamples: 1
Bootstraps Per Subsample: 1,000,000
Total Bootstraps: 1,000,000
State: Genome-wide_15_Repetitive/CNV (115-315 bp)
Smoothed Counts
Mean of modes: 173 bp ± 2.63
Raw Data
(mean ± SD from bootstrapping)
Subsample Size: 4,961
Number of Subsamples: 1
Bootstraps Per Subsample: 1,000,000
Total Bootstraps: 1,000,000
Frequency
Distance (base pairs)

**F**

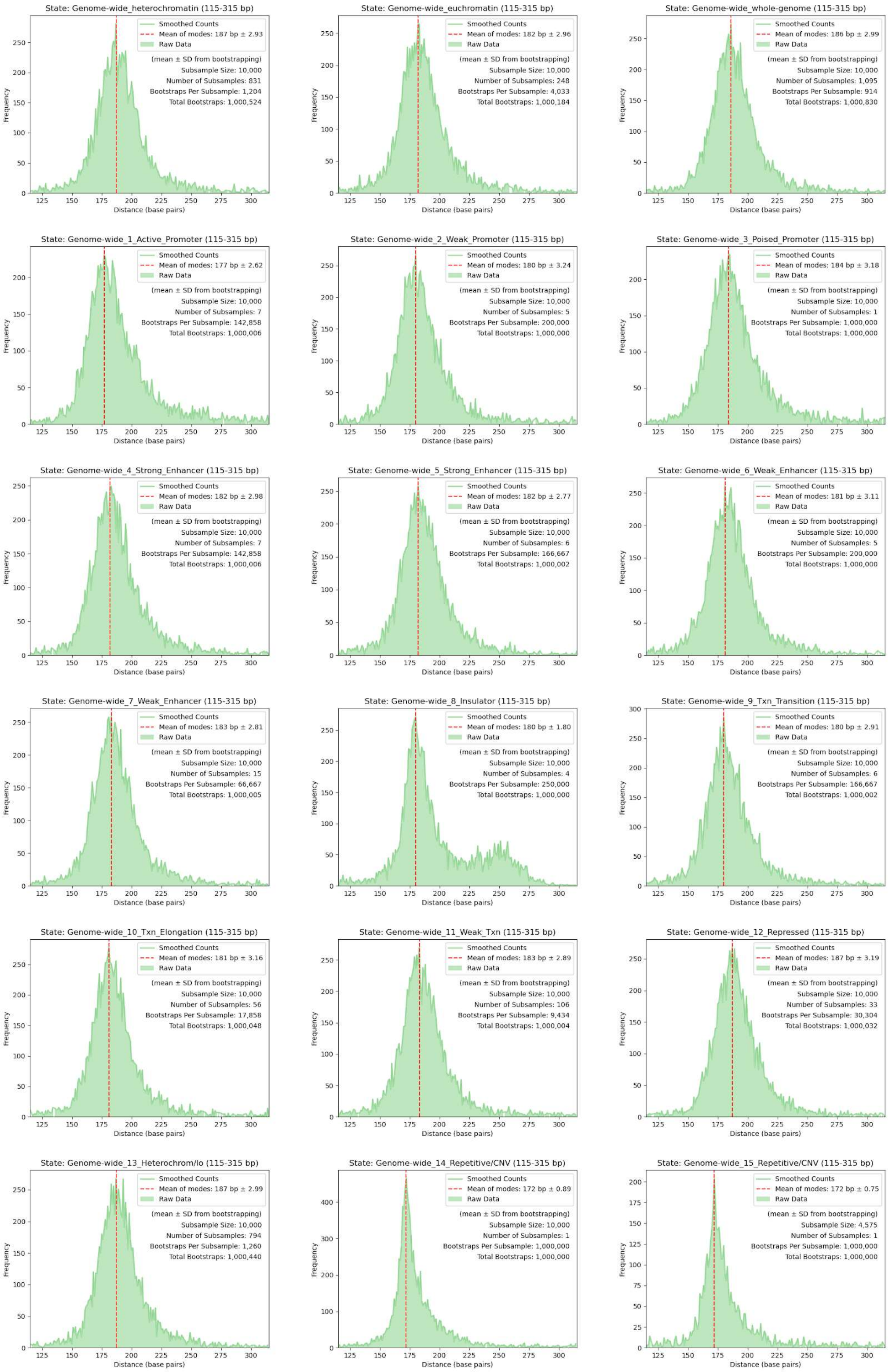
State: Genome-wide_heterochromatin (115-315 bp)
Smoothed Counts
Mean of modes: 187 bp ± 2.93
Raw Data
(mean ± SD from bootstrapping)
Subsample Size: 10,000
Number of Subsamples: 831
Bootstraps Per Subsample: 1,204
Total Bootstraps: 1,000,524
State: Genome-wide_euchromatin (115-315 bp)
Smoothed Counts
Mean of modes: 182 bp ± 2.96
Raw Data
(mean ± SD from bootstrapping)
Subsample Size: 10,000
Number of Subsamples: 248
Bootstraps Per Subsample: 4,033
Total Bootstraps: 1,000,184
State: Genome-wide_whole-genome (115-315 bp)
Smoothed Counts
Mean of modes: 186 bp ± 2.99
Raw Data
(mean ± SD from bootstrapping)
Subsample Size: 10,000
Number of Subsamples: 1,095
Bootstraps Per Subsample: 914
Total Bootstraps: 1,000,830
State: Genome-wide_1_Active_Promoter (115-315 bp)
Smoothed Counts
Mean of modes: 177 bp ± 2.62
Raw Data
(mean ± SD from bootstrapping)
Subsample Size: 10,000
Number of Subsamples: 7
Bootstraps Per Subsample: 142,858
Total Bootstraps: 1,000,006
State: Genome-wide_2_Weak_Promoter (115-315 bp)
Smoothed Counts
Mean of modes: 180 bp ± 3.24
Raw Data
(mean ± SD from bootstrapping)
Subsample Size: 10,000
Number of Subsamples: 5
Bootstraps Per Subsample: 200,000
Total Bootstraps: 1,000,000
State: Genome-wide_3_Poised_Promoter (115-315 bp)
Smoothed Counts
Mean of modes: 184 bp ± 3.18
Raw Data
(mean ± SD from bootstrapping)
Subsample Size: 10,000
Number of Subsamples: 1
Bootstraps Per Subsample: 1,000,000
Total Bootstraps: 1,000,000
State: Genome-wide_4_Strong_Enhancer (115-315 bp)
Smoothed Counts
Mean of modes: 182 bp ± 2.98
Raw Data
(mean ± SD from bootstrapping)
Subsample Size: 10,000
Number of Subsamples: 7
Bootstraps Per Subsample: 142,858
Total Bootstraps: 1,000,006
State: Genome-wide_5_Strong_Enhancer (115-315 bp)
Smoothed Counts
Mean of modes: 182 bp ± 2.77
Raw Data
(mean ± SD from bootstrapping)
Subsample Size: 10,000
Number of Subsamples: 6
Bootstraps Per Subsample: 166,667
Total Bootstraps: 1,000,002
State: Genome-wide_6_Weak_Enhancer (115-315 bp)
Smoothed Counts
Mean of modes: 181 bp ± 3.11
Raw Data
(mean ± SD from bootstrapping)
Subsample Size: 10,000
Number of Subsamples: 5
Bootstraps Per Subsample: 200,000
Total Bootstraps: 1,000,000
State: Genome-wide_7_Weak_Enhancer (115-315 bp)
Smoothed Counts
Mean of modes: 183 bp ± 2.81
Raw Data
(mean ± SD from bootstrapping)
Subsample Size: 10,000
Number of Subsamples: 15
Bootstraps Per Subsample: 66,667
Total Bootstraps: 1,000,005
State: Genome-wide_8_Insulator (115-315 bp)
Smoothed Counts
Mean of modes: 180 bp ± 1.80
Raw Data
(mean ± SD from bootstrapping)
Subsample Size: 10,000
Number of Subsamples: 4
Bootstraps Per Subsample: 250,000
Total Bootstraps: 1,000,000
State: Genome-wide_9_Txn_Transition (115-315 bp)
Smoothed Counts
Mean of modes: 180 bp ± 2.91
Raw Data
(mean ± SD from bootstrapping)
Subsample Size: 10,000
Number of Subsamples: 6
Bootstraps Per Subsample: 166,667
Total Bootstraps: 1,000,002
State: Genome-wide_10_Txn_Elongation (115-315 bp)
Smoothed Counts
Mean of modes: 181 bp ± 3.16
Raw Data
(mean ± SD from bootstrapping)
Subsample Size: 10,000
Number of Subsamples: 56
Bootstraps Per Subsample: 17,858
Total Bootstraps: 1,000,048
State: Genome-wide_11_Weak_Txn (115-315 bp)
Smoothed Counts
Mean of modes: 183 bp ± 2.89
Raw Data
(mean ± SD from bootstrapping)
Subsample Size: 10,000
Number of Subsamples: 106
Bootstraps Per Subsample: 9,434
Total Bootstraps: 1,000,004
State: Genome-wide_12_Repressed (115-315 bp)
Smoothed Counts
Mean of modes: 187 bp ± 3.19
Raw Data
(mean ± SD from bootstrapping)
Subsample Size: 10,000
Number of Subsamples: 33
Bootstraps Per Subsample: 30,304
Total Bootstraps: 1,000,032
State: Genome-wide_13_Heterochrom/lo (115-315 bp)
Smoothed Counts
Mean of modes: 187 bp ± 2.99
Raw Data
(mean ± SD from bootstrapping)
Subsample Size: 10,000
Number of Subsamples: 794
Bootstraps Per Subsample: 1,260
Total Bootstraps: 1,000,440
State: Genome-wide_14_Repetitive/CNV (115-315 bp)
Smoothed Counts
Mean of modes: 172 bp ± 0.89
Raw Data
(mean ± SD from bootstrapping)
Subsample Size: 10,000
Number of Subsamples: 1
Bootstraps Per Subsample: 1,000,000
Total Bootstraps: 1,000,000
State: Genome-wide_15_Repetitive/CNV (115-315 bp)
Smoothed Counts
Mean of modes: 172 bp ± 0.75
Raw Data
(mean ± SD from bootstrapping)
Subsample Size: 4,575
Number of Subsamples: 1
Bootstraps Per Subsample: 1,000,000
Total Bootstraps: 1,000,000
Frequency
Distance (base pairs)

**Supplementary Figure S3. Chromatin state mode interpeak distances of nucleosome protection peaks.** Nucleosome protection peaks (NPPs) are those of the CH01 (**A**,**C**,**E**) and CA01 (**B**,**D**,**F**) sample pools. (**A**,**B**) Plots of all NPP interpeak distances between 115-315 bp. (**C**,**D**) Box plots of mode interpeak distances for each chromosome. The bottom line of each box plot represents the 25th percentile, top line the 75th percentile, and red middle line the median. Whiskers extend up to a maximum of 1.5 times the height of the box. Any values that fall outside this range are classified as outliers. (**A**-**D**) Mode interpeak distances are calculated after Savitzky-Golay smoothing (window size = 25, polynomial of order = 3). (**E**,**F**) Subsampled and bootstrapped NPP interpeak distances. To account for differences in sample sizes between chromatin states, NPP interpeak distances for each state were randomly shuffled and split into subsamples of 10,000 (slightly less than the number of interpeak distances in states 3 and 14, the second and third smallest sample sizes). 1 million bootstrap iterations were then allocated for each chromatin state, with each subsample undergoing an equal share of the total bootstrap iterations. Plots are those of a single bootstrap from a single subsample whose mode matched the most frequent mode across all bootstraps (mean of modes). Protection peak data from Synder et al. (2016)[4] were categorized into Ernst et al. (2011)[5] ChromHMM chromatin states of the GM12878 B-lymphoblastoid cell line.

A

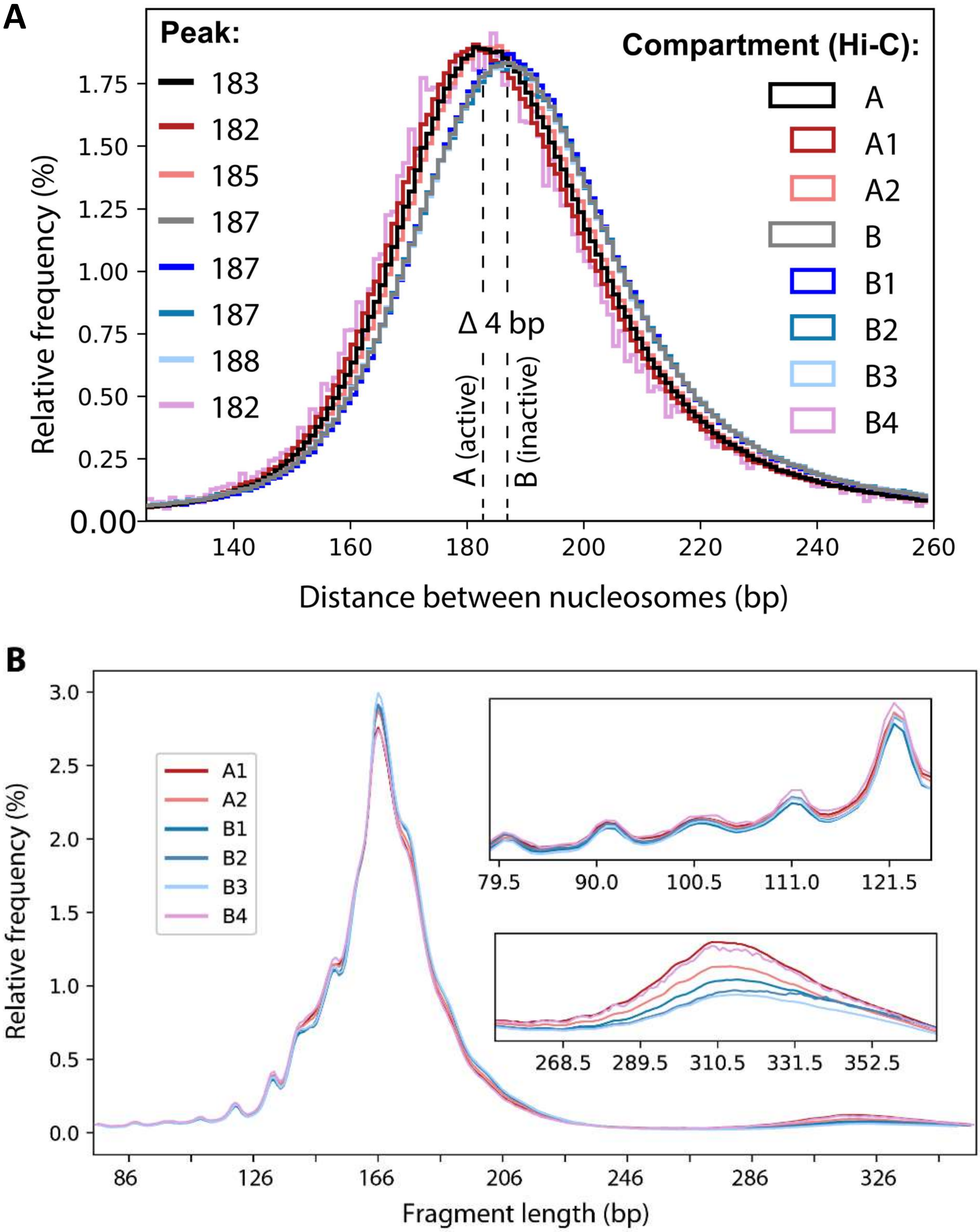

A
Peak:
183
182
185
187
187
187
188
182
Compartment (Hi-C):
A
A1
A2
B
B1
B2
B3
B4
Δ 4 bp
A (active)
B (inactive)
Relative frequency (%)
Distance between nucleosomes (bp)
B
A1
A2
B1
B2
B3
B4
Relative frequency (%)
Fragment length (bp)

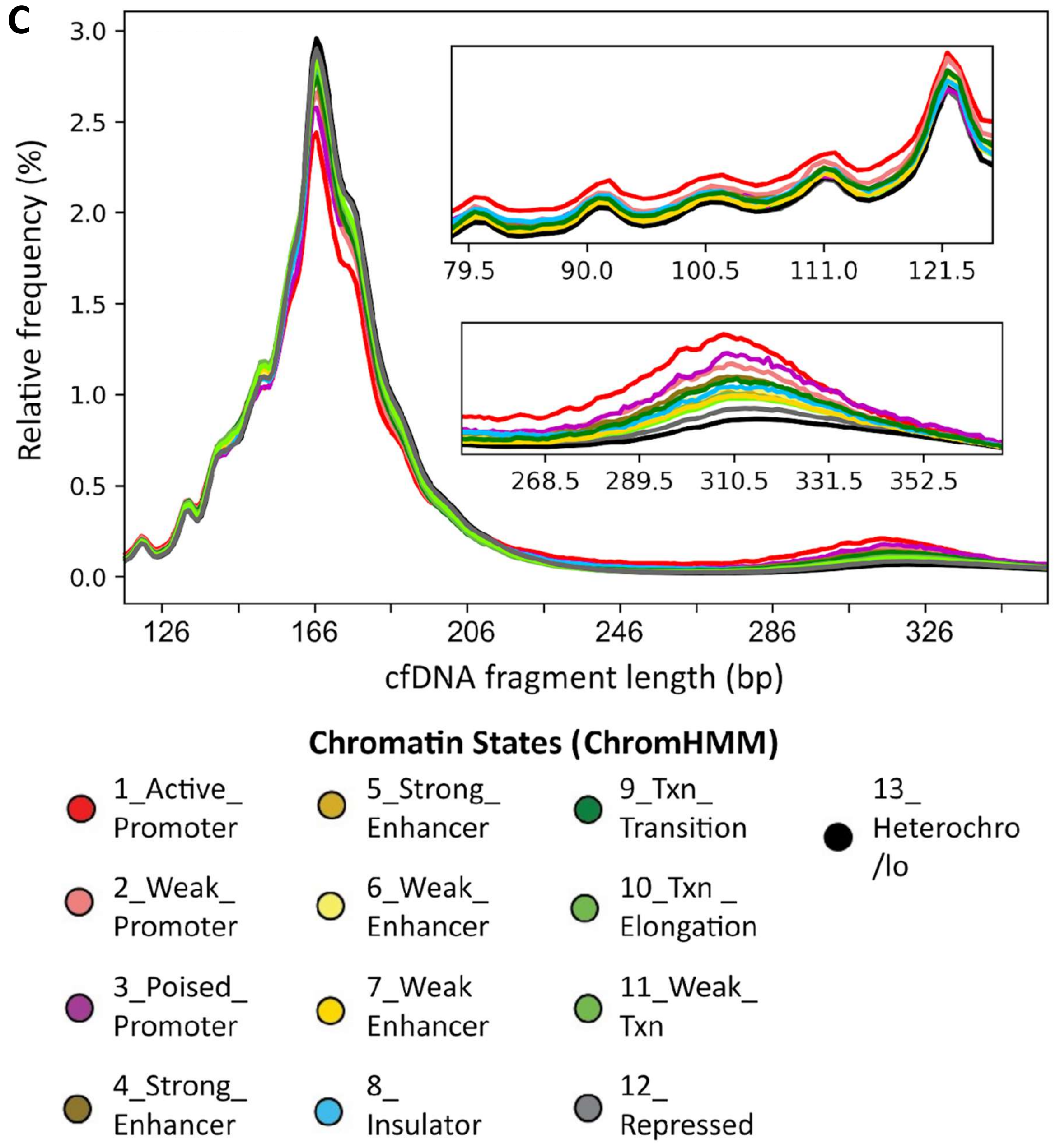


**Supplementary Figure S4. Size distributions and nucleosome spacing of cfDNA fragments aligned to chromatin subcompartments.** (**A**) Distance between adjacent nucleosomes is different for Rao et al. (2014)[6] chromatin compartments and subcompartments derived from Hi-C contact maps of the GM12878 B-lymphoblastoid cell line. (**B**) Savitzky-Golay smoothed (window size = 25, polynomial of order = 3) histograms of cfDNA fragments sizes aligned to chromatin compartments and subcompartments. (**C**) Savitzky-Golay smoothed (window size = 25, polynomial of order = 3) histograms of cfDNA fragments sizes aligned to ChromHMM chromatin states of Ernst et al. (2011)[5] from the GM12878 B-lymphoblastoid cell line.

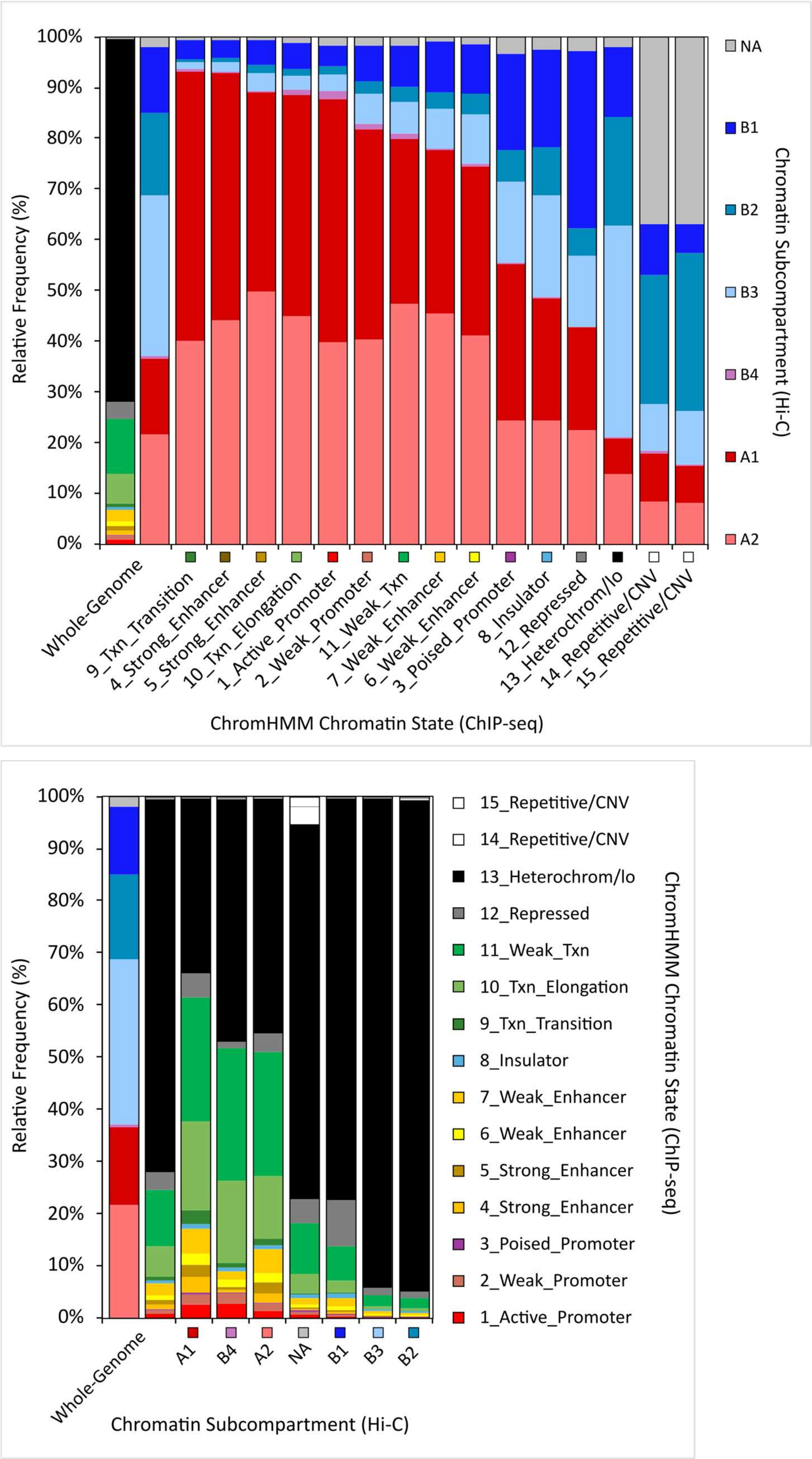


**Supplementary Figure S5. Stacked bar charts showing the correlations between chromatin states and chromatin sub-compartments.** Chromatin states are those of Ernst et al. (2011)[5] from the GM12878 B-lymphoblastoid cell line. Chromatin A/B sub-compartments are those of Rao et al. (2014)[6] from the GM12878 B-lymphoblastoid cell line.

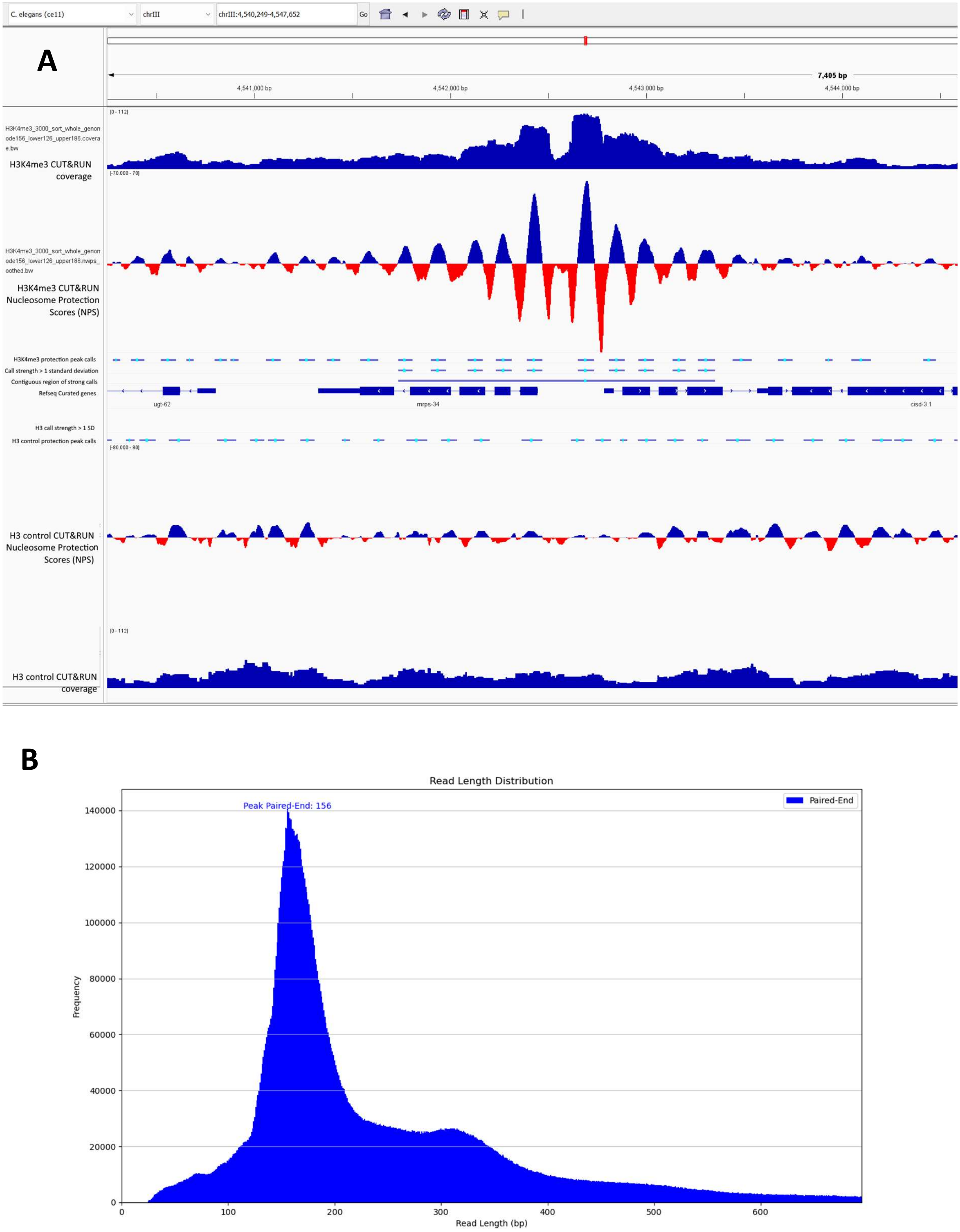


**Supplementary Figure S6. Nucleosome protecting scoring and peak calling on CUT&RUN data from *C. elegans*.** (**A**) Nucleosome protection scoring applied to CUT&RUN sequencing data. Strong nucleosome calls can be seen at sites of H3K4me3, which is associated with transcription start sites. A large gap between nucleosome calls is present, suggesting the presence of bound transcription factors at this site. (**B**) A central fragment length of 156 bp was used for scoring to match the mode fragment size of CUT&RUN, as opposed to the 166 bp used for cfDNA.

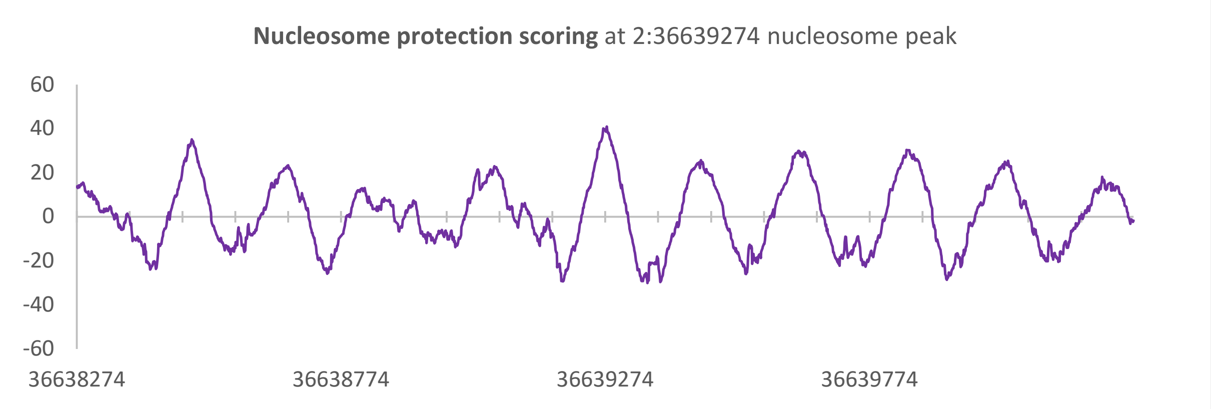
Nucleosome protection scoring at 2:36639274 nucleosome peak
60
40
20
0
-20
-40
-60
36638274
36638774
36639274
36639774

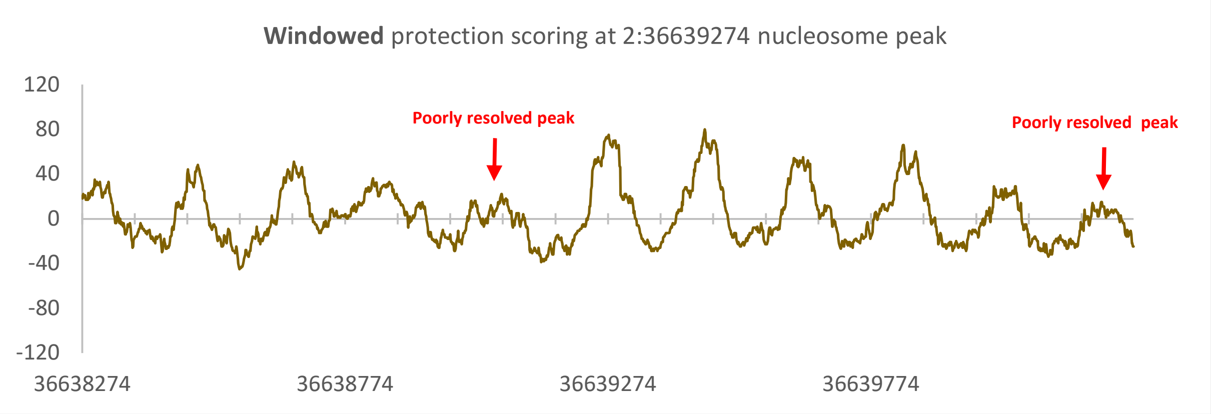
Windowed protection scoring at 2:36639274 nucleosome peak
Poorly resolved peak
Poorly resolved peak
120
80
40
0
-40
-80
-120
36638274
36638774
36639274
36639774

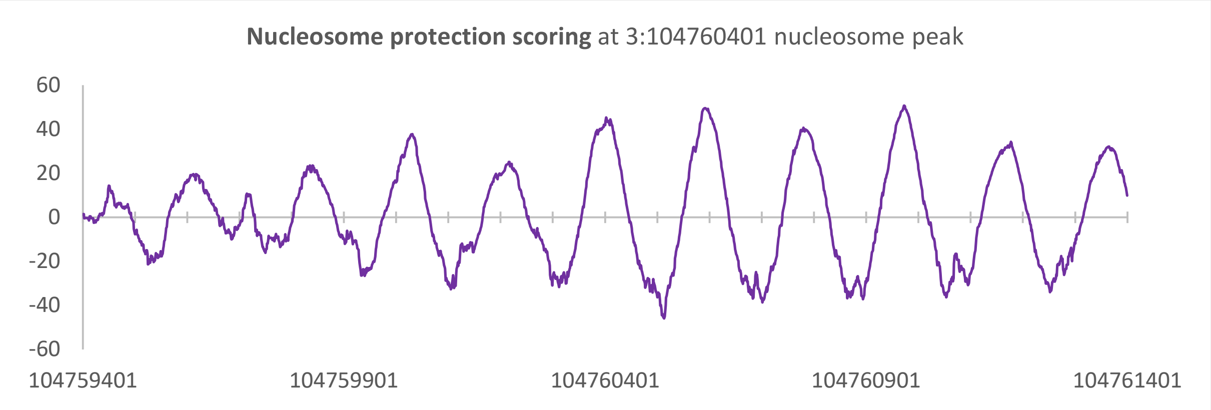
Nucleosome protection scoring at 3:104760401 nucleosome peak
60
40
20
0
-20
-40
-60
104759401
104759901
104760401
104760901
104761401

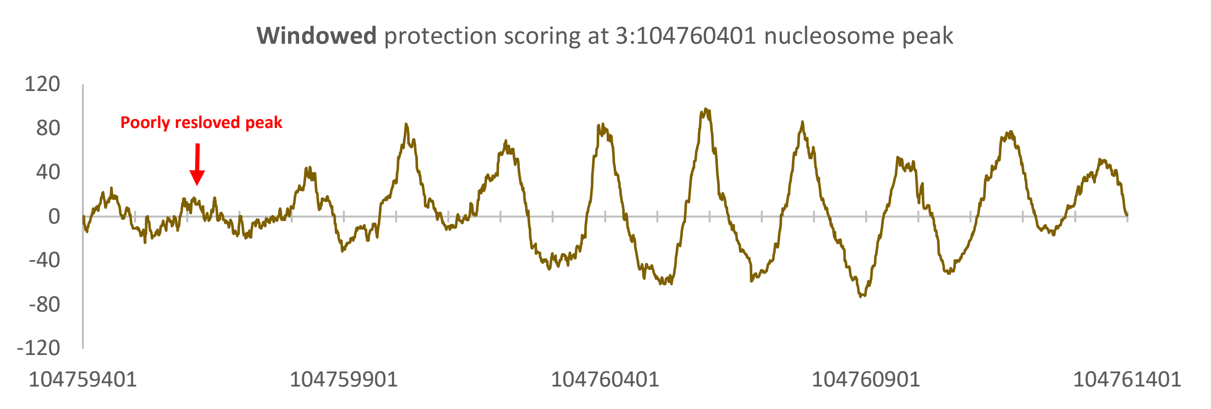
Windowed protection scoring at 3:104760401 nucleosome peak
Poorly resloved peak
120
80
40
0
-40
-80
-120
104759401
104759901
104760401
104760901
104761401

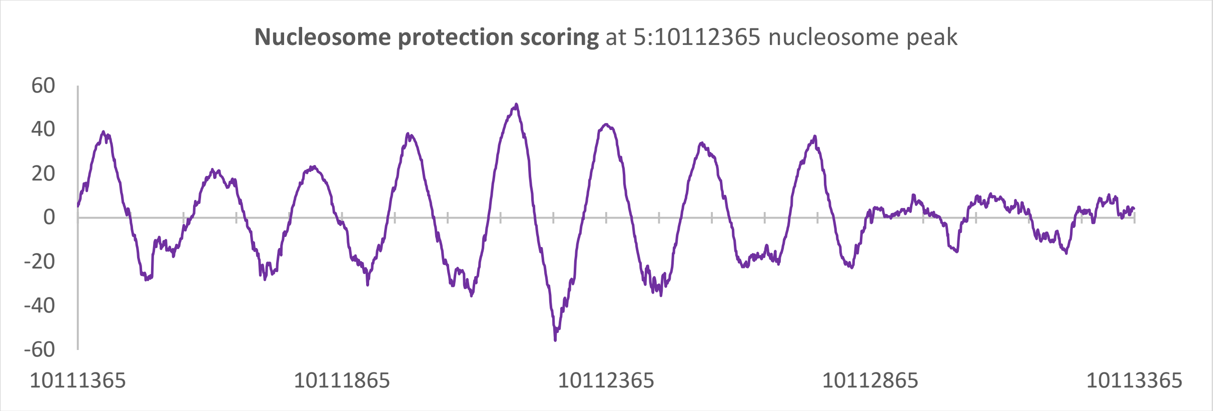
Nucleosome protection scoring at 5:10112365 nucleosome peak
60
40
20
0
-20
-40
-60
10111365
10111865
10112365
10112865
10113365

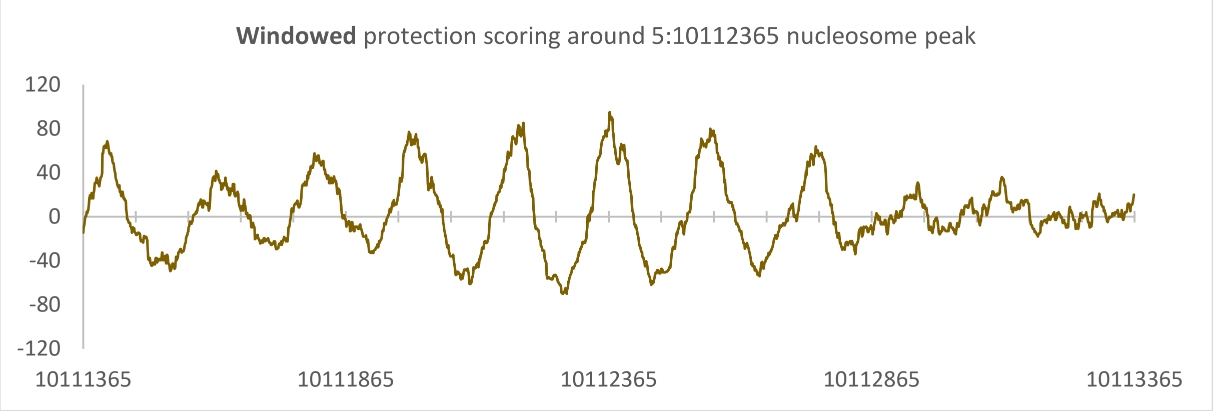
Windowed protection scoring around 5:10112365 nucleosome peak
120
80
40
0
-40
-80
-120
10111365
10111865
10112365
10112865
10113365

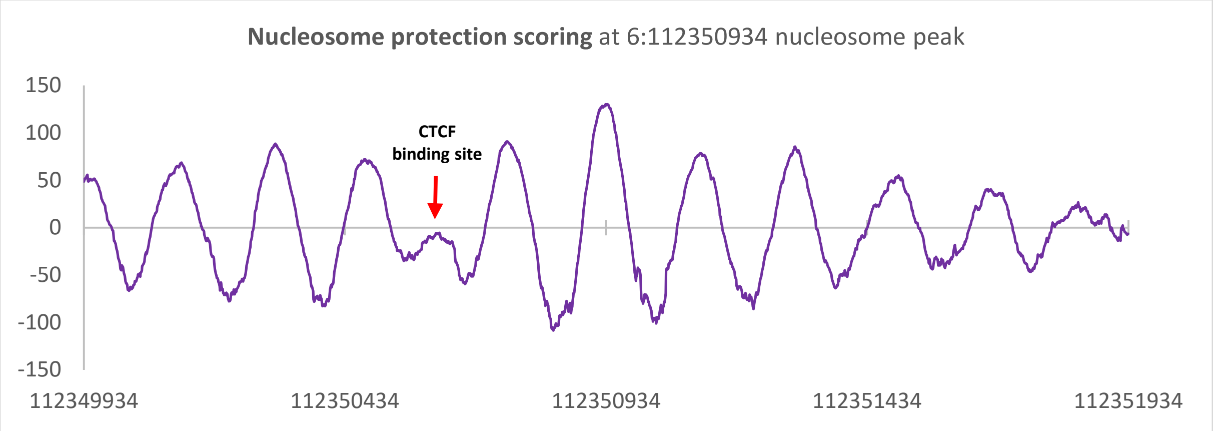
Nucleosome protection scoring at 6:112350934 nucleosome peak
150
100
50
0
-50
-100
-150
CTCF binding site
112349934
112350434
112350934
112351434
112351934

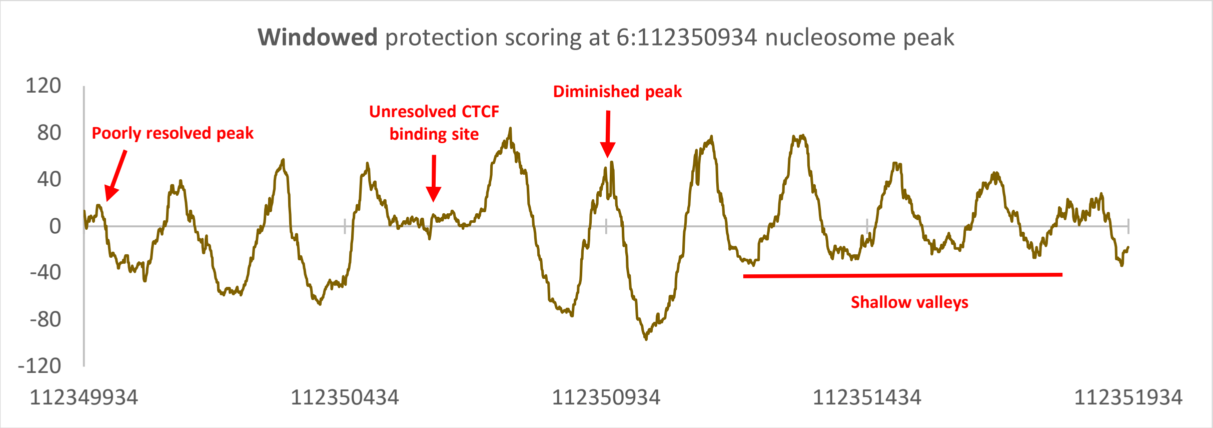
Windowed protection scoring at 6:112350934 nucleosome peak
120
80
40
0
-40
-80
-120
Poorly resolved peak
Unresolved CTCF binding site
Diminished peak
Shallow valleys
112349934
112350434
112350934
112351434
112351934

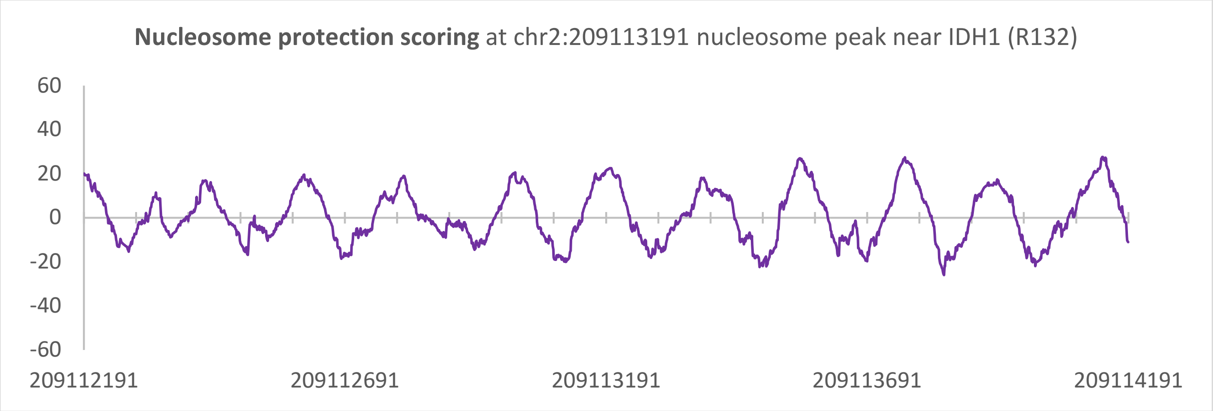
**Nucleosome protection scoring** at chr2:209113191 nucleosome peak near IDH1 (R132)
60
40
20
0
-20
-40
-60
209112191
209112691
209113191
209113691
209114191

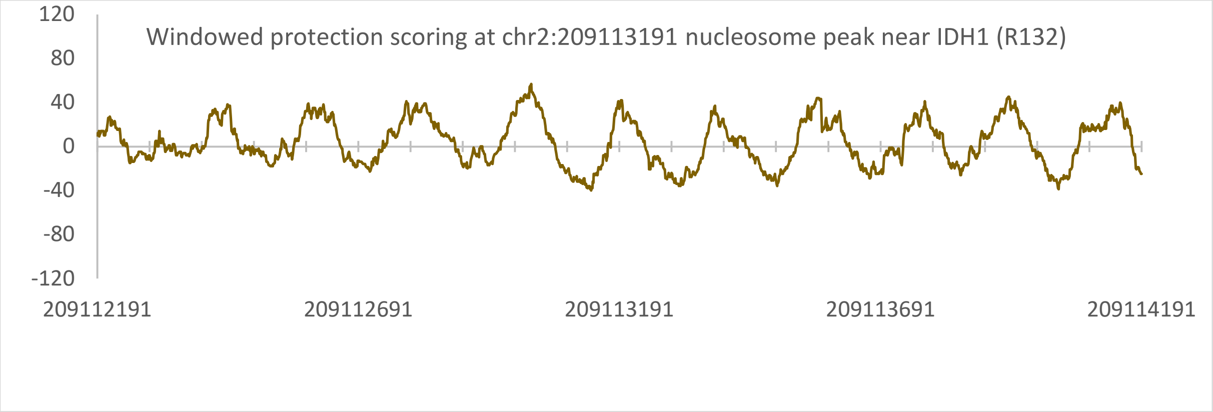
Windowed protection scoring at chr2:209113191 nucleosome peak near IDH1 (R132)
120
80
40
0
-40
-80
-120
209112191
209112691
209113191
209113691
209114191

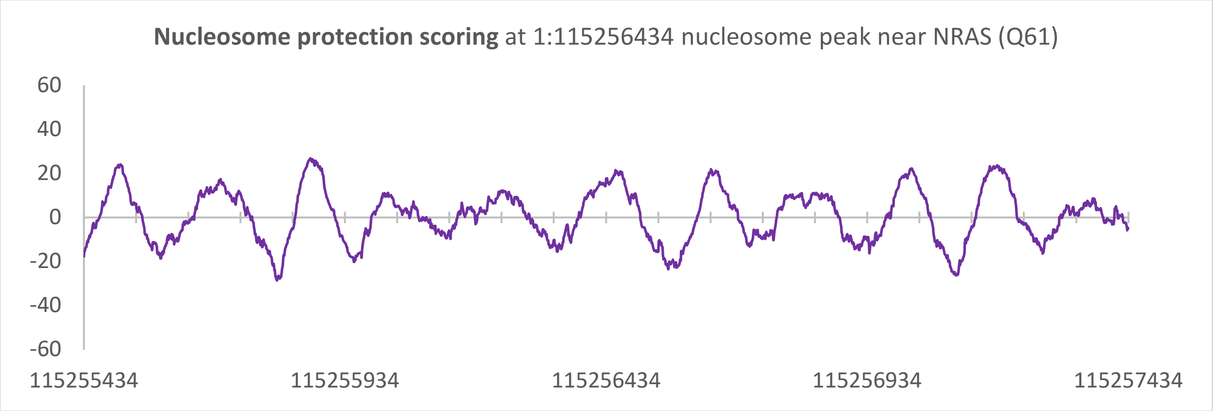
**Nucleosome protection scoring** at 1:115256434 nucleosome peak near NRAS (Q61)
60
40
20
0
-20
-40
-60
115255434
115255934
115256434
115256934
115257434

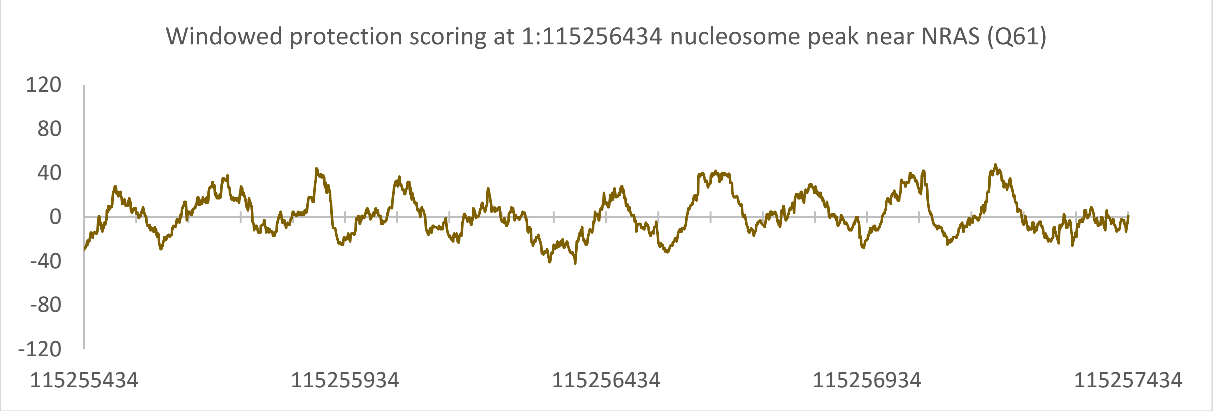
Windowed protection scoring at 1:115256434 nucleosome peak near NRAS (Q61)
120
80
40
0
-40
-80
-120
115255434
115255934
115256434
115256934
115257434

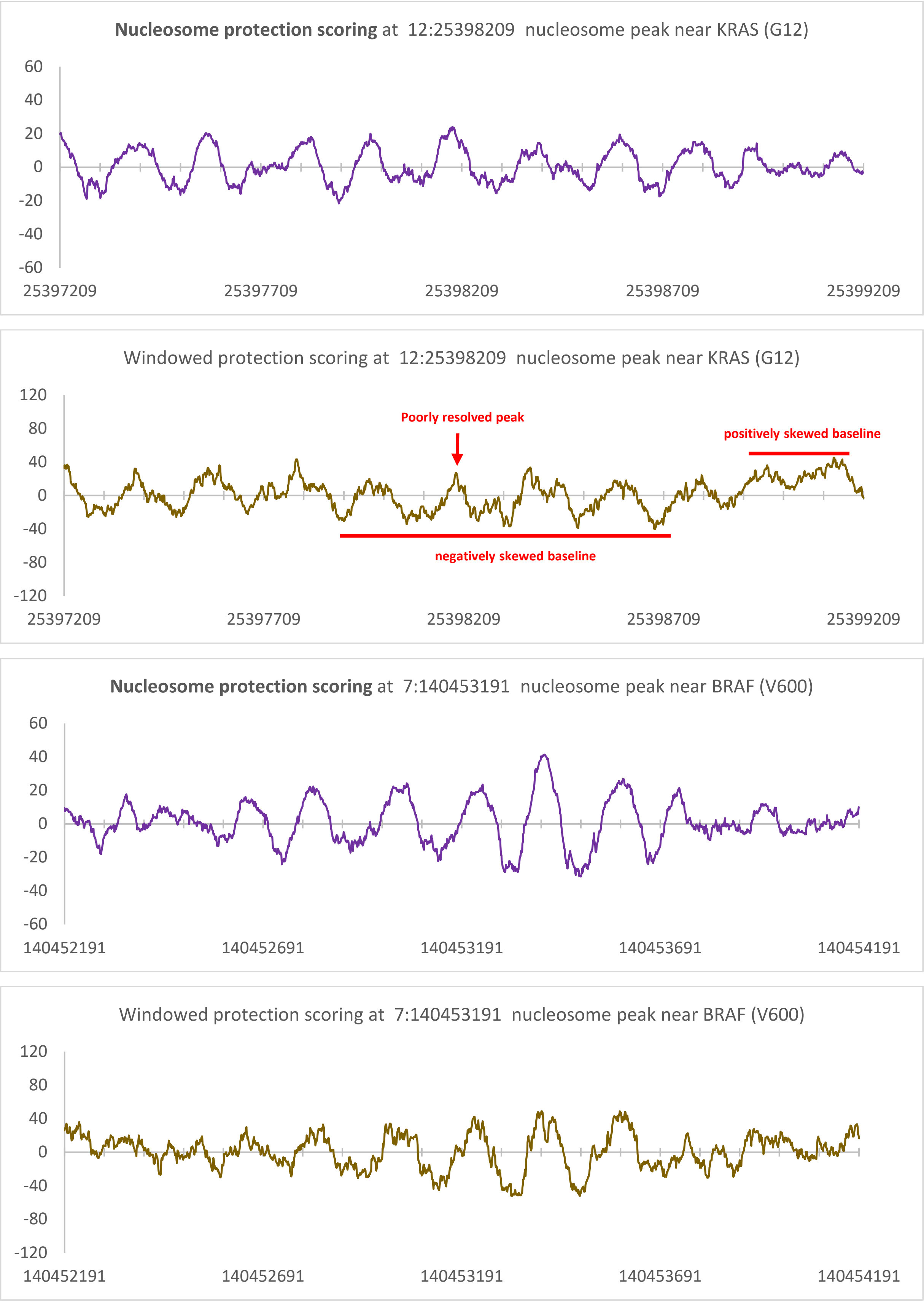


**Supplementary Figure S7. Comparison of new model nucleosome prediction scoring method with windowed protection scoring.** New method of nucleosome protection scoring, and previous method of windowed protection scoring were conducted on cfDNA sequences from all the female samples in the CA01 sample pool. Data presented has been left unsmoothed for better comparison between methods. However, scores are Savitzky-Golay smoothed (window size = 21, polynomial of order = 3) in both methods for peak calling.

**A**

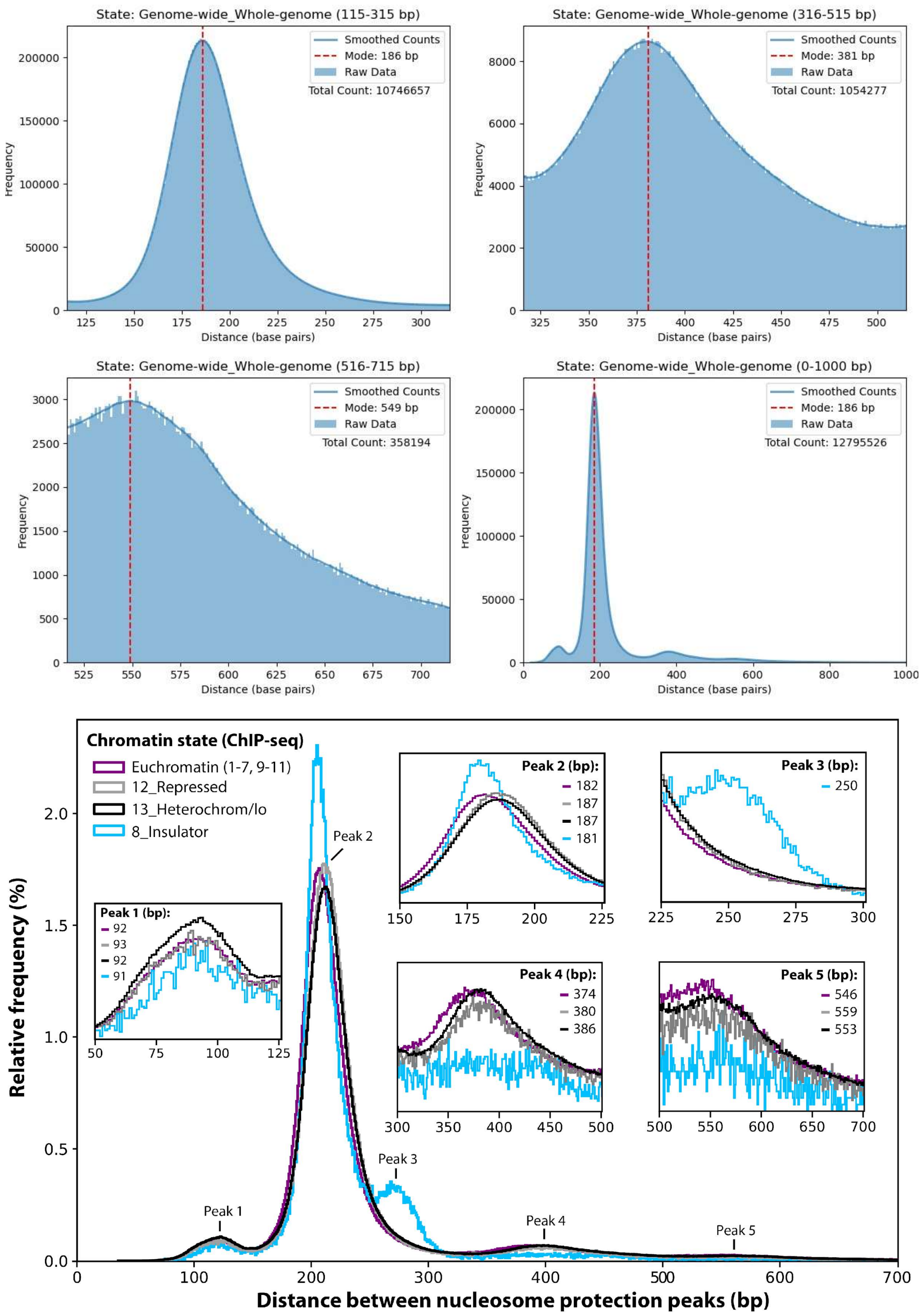
State: Genome-wide_Whole-genome (115-315 bp)
Smoothed Counts
Mode: 186 bp
Raw Data
Total Count: 10746657
Frequency
Distance (base pairs)
State: Genome-wide_Whole-genome (316-515 bp)
Smoothed Counts
Mode: 381 bp
Raw Data
Total Count: 1054277
State: Genome-wide_Whole-genome (516-715 bp)
Smoothed Counts
Mode: 549 bp
Raw Data
Total Count: 358194
State: Genome-wide_Whole-genome (0-1000 bp)
Smoothed Counts
Mode: 186 bp
Raw Data
Total Count: 12795526
Chromatin state (ChIP-seq)
Euchromatin (1-7, 9-11)
12_Repressed
13_Heterochrom/lo
8_Insulator
Peak 1 (bp): 92, 93, 92, 91
Peak 2 (bp): 182, 187, 187, 181
Peak 3 (bp): 250
Peak 4 (bp): 374, 380, 386
Peak 5 (bp): 546, 559, 553
Peak 1
Peak 2
Peak 3
Peak 4
Peak 5
Relative frequency (%)
Distance between nucleosome protection peaks (bp)

B

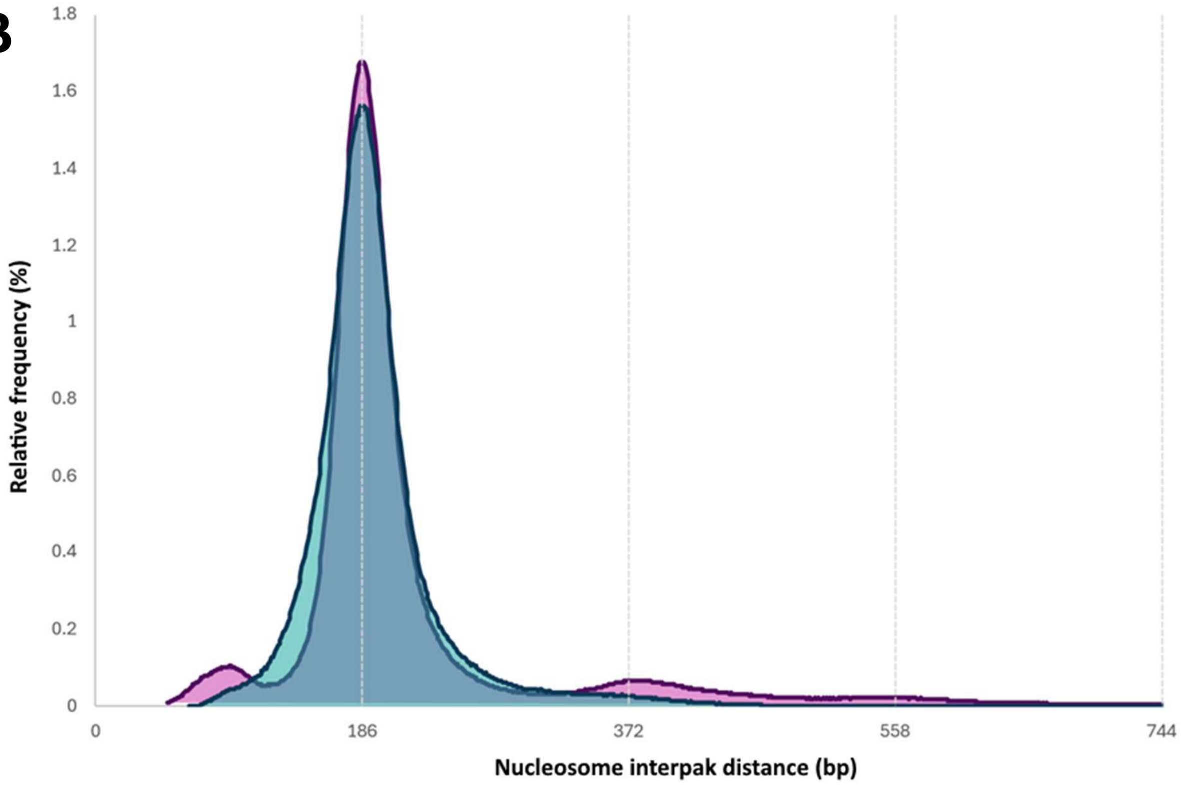


**Supplementary Figure S8. Nucleosome spacing histograms of healthy pooled CH01 cfDNA sample**.

(**A**) Protection peak data from Synder et al. (2016)[4] were categorized into Ernst et al. (2011)[5] ChromHMM chromatin states of the GM12878 B-lymphoblastoid cell line. Mode interpeak distances after Savitzky-Golay smoothing (window size = 25, polynomial of order = 3) are displayed for all major and minor peaks. The CH01 nucleosome map nears saturation, with nearly 13 million nucleosome peak calls in the 2.86 billion bp of the mappable GRCh37 genome, equating to an average of one nucleosome every 221 bp and 84% saturation (with saturation based on the mode interpeak distance of 186 bp). Furthermore, the NPP interpeak distance histogram shows one major peak with a mode of 186 bp, and two minor peaks at 381 bp and 550 bp, which are increments of the 186 bp peak. If we assume distances corresponding each incremental peak contain additional unmapped nucleosomes (1 for 316-515 bp and 2 for 516-715 bp), the number of nucleosomes increases by 1.77 million, which averages to one nucleosome every 195 bp and 95% saturation.

(**B**) Comparison of genome-wide nucleosome spacing distributions between scoring methods. Density plots of the interpeak distances between nucleosome peak calls using windowed protection scoring (pink) and nucleosome protection scoring (blue) methods.

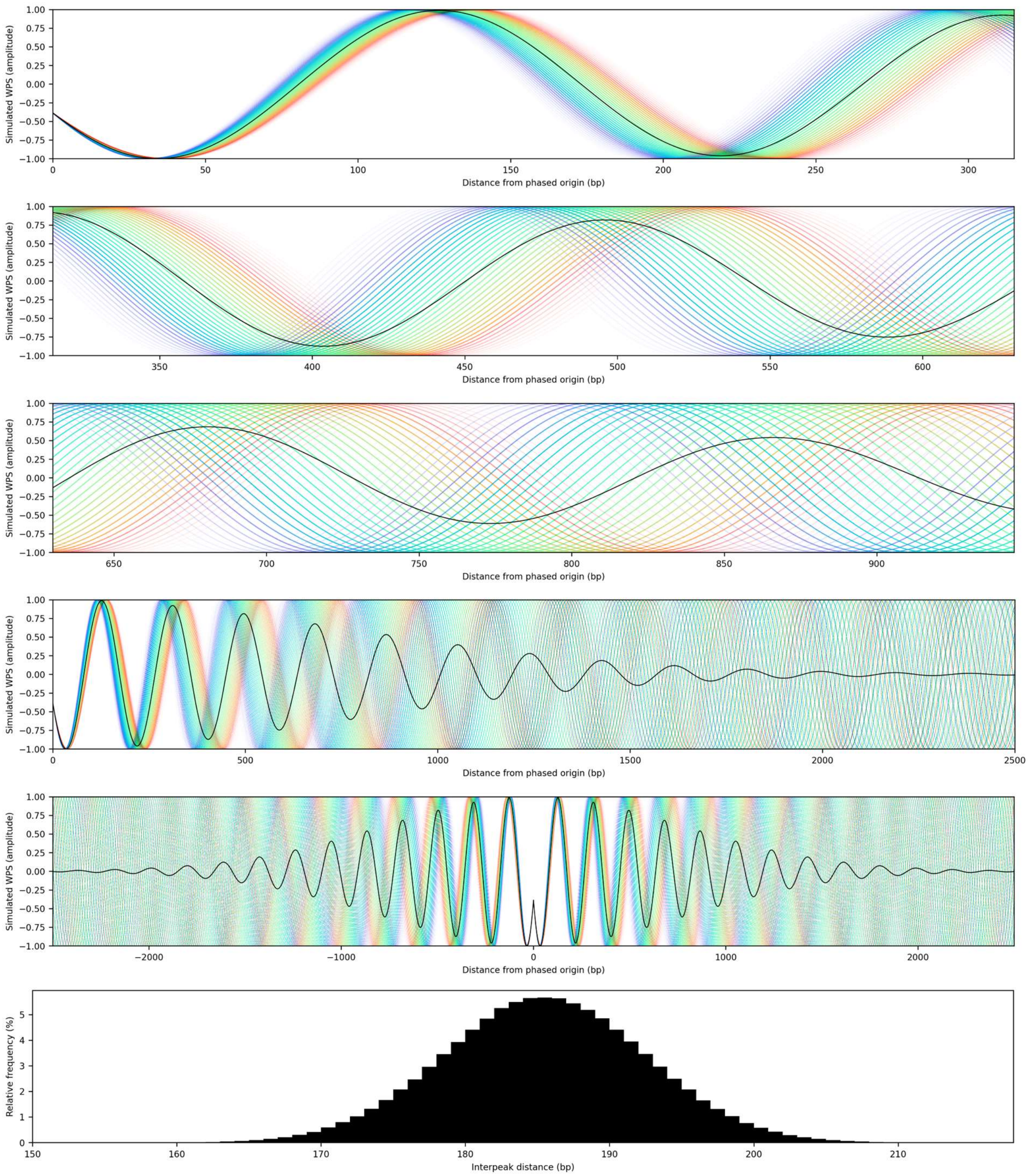


**Supplementary Figure S9. Sine wave phasing simulation of CTCF binding sites using Gaussian distribution of interpeak distances.** Rainbows are composed of sine waves at different frequencies. Each coloured line is a sine wave with an interpeak distance between 151 bp [violet] and 218 bp [red]). The black line is an averaged summation of all the sine waves depicted in rainbow, with each interpeak distance included multiple times based on the relative frequencies in the depicted histogram. This histogram is a Gaussian distribution generated to produce a similar signal fading and shape as aggregate WPS plots aligned to occupied CTCF binding sites. Multiple plots at different scales are included for greater clarity of wave dispersion and fading of aggerate signal.

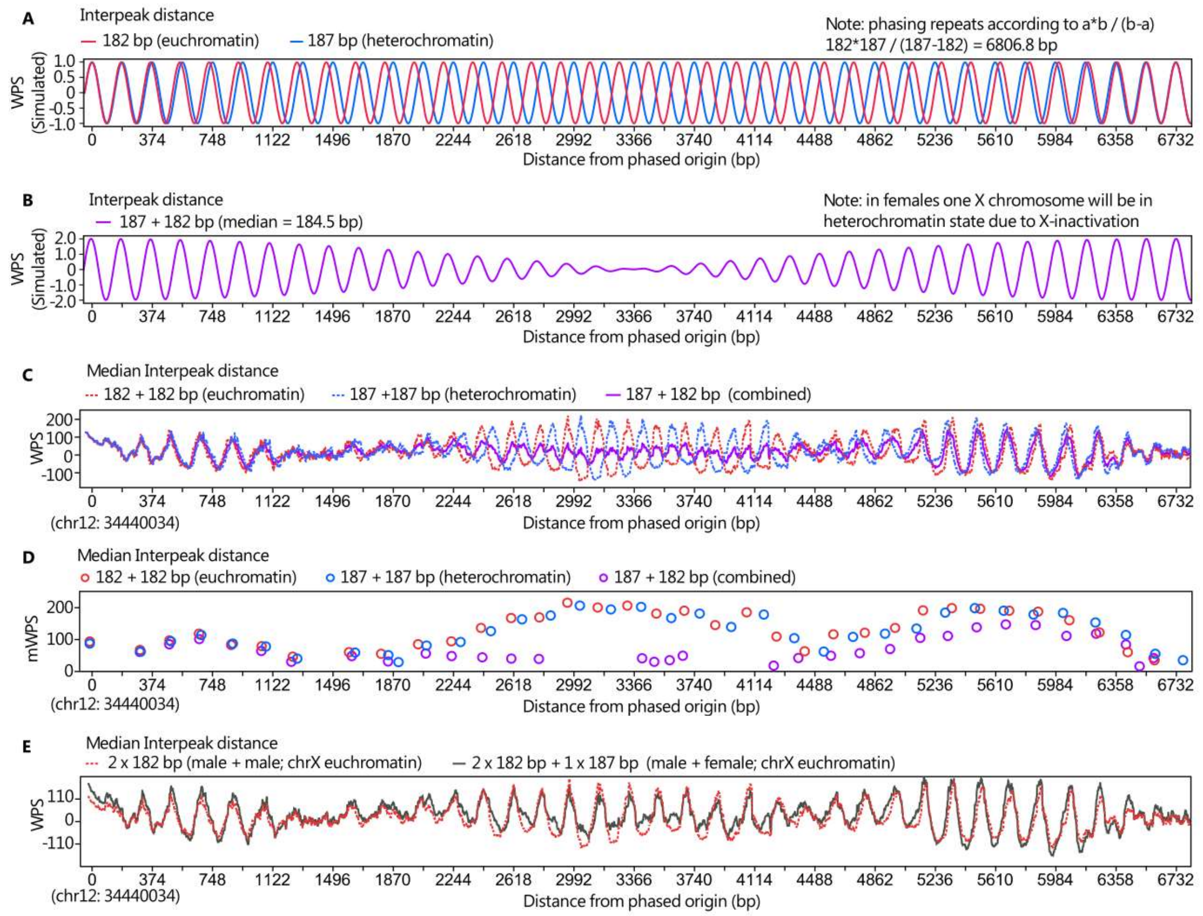


**F**

| Sample | Median interpeak distance (115-315bp): Heterochromatin chrX | chr7 | 1-22 | Euchromatin chrX | chr7 | 1-22 | Health Status | Mean Coverage | Sex | Δ median spacing (Hetero - Eu) chrX | chr7 | 1-22 |
|---|---|---|---|---|---|---|---|---|---|---|---|---|
| BRA01 | 190 | 190 | 190 | 188 | 186 | 186 | Cancer | 17.28 | Mixed† | 2 | 4 | 4 |
| COL01 | 189 | 188 | 188 | 187 | 185 | 185 | Cancer | 19.94 | Mixed† | 2 | 3 | 3 |
| BRE01 | 188 | 188 | 188 | 187 | 185 | 185 | Cancer | 23.12 | Female† | 1* | 3 | 3 |
| IC17 | 186 | 186 | 186 | 183 | 182 | 182 | Cancer | 28.25 | Male | 3 | 4 | 4 |
| IC37 | 186 | 186 | 186 | 185 | 182.5 | 183 | Cancer | 30.59 | Female | 1* | 3.5 | 3 |
| IH02 | 188 | 188 | 188 | 185 | 184 | 184 | Healthy | 30.08 | Male | 3 | 4 | 4 |
| BBC01 | 189 | 188 | 189 | 186 | 184 | 185 | Cancer | 62.17 | Mixed† | 3 | 4 | 4 |
| BH01 | 190 | 190 | 190 | 186 | 186 | 185 | Healthy | 96.86 | Mixed† | 4 | 4 | 5 |
| IH01 | 189 | 189 | 189 | 186 | 186 | 185 | Healthy | 103.89 | Male | 3 | 3 | 4 |
| CA01♀ | 188 | 188 | 188 | 186 | 183 | 183 | Cancer | 121.19 | Female† | 2* | 5 | 5 |
| CA01♂ | 187 | 187 | 187 | 183 | 183 | 183 | Cancer | 150.80 | Male† | 4 | 4 | 4 |
| CH01 | 189 | 190 | 190 | 185 | 185 | 185 | Healthy | 231.32 | Mixed† | 4 | 5 | 5 |
| CA01 | 189 | 189 | 189 | 185 | 184 | 184 | Mixed | 697.76 | Mixed† | 4 | 5 | 5 |

**Supplementary Figure S10. Effects of X-inactivation on NRL of female chrX euchromatin.** (**A**) Sine wave phasing simulation at frequencies of 182 bp (euchromatin) and 187 bp (heterochromatin) NRLs. (**B**) Beat frequency resulting from the addition of waves with 182 and 187 bp frequencies from plot A, as would occur in euchromatin regions of female X chromosomes. (**C**) WPS plots of BBC01 sample pool at a well-studied alpha-satellite array. Fragment positions were artificially shifted to produce NRLs corresponding to the mode spacing of euchromatin and heterochromatin. These adjusted sequence data were then combined to simulate different combinations of euchromatin and heterochromatin within the same region, with 182 + 187 simulating the euchromatin of female chrX. (**D**) Positions of nucleosome protection peak (NPP) calls from plot C plotted against their maximum WPS (mWPS). (**E**) WPS of adjusted sequence data combined to simulate chrX euchromatin of male sample pool versus a mixed-sex sample pool. (**F**) Median NPP spacing in heterochromatin and euchromatin. 'Heterochromatin' is composed of ChromHMM states 12 and 13 and 'Euchromatin' of states 1-2, 4-7, and 9-10 of the GM12878 cell line. Poised promoter (3) and weak transcription (11) have been excluded due to the ambiguity of these states.

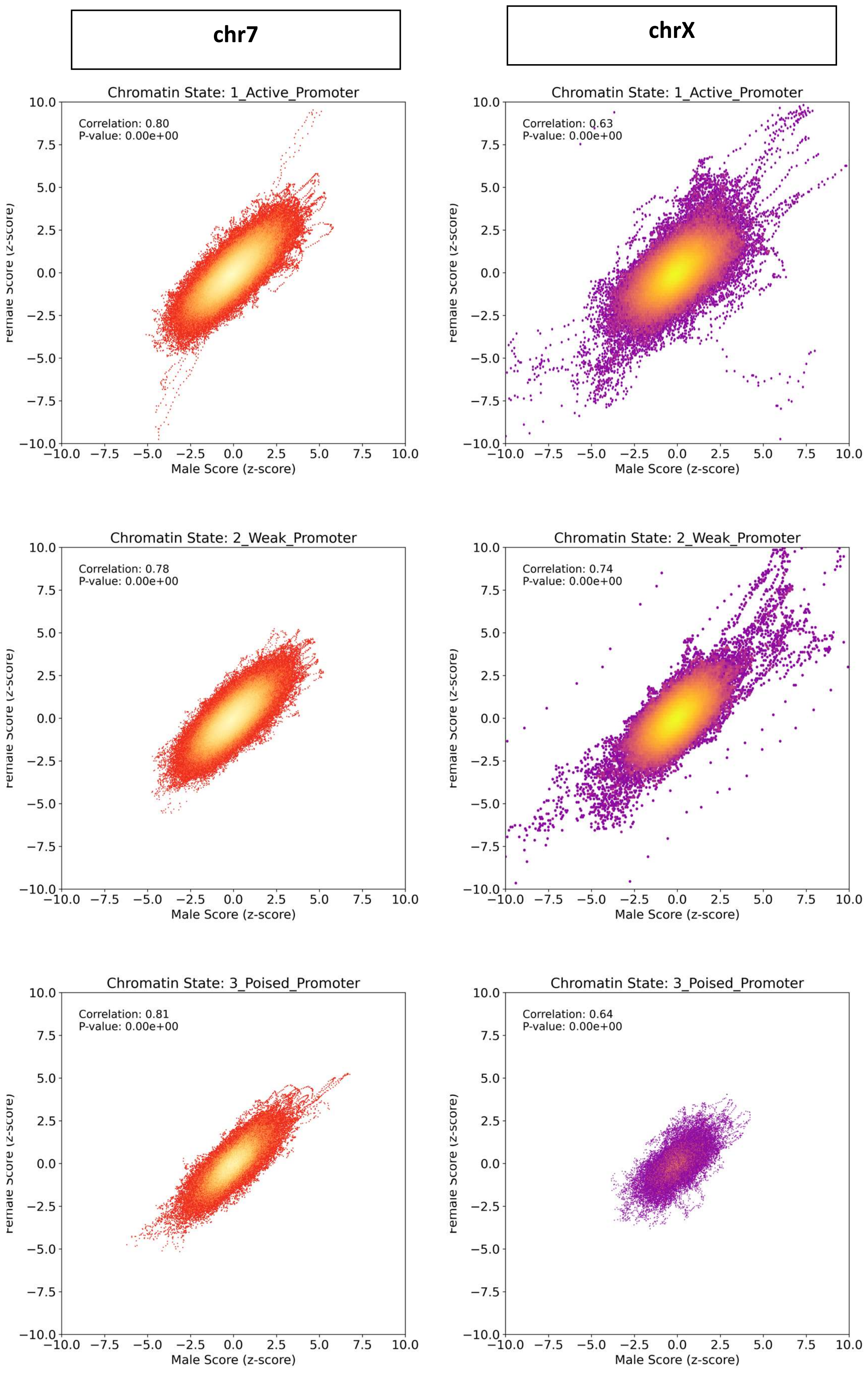

chr7
chrX
Chromatin State: 1_Active_Promoter
Correlation: 0.80
P-value: 0.00e+00
Chromatin State: 1_Active_Promoter
Correlation: 0.63
P-value: 0.00e+00
Chromatin State: 2_Weak_Promoter
Correlation: 0.78
P-value: 0.00e+00
Chromatin State: 2_Weak_Promoter
Correlation: 0.74
P-value: 0.00e+00
Chromatin State: 3_Poised_Promoter
Correlation: 0.81
P-value: 0.00e+00
Chromatin State: 3_Poised_Promoter
Correlation: 0.64
P-value: 0.00e+00
Male Score (z-score)

**chr7** | **chrX**

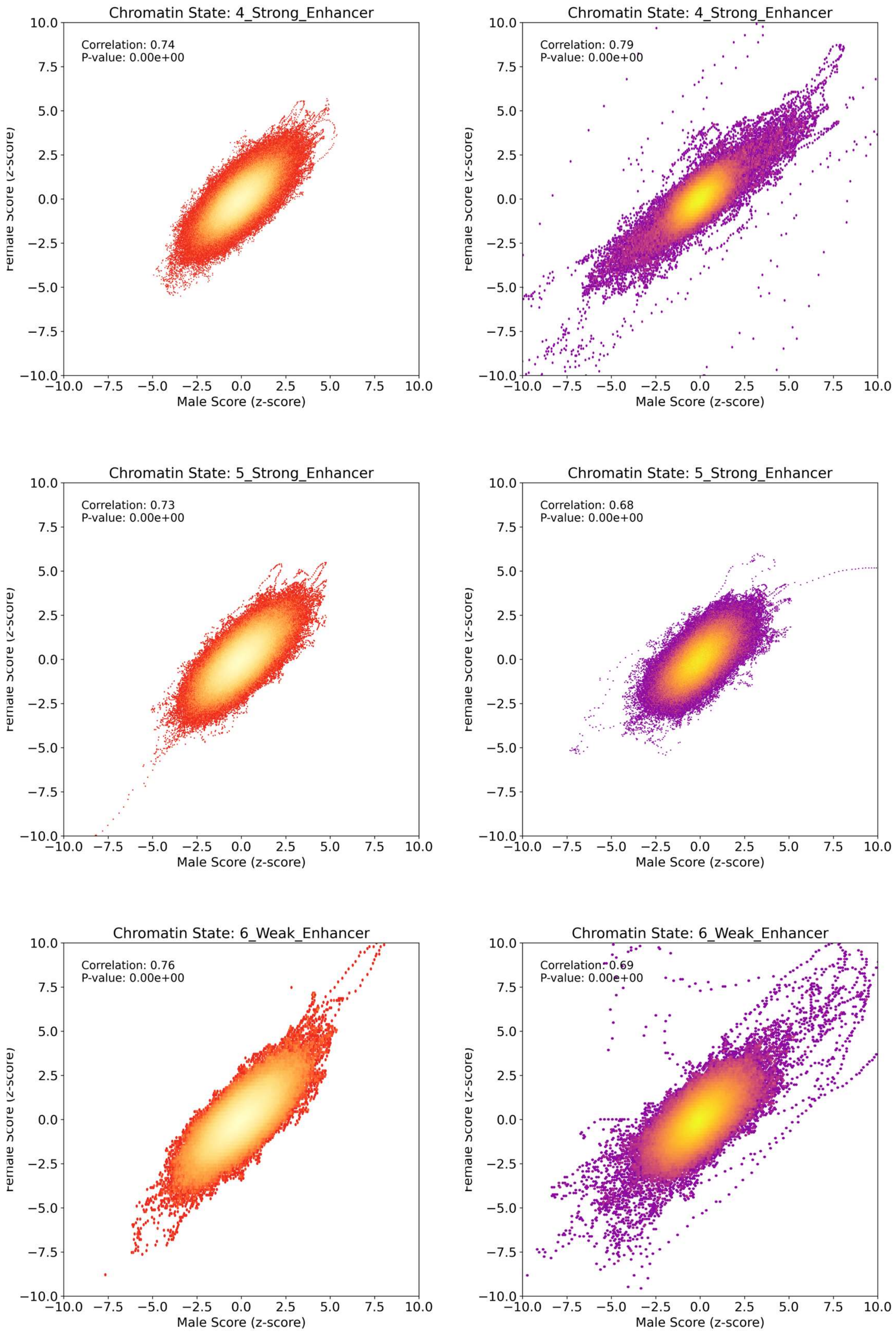
Chromatin State: 4_Strong_Enhancer
Correlation: 0.74
P-value: 0.00e+00
Chromatin State: 4_Strong_Enhancer
Correlation: 0.79
P-value: 0.00e+00
Chromatin State: 5_Strong_Enhancer
Correlation: 0.73
P-value: 0.00e+00
Chromatin State: 5_Strong_Enhancer
Correlation: 0.68
P-value: 0.00e+00
Chromatin State: 6_Weak_Enhancer
Correlation: 0.76
P-value: 0.00e+00
Chromatin State: 6_Weak_Enhancer
Correlation: 0.69
P-value: 0.00e+00
Male Score (z-score)

**chr7**

**chrX**

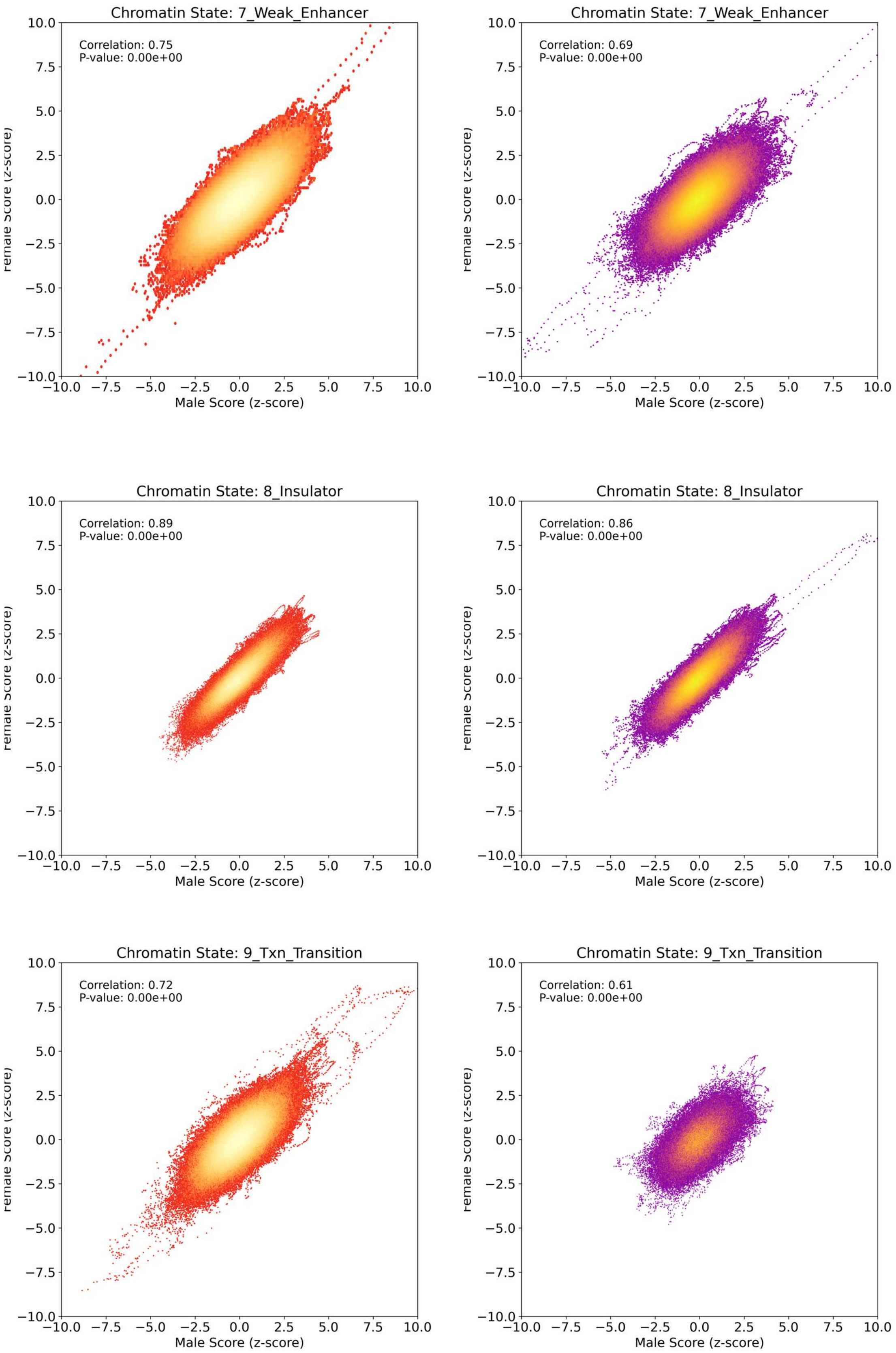
Chromatin State: 7_Weak_Enhancer
Correlation: 0.75
P-value: 0.00e+00
Chromatin State: 7_Weak_Enhancer
Correlation: 0.69
P-value: 0.00e+00
Chromatin State: 8_Insulator
Correlation: 0.89
P-value: 0.00e+00
Chromatin State: 8_Insulator
Correlation: 0.86
P-value: 0.00e+00
Chromatin State: 9_Txn_Transition
Correlation: 0.72
P-value: 0.00e+00
Chromatin State: 9_Txn_Transition
Correlation: 0.61
P-value: 0.00e+00
Male Score (z-score)
Female Score (z-score)

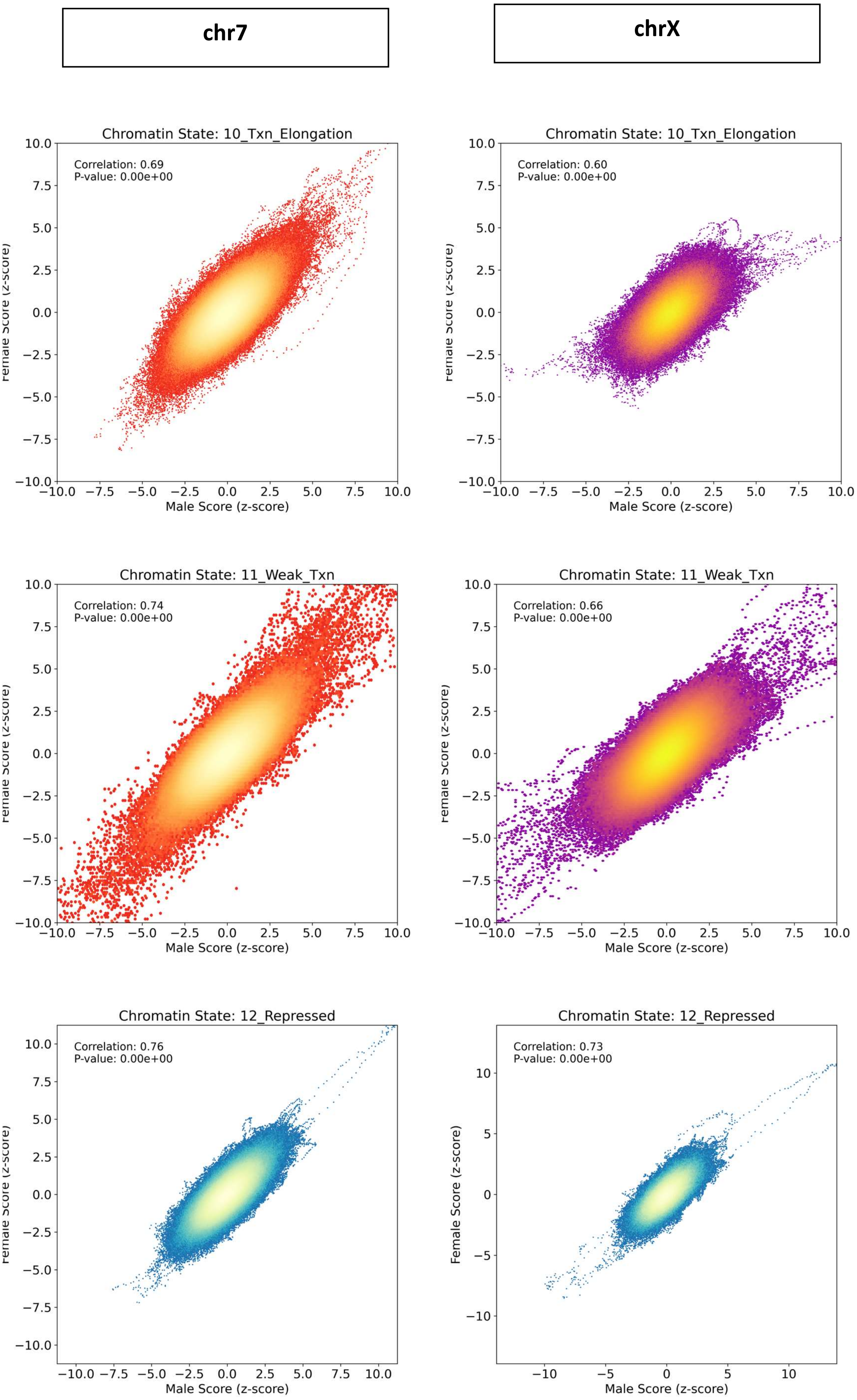
chr7
chrX
Chromatin State: 10_Txn_Elongation
Correlation: 0.69
P-value: 0.00e+00
Correlation: 0.60
P-value: 0.00e+00
Chromatin State: 11_Weak_Txn
Correlation: 0.74
P-value: 0.00e+00
Correlation: 0.66
P-value: 0.00e+00
Chromatin State: 12_Repressed
Correlation: 0.76
P-value: 0.00e+00
Correlation: 0.73
P-value: 0.00e+00
Male Score (z-score)
Female Score (z-score)

**chr7** **chrX**

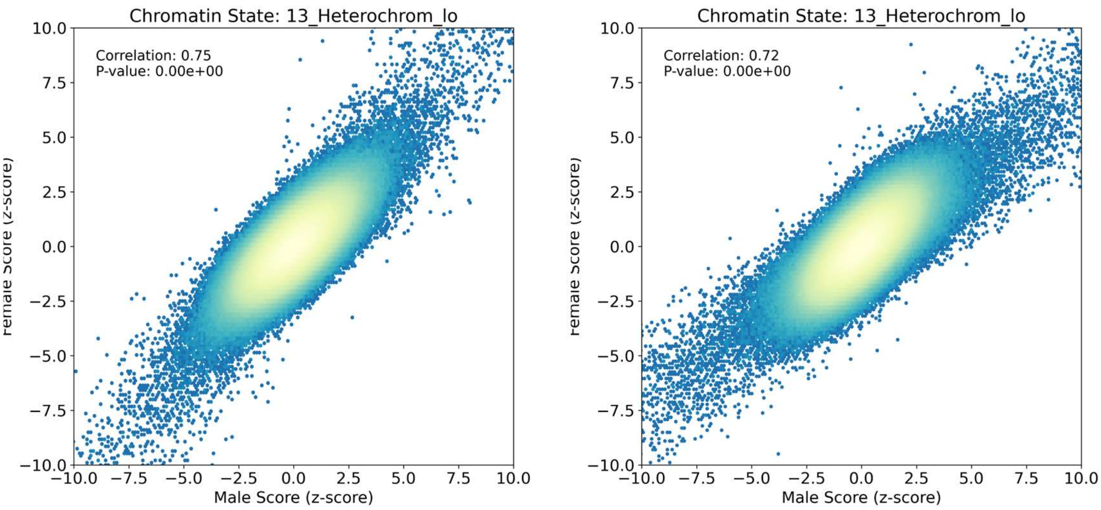


**Supplementary Figure S11. Correlation between male vs female nucleosome protection scores on chr7 versus chrX.** Correlations and P-values are those of Pearson correlation tests.

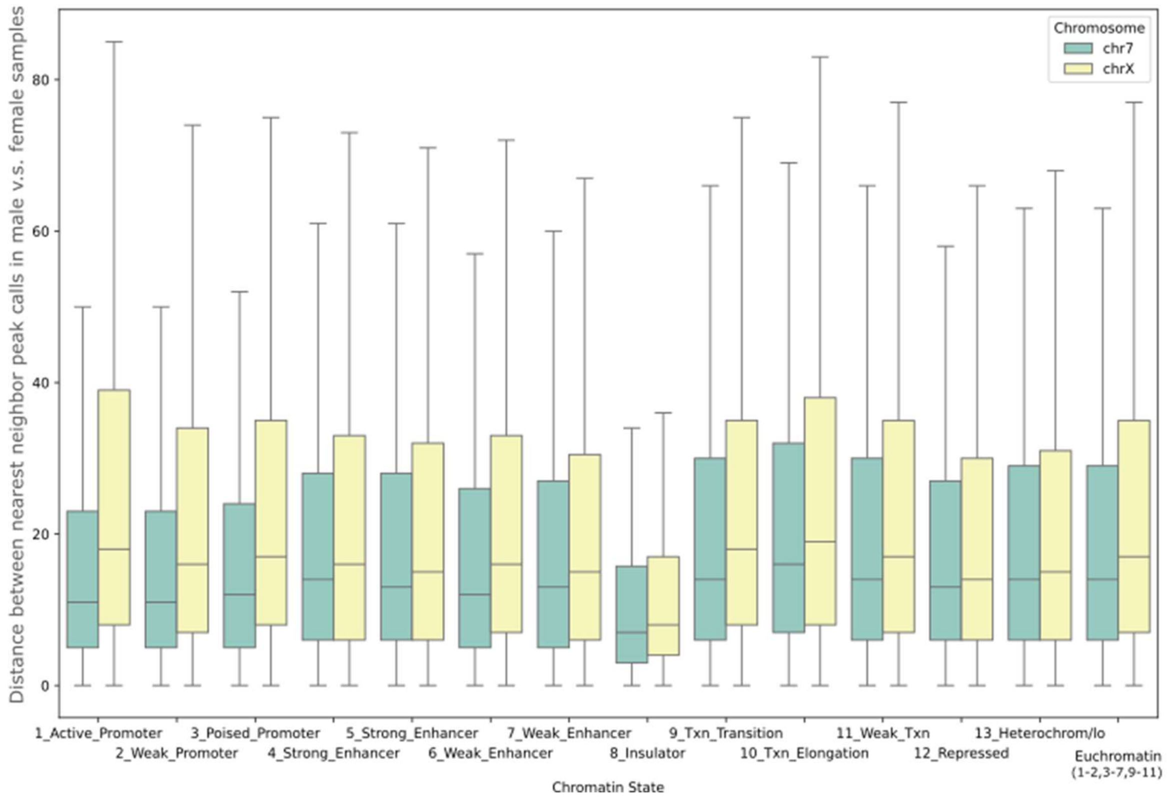


**Supplementary Figure S12.** Comparison of the absolute distances between nearest neighbor nucleosome peak calls on chromosome 7 and chromosome X in male and female sample pools. The analysis tests the hypothesis that euchromatin regions on chromosome X should exhibit greater distances between peaks compared to chromosome 7, reflecting nucleosome repositioning associated with X-inactivation.

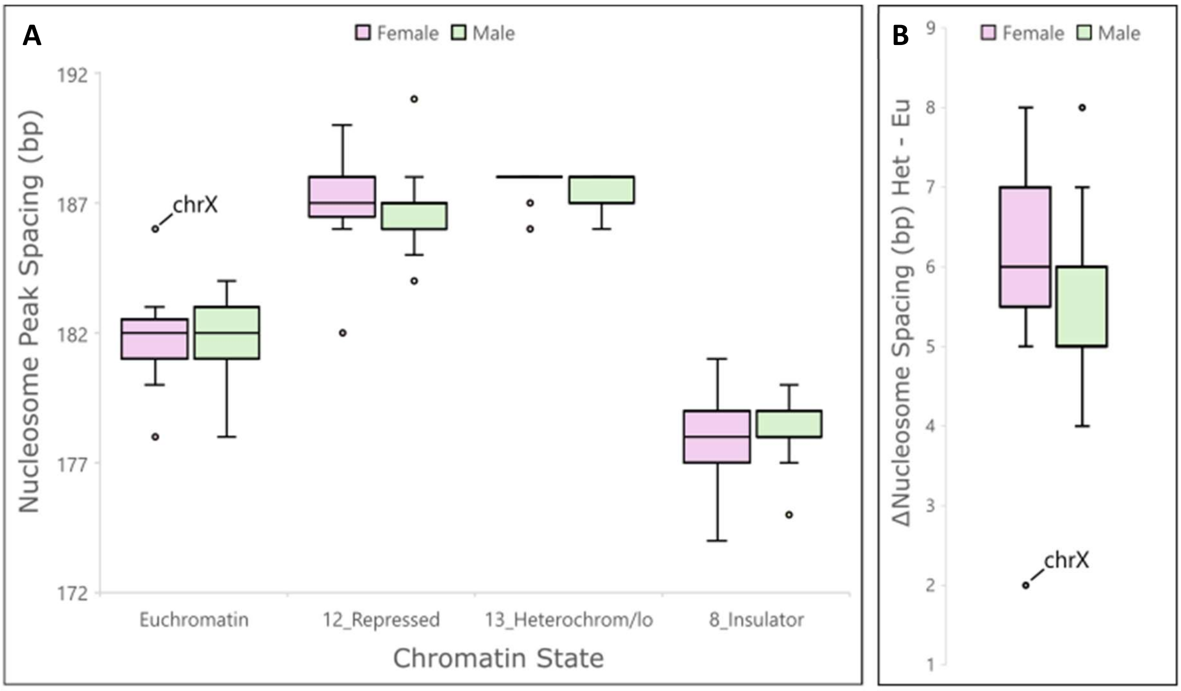


**Supplementary Figure S13.** Genome-wide analysis of differences in nucleosome spacing between heterochromatin and euchromatin regions, highlighting a reduction in female chrX for female cfDNAs, as predicted by our model. Samples are those of the CA01 pool. The depth of the male sample pools is around twice that of the female pool. To match, the reads from the male pool were down-sampled to 50%, except for chrX as the single copy of this chromosome already results in 50% lower coverage. Data was analyzed using our new nucleosome protection scoring (NPS) and peak calling model. (**A**) Mode interpeak distances for chromatin states on each chromosome calculated after Savitzky-Golay smoothing (window size = 35, polynomial of order = 3). (**B**) The difference in nucleosome spacing from panel A, highlighting the much lower difference in nucleosome spacing between heterochromatin and euchromatin on the chrX of female samples.
