## Supplementary material for "Unraveling cfDNA Fragmentation Patterns: Linking Chromatin Architecture to Cancer Diagnostic Sensitivity": Methods

### Patient samples

Plasma samples from 121 patients with breast or colorectal cancer were collected along with blood from 32 healthy controls for ddPCR analyses. All colorectal cancer patient cfDNA samples, and the majority of breast cancer patient cfDNA samples, were isolated using our in-house protocol based on the method described in Hufnagl et al. (2013)[1]. The three 4-plex ddPCR analyses presented in **Figure 2** were run on all 121 cancer patient cfDNA samples. However, 4 breast cancer and 2 colorectal cancer cfDNA samples were excluded from analysis after subsequent Agilent Bioanalyzer 2100 fragment size measurements run on all cfDNA samples did not show the ≈166 bp peak characteristic of apoptotically derived cfDNA. A further 13 colorectal cfDNA samples were excluded from analysis because they failed to produce adequate separation between 4-plex ddPCR amplitude/fluorophore clusters in at least one of the three 4-plexes. Furthermore, 14 colorectal cfDNA samples failed to amplify (i.e., showed little to no positive droplets across all three of the 4-plex ddPCR assays). The samples used in the somatic mutation analyses shown in **Figure S2** were a subset of those detailed above, selected based on the availability of remaining material after the initial ddPCR run. As for sequencing, the breast cancer and colorectal cancer pools contained 16 and 13 samples, respectively. The brain cancer pool contained cfDNA extracted from the plasma of 19 patients: 4 with astrocytomas, 11 with glioblastomas, and 4 with oligodendrogliomas. The plasma samples from breast cancer patients used in this study were recruited for a National Breast Cancer Foundation (NBCF) adjuvant clinical trial and plasma samples from healthy donors were received from the Red Cross Blood Service, Australia. The use of these human plasma samples was approved by the Bellberry Human Research Ethics Committee (application number 2015-12-817-A-6). The plasma samples from colorectal cancer patients were recruited for a study by the John Hunter Hospital and the use of these samples was approved by the Hunter New England Human Research Ethics Committee (reference number 11/04/20/4.03). Finally, the plasma samples from brain cancer patients were from The Wesley-St Andrew's Research Institute (WSRI) Tissue Bank, which has ethics approval from the UnitingCare Health Human Research Ethics Committee (HREC).

### In-house cfDNA extraction method

Plasma digests were conducted in DNA LoBind 5 ml tubes with 200 µl 1X Low TE, 900 µl plasma, 110 µl buffer (250 mM EDTA, 750 mM NaCl, 10 mM Tris), 110 µl 10% SDS, and 22 µl Proteinase K (20 mg/ml, NEB). Samples were then incubated for 2 hours in a 56°C water bath, mixing by hand every 30 minutes. Next, 1 volume (1342 µl) of pH 8.0 phenol/chloroform/isoamyl alcohol was added to tubes along with a large drop of silicone. Tubes were then vortexed for 3 seconds, revert mixed a few times, incubated for 5 minutes at room temperature, and then centrifuged at 13000 rpm for 15 minutes. The upper phase containing DNA was then removed and transferred into new LoBind 5 mL tubes. 2 µl of Glycoblue was added to tubes, along with 630 µl of 7.5M ammonia acetate and 2 volumes (2520 µl) of cold 100% ethanol and mixed well by inversion. Tubes were then incubated overnight at -20°C, then spun at room temperature for 30 minutes at 12000 rpm to pellet the DNA. DNA pellets were washed twice with 70% ethanol then air-dried and resuspended with 40 µl of 1X Low TE. The remaining breast cancer cfDNA samples were extracted using QIAamp Circulating Nucleic Acid Kit (Qiagen) using the manufacturer's recommended protocol, except for those where EconoSpin® All-In-One Mini Spin Columns (Epoch Life Sciences) were used rather than the columns supplied in the kit.

### PCR assay design

We initially utilized nucleosome positioning data from Snyder et al. (2016)[2] to inform placement of cfDNA breakpoint assays. This study calculated per-nucleotide windowed protection scores (WPS) by summing cfDNA fragments (120-180 bp) that fully overlapped a 120 bp window and subtracting those that partially overlapped. Scores were normalized to the median coverage within a 1000 bp window, and contiguous regions between 50 to 150 bp with scores elevated above this median were identified as nucleosome-protected peaks. This windowed protection scoring and peak calling method was used to predict nucleosome positions in cfDNA from over 50 healthy patients in the CH01 sample pool.

We analyzed this CH01 dataset to identify nucleosome protection sites that could be functionally validated using multiplexed droplet-digital PCR (ddPCR) assays. In selecting these sites the following criteria were enforced: (1) nucleosome sites selected for validation should exhibit a windowed protection score exceeding two standard deviations above the mean, which identified 117,069 sites; and (2) ddPCR validation of breakpoint prediction should examine multiple nucleosome protection peaks on different chromosomes, to minimize confounding variables that could be present at a single locus (e.g., copy-number variation, assay bias). To fine-map cfDNA breakpoints, these regions were processed through our PrimerSuite software (1) (www.primer-suite.com), generating sets of three ddPCR assays for each region that met strict parameters. For each region, one assay was centered on the nucleosome binding site, where the maximum amount of intact cfDNA molecules should be present; and two additional flanking assays were centered where the most DNA breakage should occur, 80bp upstream and 80bp downstream of the nucleosome binding site. To minimize variance when comparing data between different assays, all assays were designed to be exactly 100bp in size, with similar oligonucleotide annealing temperatures and GC content. From the sets produced, four regions (each with three assays) were selected using our PrimerDimer algorithm to produce optimal multiplexing that minimized primer-dimer interactions (PrimerROC REF). The final 12 assays were grouped into three multiplex ddPCR reactions, each containing four assays.

More specifically, assays were designed with the following parameters using a combined and modified in-house version of our previously published primer design, multiplexing and dimer prediction tools[3,4]: equal length amplicons of 100 bp centered either 0 or 80 bps from protection peak, minimal difference in primer length (20-24 bp) and the same $\Delta G°_{60}$ of -18.5 kcal/mol, and maximum probe length of 30 bp and the same $\Delta G°_{60}$ of -21 kcal/mol. The one exception to these parameters is the chr3 downstream breakpoint assay, which has a 30 bp reverse primer and is centered 88 bp from the protection peak. The primers for this assay were manually redesigned around the algorithmically designed probe after initial ddPCR tests showed poor performance.

Using the getFasta function from BEDtools[5] we extracted sequences surrounding desired nucleosome protection peaks of the CH01 sample. We then ran a modified version of our PrimerSuite[3] primer design algorithm on these sequences to produce all potential amplicons within specified length and $\Delta G°_{60}$ parameters centered on nucleosome peaks or 80 bp flanks. Each primer pair was then screened using Bowtie 2[6] version 2.3.0-legacy in paired-end mode for those forming unique single-mapped amplicons in a 1 kb window of the hg19 genome. The remaining assays were then run through a multiplex design Python script that incorporated our PrimerDimer[4] algorithm to create a multiplex that included assays targeting four different chromosomes and minimized potential for primer dimer formation.

### Droplet digital PCR

ddPCR analysis was performed on cfDNA samples with Bioanalyzer 2100 size peaks of ≈166 bp extracted from the blood plasma of 54 breast (**Figure S14**) and 34 colorectal (**Figure S15**) cancer patients. Nucleosome protection peak assays were algorithmically designed to target peaks with similarly high levels of nucleosome protection, with WPS between 148-153 in the CH01 map of Snyder et al. (2016)[2], two standard deviations above the mean WPS of 63.7 (SD = 41.4). For the cancer hotspot assays, we ran a 16-sample subset from breast cancer and a 14-sample subset from colorectal cancer patients. Each hotspot had one amplicon designed to center on the nearest nucleosome protection peak and another with the mutation somewhere between the two primers while attempting to maximize the distance of the amplicon from the nearest nucleosome peak, simulating a worst-case scenario for cfDNA assay design that fails to take nucleosome positioning into account. To minimize sample expenditure, assays were run on a Bio-Rad QX100 as ddPCR quadruplexes using the amplitude multiplex method developed by Dobnik et al. (2016)[7] in which probe concentrations are varied to alter the resulting levels of fluorescence amplitude, allowing for the detection of two targets per fluorescence channel. All amplicons were designed to be 100 bp, a length that falls well within the mode 166 bp of cfDNA fragments.

Samples were quantified using a QX100™ ddPCR™ System (Bio-Rad). Reactions were made to a 22 µL total volume and consisted of 1-6 µL of DNA, 2.36-7.36 µL $H_2O$, 11 µL 2x Supermix for Probes, 0.44 µL 10 u/µL HindIII, 2.2 µL 10x primer/probe 4-plex mix (5 µM per primer, 1 probe at 1.25 µM and 1 at 0.75 µM per FAM and HEX fluorophore). Droplets were generated using a QX200™ AutoDG™ Droplet Digital™ PCR System. Cycling conditions were 10 minutes at 95°C, followed by 45 cycles of 30 sec at 95°C and 2 min at 60°C, and ending with a 10 min 98°C enzyme deactivation step. Concentrations for each assay were determined in QuantaSoft™ Analysis Pro 1.0.596 by setting wells to Amplitude Multiplex mode and using the 2D Amplitude window.

### Creating gDNA samples with cfDNA-sized fragmentation

50 µL of blood pooled gDNA (50 ng/µL) was sonicated using a Covaris S2 Focus Ultrasonicator with the manufacturer's recommended settings for a 150 bp peak. Sonicated DNA was then run in a 2% E-Gel EX agarose (Invitrogen) on an E-gel Power Snap for 15 min. The gel was then cut into four sections around 150 bp of the sample using a UV lamp. Fractions were then purified using QIAquick Gel Extraction Kit (Qiagen) according to the manufacturer's protocol. The size distribution profiles were measured using a 2100 Bioanalyzer (Agilent Technologies) to confirm similar fragmentation profiles to cfDNA samples (**Figure S16**).

### cfDNA sequencing

Paired-end sequencing of cfDNA fragments was performed using a NextSeq 550 running either High-Output or Mid-Output kits. Individual cfDNA samples were first pooled into different cancer types. End repair was performed using NEB Ultra End Prep and adapter ligation by NEB Ultra II Ligation. Different IDT UMI adapters were ligated to each pooled sample. Amplification was then performed with Q5 Hot Start High-Fidelity 2 Master Mix and p5/p7 primers: 95°C for 5 min; 98°C for 30 sec; 7 cycles of 98°C for 15 sec, 57°C for 20 sec, and 72°C for 90 sec; and a final 72°C for 3 min step. Bead clean-up was performed using Sera-Mag SpeedBeads. Pool concentrations were then measured using a 2100 Bioanalyzer (Agilent Technologies) and equal concentrations of each pool were combined into a pooled sample for sequencing.

### Graphing and statistical analyses

Box plots and stacked bar plots were created using IBM SPSS Statistics 24. All other graphing was performed in Python 3 using the Matplotlib and Seaborn packges. Modal values presented are those of Savitzky-Golay smoothed peaks. Contiguous chromatin state/compartment regions were randomly shuffled using our chromatin_state_shuffler.py python script (https://github.com/traulab/BeadsOnASpring). Aligning to protein binding sites was performed using the center nucleotide of sequence motifs and pairing nucleosome protection peaks with the closest of these motifs. Distances of paired nucleosome protection peaks were plotted relative to this center. Nucleosome protection peaks were similarly paired with a randomly selected subset of 10,000 nucleosome protection peak. FIMO version 5.3.3 (https://meme-suite.org/meme/tools/fimo) was run on the hg19 genome using JASPER (http://jaspar.genereg.net) motifs MA0466.3 (CEBPB), MA0098.3 (ETS), MA0496.2 (MAFK) and MA0139.1 (CTCF) using a p-value threshold of 1.0E-4. ENCODE ChIP-seq peaks from wgEncodeRegTfbsClusteredV3.bed (https://hgdownload.cse.ucsc.edu/goldenpath/hg19/encodeDCC/wgEncodeRegTfbsClustered) were used for intersecting with FIMO binding sites to deduce occupied binding sites[8–10]. Transcription start sites are those of SwitchGear Genomics (https://switchgeargenomics.com) downloaded in BED format from the switchDbTss table on the UCSC Table Browser (https://genome.ucsc.edu/cgi-bin/hgTables)[11]. Windowed protection scoring for aggregate alignment to binding sites, transcription start sites and nucleosome protection peaks was performed using our in-house fragScorer.py python script. Aggregate WPS alignments were performed using our in-house scoreAligner.py python script. Nucleosome protection peaks flanking TF binding sites and transcription start sites were extracted using our in-house primerNuc.py python script (https://github.com/traulab/BeadsOnASpring).

### Nucleosome position maps

Nucleosome protection peak data includes the CH01 (GSE71378_CH01.bb), BH01 (GSE71378_BH01.bb), CA01 (GSE71378_CA01.bb), IH01 (GSE71378_IH01.bb), and IH02 (GSE71378_IH02.bb) samples from Synder et al. (2016)[2], and protection peak maps we generated for the IC17, IC37, CA01♀, CA01♂, BRA01, BRE01, COL01 and BBC01 samples. Each of our pools contained ~15 individual patient cfDNA samples of roughly equal concentrations (detailed above), and the whole genome cfDNA sequencing data from all three pools were also merged to create the BBC01 pooled sample data. We used the peak calling scripts and Linux pipeline from Snyder et al. (2016)[6] to create nucleosome maps for each of these samples. Nucleosome position maps were intersected with chromatin state data from Ernst et al. (2011)[12] (wgEncodeBroadHmmGm12878HMM) and chromatin compartment data from Rao et al. (2014)[13] (GSE63525_GM12878_subcompartments) using BEDtools[5]. These datasets are available online and can be located by visiting their respective publications. All these datasets use the hg19 genome assembly coordinates.

### DHS tissues of origin analysis

Using our in-house NPP_spacing_at_DHSs.py python script (https://github.com/traulab/BeadsOnASpring), nucleosome position maps of each sample were run against DNase-seq peak data for 125 cell lines (http://mirror.vcu.edu/vcu/encode/DNase-seq_Peaks/) to determine the bp distances between nucleosome protection peaks flanking the centers of each DHS peak.

### Nucleosome phasing simulation

The BAM file of BBC01 was parsed for pair-end reads within a 10,000 bp region of nucleosome array starting from chr12:34440000 (GRCh37). The median spacing of nucleosome protection peaks for this region was 197 bp. In order to produce nucleosome spacing that corresponded to genome-wide modes of heterochromatin (187 bp) and euchromatin (182 bp) we shifted the start and end positions of each sequenced fragment from the combined BBC01 sample pool towards an arbitrary point of origin, with progressively increasing distances the further fragment centers were from this origin. The following adjustments were performed on each pair-end read to shift its start and end positions:

distance = base pairs between the center point of read and the 34440000 origin

new_NRL = 182 or 187

ratio = new_NRL / 197

subtractor = distance – distance * ratio

new_start_position = read_start – subtractor

new_end_position = read_end – subtractor
